# Aneuploidies are an ancestral feature of trypanosomatids, and an ancient chromosome duplication is maintained in extant species

**DOI:** 10.1101/2023.06.26.546280

**Authors:** João Luís Reis-Cunha, Samuel Alexandre Pimenta Carvalho, Laila Viana Almeida, A Anderson Coqueiro-dos-Santos, Catarina De Almeida Marques, Jennifer Black, Jeziel Damasceno, Richard McCulloch, Daniella Castanheira Bartholomeu, Daniel Charlton Jeffares

## Abstract

**Background:** Aneuploidy is widely observed in both unicellular and multicellular eukaryotes, usually associated with adaptation to stress conditions. Chromosomal duplication stability is a tradeoff between the fitness cost of having unbalanced gene copies and the potential fitness gained from increased dosage of specific advantageous genes. Trypanosomatids, a family of protozoans which include species that cause neglected tropical diseases, are a relevant group to study aneuploidies, as their life cycle has several stressors that would benefit from the rapid adaptation provided by aneuploidies.

**Results:** By evaluating the data from 866 isolates covering 7 Trypanosomatids genera, we have observed that aneuploidies are present in the majority of clades, and have a reduced occurrence in a specific monophyletic clade that has undergone large genomic reorganisation and chromosomal fusions. We have also identified an ancient chromosomal duplication that was maintained across these parasite’s speciations, which has increased sequence diversity, unusual gene structure and expression regulation.

**Conclusion:** Aneuploidies are an important and ancestral feature in Trypanosomatids. Chromosomal duplication/loss is a constant event in these protozoans, common in *Leishmania* and *Leptomonas* and repressed in *T. brucei* and closely related protozoans. The number of chromosomes with extra copies in a given isolate is usually low, and only one chromosomal duplication was kept for long enough to greatly impact its nucleotide diversity. The thigh control of gene expression in this chromosome suggests that these parasites have adapted to mitigate the fitness cost of having this ancient chromosomal duplication.

## 1 Introduction

Aneuploidy, the presence of an unequal number of chromosomal copies, is widely found in nature, being observed in protozoans, yeast, plants and animals. Most aneuploidy studies have focused on cancer or yeast, where it has been linked with loss of cell cycle control, survival in stressful conditions, and adaptability. In yeast, the stability of chromosomal duplications was shown to be a tradeoff between the fitness cost of having unbalanced gene copies and the potential fitness gained from increased dosage of specific advantageous genes [1]. In this context, specific chromosomal duplications were linked to resistance against antifungal agents [2–5]. Aneuploidies are the most common genomic alteration in cancer, present in ∼90% of solid and ∼50% of hematopoietic tumours (Reviewed in [6]). Whilst aneuploid cancer cells can have reduced growth rates, particular combinations of chromosomal duplication can be advantageous under specific conditions [1,7–9], and promote chemotherapy resistance [10]. Human hepatocytes - cells which typically persist under constant stress - are largely aneuploid and polyploid, which provides resistance to injury and promotes liver regeneration [11]. Hence, aneuploidies have been shown to provide a rapid and often transient solution to stress conditions, emerging rapidly and more frequently than other genetic alterations, such as Single Nucleotide Polymorphism (SNPs), short insertion/deletions and gene copy number variants (CNV) [1,12].

Trypanosomatids, a family of monoxenous or heteroxenous obligatory single-celled parasites from the Kinetoplastid order (phylum Euglenozoa), are an interesting group to study aneuploidies. Their life cycle presents several stressors that could benefit from rapid and transient generation of aneuploidies, such as: having to evade the host immune response; adapt to a change of environment when moving between insect vectors and vertebrate hosts (for heteroxenous parasites); and undergoing the cellular reorganisation of changing life cycle forms, such as flagellated and near unflagellated promastigotes and amastigotes in *Leishmania,* or trypomastigotes and amastigotes in *T. cruzi.* These core features of the trypanosomatid life cycle likely require widespread changes in gene expression, which may be facilitated by selected chromosomal expansion patterns [13–15]. Moreover, trypanosomatid biology is unusual, and one reason for this aneuploidy-driven gene expression control is the near universal use of polycistronic transcription in these organisms, limiting their capacity to alter the transcription of individual genes via promoters, and instead increasing their reliance on mechanisms of gene expansion and contraction and post-transcriptional control mechanisms including mRNA decay or translational control [16,17]. Hence, chromosomal duplications could directly impact gene expression in these organisms by offering more templates to be transcribed [14,18,19].

The focus of aneuploidy study in Trypanosomatids is largely directed towards *Leishmania, Trypanosoma cruzi* and *T. brucei*, which causes devastating neglected tropical diseases (NTDs) in humans and animals, imposing severe health and economic burdens, especially in developing countries [20–22]. These three pathogens represent a relevant group to study aneuploidy, due to contrasting evidence for its presence across the different clades. Notably, aneuploidy is common amongst *Leishmania* species [14,19,23] and *T. cruzi* groups [24,25], but appears to be rare in *T. brucei* [26,27]. Aneuploidy has also been evaluated in other trypanosomatids [28–30], though our understanding of this phenomenon across the full Trypanosomatid clade is limited.

Aneuploidies have been widely studied in the *Leishmania* genus, where they have been documented in all species and arise potentially stochastically in individual cells within a population, in a process called “mosaic aneuploidy” [23,31–33]. Recent single-cell sequencing experiments have demonstrated the coexistence of a complex diversity of karyotypes within *Leishmania* clonal lines [34], which is in accordance with a model wherein different patterns of chromosomal expansions are constitutively generated in culture and likely selected for under more restrictive conditions. Notably, the pattern of chromosomal expansions in *Leishmania* varies when it was transferred from growth in culture, to the insect vector and to the mammalian host, further reinforcing the notion that different environments favour specific combinations of chromosomal duplications, thus conferring increased fitness [14]. Chromosomal duplications were also shown to directly impact gene expression in *Leishmania*, where chromosomes (chrs) with extra copies were highly expressed when compared to disomic counterparts, to the sole exception of *Leishmania* chromosome 31 (Leish Chr31). Leish Chr31 was found to be expressed as though being disomic, despite having ∼4 copies [14], suggesting that specific expression control mechanisms operate in this chromosome (chr). Both the basis for this expression control in Leish Chr31 and the persistent supernumerary status across all *Leishmania* species remain unknown. Aneuploidy, more specifically chr duplication followed by loss, was also associated with loss of heterozygosity (LoH) in *Leishmania*, in a process known as haplotype selection [19]. In this scenario, a duplication of a copy from a disomic chr with the haplotypes “A” and “B”, generates a trisomy (ABB), followed by the loss of one of the haplotypes, resulting in reversion to disomy (BB) and, consequently, LoH. This process enables the protozoan to eliminate potentially disadvantageous chromosomal copies and alleles without sexual recombination, and indicates that aneuploidy is a relevant process in *Leishmania*’s survival and evolution.

To date, analysis of aneuploidy beyond *Leishmania* has been focused on genome surveys in *T. brucei* strains and *T. cruzi* DTUs [24–26], suggesting the process is rare in the former and somewhat more common in the latter. Though some examples of LoH and trisomy have been documented in *T. brucei*, such as during adaptation of isolates to culture [27,35], the prevalence and impact has not been explored to the same depth as in *Leishmania*. More importantly, it is unclear when the capacity for aneuploidy arose in trypanosomatid evolution. In the present work, we evaluated aneuploidy prevalence across the trypanosomatid clade, examining isolates and clades with near chromosome-level-assembled reference genomes, and availability of Whole Genome Sequencing (WGS) read libraries. We evaluated not only the presence/absence of aneuploidies across trypanosomatids, but also measured the level of aneuploidy in each isolate, testing for chrs that are constantly expanded within and across clades. We found evidence for aneuploidy across the trypanosomatids, and detected an ancient, stable chromosomal duplication, which is both retained in almost all evaluated extant species, and for which gene expression is tightly controlled. This chr is expressed as if it was disomic, which could be a solution to the deleterious effects of having long-term full chromosomal extra copies.

## 2 Methods

### 2.1 Reference genomes, WGS read libraries pre-processing, mapping and chromosomal somy estimation

To estimate the occurrence of aneuploidies in Trypanosomatidae, we evaluated aneuploidy presence across 7 different genera, listed in (Table 1). To that end, we used a representative dataset of 866 whole genome sequencing (WGS) read libraries downloaded from the National Centre for Biotechnology Information (NCBI) Sequence Read Archive (SRA) using Fastq-dump (https://trace.ncbi.nlm.nih.gov/Traces/sra/sra.cgi?view=software). Only Illumina sequencing reads from publicly available dataset were used (Table 1 and Supplementary table 1). Each read library was filtered using fastp v2.10.7 [36], with the parameters: average Q20, minimal length 50 and removing the read extremities with base quality lower than Q25. Next, for each species/genera the reads were mapped to the reference genome listed in Table 1, using BWA-mem v.0.7.17 [37], retaining only reads with mapping quality 30 or higher using SAMtools v.1.10 [38]. The number of mapped reads was estimated using Samtools Flagstat [38]. The chromosomal somy for each sample was estimated using the median read depth coverage of single copy genes in a given chr with non-outlier coverage (Grubb’s tests, with P<0·05), normalised by the genome coverage, estimated by the coverage of all single copy genes in the genome, using Samtools depth [38]. The single copy genes were selected using OrthoFinder v. 2.5.4 [39,40], and the number of genes used in each dataset can be seen in Table 1. Only read libraries with genome coverage equal or greater than 10x were used in posterior analysis. Even though the read libraries were mapped in the full reference assemblies, only chrs/scaffolds larger than 75 kb were used in the Chr Copy Number (CCN) estimations. The full list of SRAs, library read counts, percentage of mapping and CCN estimations can be seen in Supplementary Table 1. The heatmaps and dot plots representing the CCN variation (CCNV) were generated using R using the libraries heatmap2 (https://cran.r-project.org/web/packages/gplots/index.html) and ggplot2 [41].

**Table 1:**
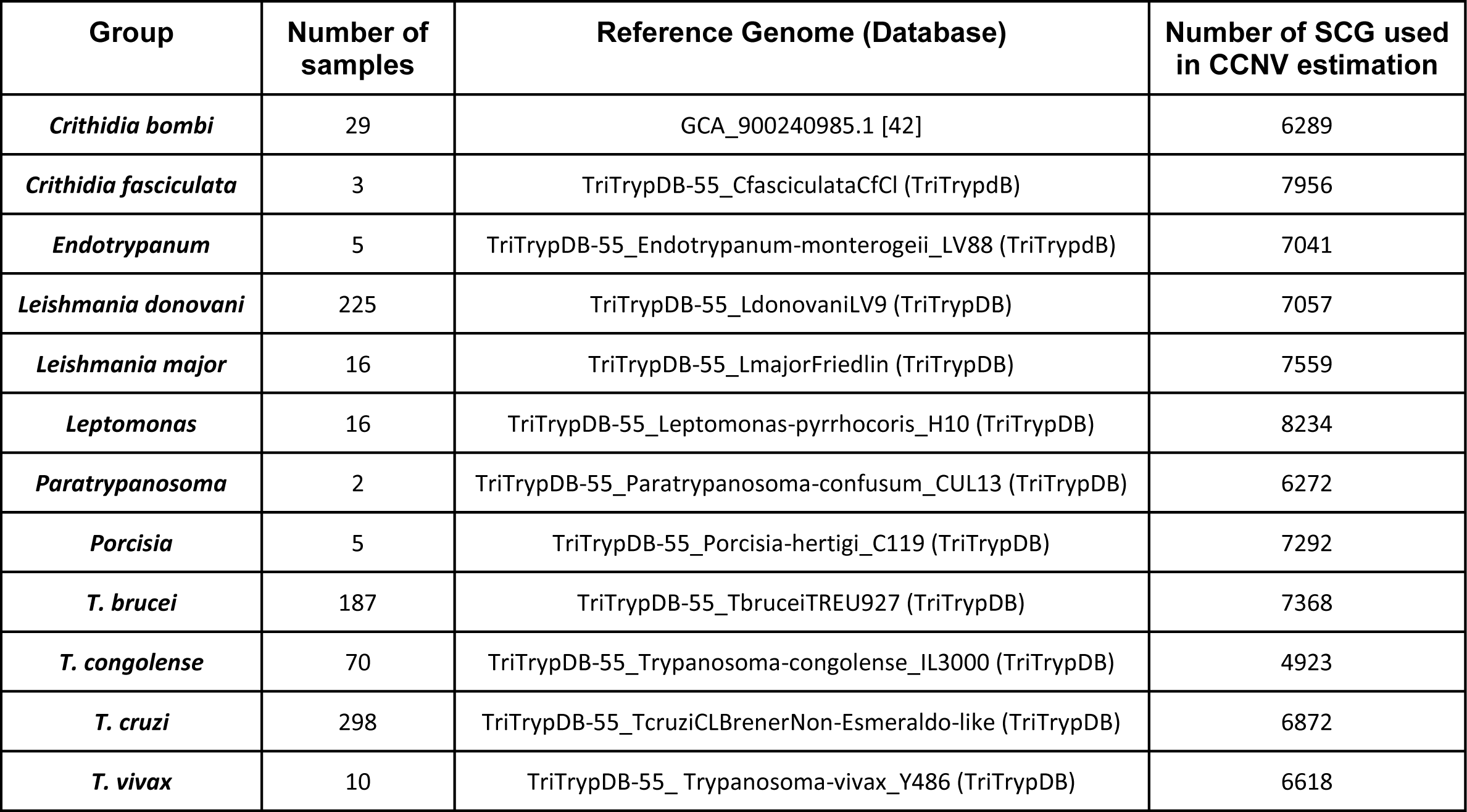
Read library collections and reference genomes. SCG: Single Copy Genes.

The intra-isolate assessment of aneuploidy and chr duplication was measured using two metrics: the isolate mean chr copy number, representing within-isolate genome expansion (**IGE**), and the isolate standard deviation of chromosomal copies, which represents within-isolate aneuploidy level (**IAL**). For IGE, the somy of each chromosome in an isolate is summed, and divided by the number of chromosomes. The IAL corresponds to the standard deviation of chromosomal copies within an isolate. The population variation of each specific chromosome copy was also measured with two metrics: chr mean copy number across isolates, representing chr duplication occurrence (**CDO**) and chr copies standard deviation across isolates, which represents chr somy variation (**CSV**). For CDO the mean copy number of a given chromosome was estimated by summing the values for the same chromosome in each isolate, and dividing by the number of isolates, while CSV represents the standard deviation of the copy of the same chromosome across isolates. Mean and standard deviation of chromosomal copies within isolates and within chrs were estimated using R.

### 2.2 Phylogenetic and gene synteny analysis

The Trypanosomatids phylogeny was estimated by maximum likelihood, using the single copy genes that were shared in 30 different Kinetoplastida groups/isolates (Supplementary Table 2). In short, the predicted proteome of the 30 groups were filtered to only keep annotated proteins that start with a methionine, end with a stop codon and had more than 100 amino acids. Then, Orthofinder v.2.5.4 [39,40] was used to identify the 216 single copy gene clusters that were present in all evaluated samples and align their sequences using MAFFT, 7.470 [43]. Parallel Alignment and back-Translation (ParaAT) [44] was used to convert the protein alignment in nucleotide alignments and TrimAI [45] was used to polish the alignments and remove gap regions, resulting in a final alignment containing 122,937 positions. ModelTest-NG [46] was used to select the best nucleotide substitution model for each gene partition and catfasta2phyml (https://github.com/nylander/catfasta2phyml) was used to concatenate the alignment of all genes. Finally, the maximum likelihood phylogeny was estimated using RAxML-NG [47], using the best model for each gene interval in the concatenated alignment and 1,000 bootstrap replicates. The tree representation was generated in R using ggtree [48]. The colorstrip representing the presence/absence of aneuploidies was based on data from this work and from the literature.

### 2.3 Gene sharing and gene organisation in *Leishmania* Chr31 and related sequences

The orthologous genes between *Leishmania major* chr 31 and chrs/scaffolds in each of the other 10 species/groups listed in Table 1 were identified using OrthoFinder v.2.5.4 [39,40]. The chrs/scaffolds with the highest numbers of shared ortholog genes with *Leishmania* Chr31 were identified, and used in further analysis. The bar charts representing the number of ortholog genes in each chr/scaffold for each species/group were generated in R. The gene coordinates and orientation of each ortholog gene as well as the chr/scaffold size was recovered from the General File Format (GFF) files, obtained from NCBI, TriTrypDB v55, or as supplementary data in the publication [42], in accordance with Table 1. The synteny circular plots were generated with Circa (https://omgenomics.com/circa/). The chromosomal gene disposition plots were generated in R using the library genoPlotR [49].

### 2.4 Nucleotide diversity (π) estimation

The Nucleotide diversity (π) was estimated in three representative datasets that contained more than 10 isolates and evidence of consistent aneuploidy: *L. donovani* EA_HIV set (113 samples); *C. bombi* (29 samples); and *Leptomonas* (16 samples). *T. cruzi* was not used due to its large content of repetitive multigene families [50–52]. Read filtering and mapping was done as described in “2.1-WGS read libraries pre processing, mapping and chromosomal somy estimation”. Read groups were assigned for the filtered mapped read libraries, using PicardTools v.2.21.6 AddOrReplaceReadGroups (https://github.com/broadinstitute/picard). SNPs and indels were called using the Genome Analysis Toolkit (GATK) v.4.1.0.0 HaplotypeCaller and the discovery genotyping mode of Freebayes v. 1.3.5 (https://github.com/ekg/freebayes), with a minimum alternative allele read count of 5. Only SNP/Indel positions that were identified by both callers were kept. For each dataset, the single-sample VCFs were merged with VCFtools v.0.1.16 and regenotyped using Freebayes. Next, the VCF file was filtered using BCFtools v.1.12 [53], to select only biallelic SNPs, with call quality above 200, coverage greater than a quarter of the genome coverage, and lower than four times the genome coverage, with mapping quality 40 or higher and properly paired reads (-m2 -M2 -i ’ TYPE=“snp” & QUAL > 200 & INFO/DP > Cov/4 & INFO/DP < Cov*4 & INFO/MQM >40 & INFO/MQMR >40 & INFO/PAIRED > 0.9). The Nucleotide diversity (π) was estimated using two approaches: 1-Using 10 kb sliding windows with VCFtools v.0.1.16 --window-pi [54], correcting for reporting windows with “0” SNPs using a custom R script; 2-Estimating π in each gene using custom R scripts. The comparisons of the π 10 kb in windows in duplicated chrs in each dataset was performed using Kruskal-Wallis and the Dunn’s post-test with Bonferroni correction for multiple tests, implemented in the dunnTest function from the FSA (https://www.rdocumentation.org/packages/FSA/versions/0.9.4) R package. The plots were generated in R.

### 2.5 Gene expression control in *Leishmania* Chr31

To evaluate the gene expression regulation in *L. donovani* Chr31, a dataset containing six parasite clones with matched WGS and RNAseq described in [19] Supplementary Table 3 was used. Both the WGS and RNAseq read libraries were downloaded, polished and mapped to the *L. donovani* LV9 reference genome as described in the section “2.1-*WGS read libraries pre processing, mapping and chromosomal somy estimation*”. To assess if the overall gene expression in a chr was impacted by its copy number (confirming the results shown in [19]), the chromosomal coverage obtained using the WGS reads and the RNAseq reads was estimated with Samtools v.1.10 Coverage [38], and normalised by the genome coverage, estimated as the mean coverage of all chrs. The correlation between the CCN and expression was estimated using the cor.test function in R.

Next, to assess how *L. donovani* regulates the gene expression of Leish Chr31, we compared the Alternate Allele Read Depth (AARD) in heterozygous positions in both WGS and RNAseq libraries in the same clone. There are two main scenarios: 1-All Leish Chr31 chromosomal copies are downregulated similarly; 2-Two chromosomal copies are silenced, and two are expressed. In unbalanced heterozygous positions, for example in a SNP position in a tetrasomic chr with three chrs with an alternate allele “C” and one chr with a reference allele “A”, the WGS AARD would be “0.75”. In the scenario “1”, the RNAseq AARD would match the WGS AARD (i.e. 0.75), as all copies would be silenced similarly. On the other hand in scenario “2”, if chrs “AC” were expressed and “CC” were silenced, the AARD would be lower (0.5); or if chrs “AC” were silenced and “CC” were expressed it would be higher (1.0). Hence, a positive correlation between WGS and RNAseq in heterozygous SNP regions would be an indication of the scenario “1”; while a lack of correlation would be indicative of scenario “2”. The SNP call and filtering was performed as described in the section “2.4-*Nucleotide diversity (π) estimation*”. To estimate the AARD, the multi-sample VCF was imported in R and converted in a table using the library vcfR [55], and processed with custom R scripts. The correlation between the AARD in the WGS and RNAseq datasets was estimated using cor.test, in R.

To evaluate if the lower expression in Leish Chr31 could be caused by an differential codon usage, the “Measure Independent of Length and Composition” (MILC) value was estimated for each chr, using the R library coRdon (https://bioconductor.org/packages/release/bioc/html/coRdon.html) and the *L. donovan*i LV9 annotated CDS downloaded from TriTrypDB v55..

### 2.6 Comparison of disomic or tetrasomic SNPs callers in chr31 Minor Allele Frequency (MAF), nucleotide diversity and Synonym/Non-synonym mutation estimations

To evaluate the impact somy aware calling SNPs in Leish Chr31, we performed the SNP calling in Leish Chr31 assuming disomy or tetrasomy, as described previously (sections “*WGS read libraries pre processing, mapping and chromosomal somy estimation*” and “*Nucleotide diversity (π) estimation*”) using the *L. donovani* EA_HIV dataset and changing the ploidy value to 4. The nucleotide diversity and MAF were estimated based on the mean number of differences between the haplotypes in the population, using custom scripts in R optimised to disomic or tetrasomic data. The number, location and potential biological impact of synonym and non-synonym SNPs was estimated with SNPeff v.5.0d [56].

### 2.7 Evaluation of gene functions in consistently duplicated chrs

The evaluation of biological relevant functions in chrs that were consistently with extra copies was performed by manual evaluation of gene annotations and by *Gene Set Enrichment Analysis by ontology* (GESEA). This analysis was performed with the representative well annotated *L. donovani* (BPK282A1), *T. brucei* (TREU927) and *T. cruzi* (CL Brener) genomes. The whole genome CDSs from these three strains were downloaded from TriTrypDB v55, translated with SeqKit v.0.12.0 and the ortholog groups between these samples was estimated with OrthoFinder v.2.5.4 [39,40]. The GESEA analysis was performed using the TopGO package, using the elim-fisher and multiple-test correction by Benjamini-Hochberg, with GO Annotation Files (GAF) downloaded from TriTrypDB v55. Three comparisons were made: **1-Shared in all species:** Genes in orthogroups with members from *L. donovani* Chr31, *T. cruzi* Chr31 and in the interval 1.0 to 1.5 Mb in *T. brucei* chr4 and 2.0 to 2.5 Mb chr8 compared with all genes in the three species (All TriTryps). In this comparison only GOs with at least three annotated genes in the test set were evaluated; **2-Shared between two species:** Genes that were present in these chrs only in two species compared with all genes (sets: Leishmania_Tbrucei, Leishmania_Tcruzi, Tbrucei_Tcruzi). In this comparison, only GOs with at least two annotated genes in the test set were evaluated; **3-Exclusive genes:** Genes from the aforementioned chrs that are exclusive for each species (*Leishmania, T. brucei, T. cruzi*).

## 3. Results

### 3.1 Aneuploidy is an ancestral characteristic in Trypanosomatids, but is rare in the *T. brucei* clade

To evaluate the occurrence of aneuploidy across Trypanosomatids, we determined CCNV in a representative dataset containing 866 isolates from 7 genera: *Crithidia, Endotrypanum, Leishmania, Leptomonas, Paratrypanosoma, Porcisia* and *Trypanosoma* (**Table 1**). This gave us an unprecedented resolution to comprehend not only the presence/absence of chromosomal expansions, but also to identify their patterns and sharing within and across clades. We found clear evidence for consistent aneuploidies in all evaluated taxa, to the exception of the closely related *Trypanosoma* species: *T. brucei, T. congolense* and *T. vivax*. Heatmaps of individual chr copy number in *Leptomonas* (aneuploid) and in *T. vivax* (euploid) can be seen in Figure 1A and Figure 1B, respectively, and for all the other clades in Supplementary Figure 1.

**Figure 1.**
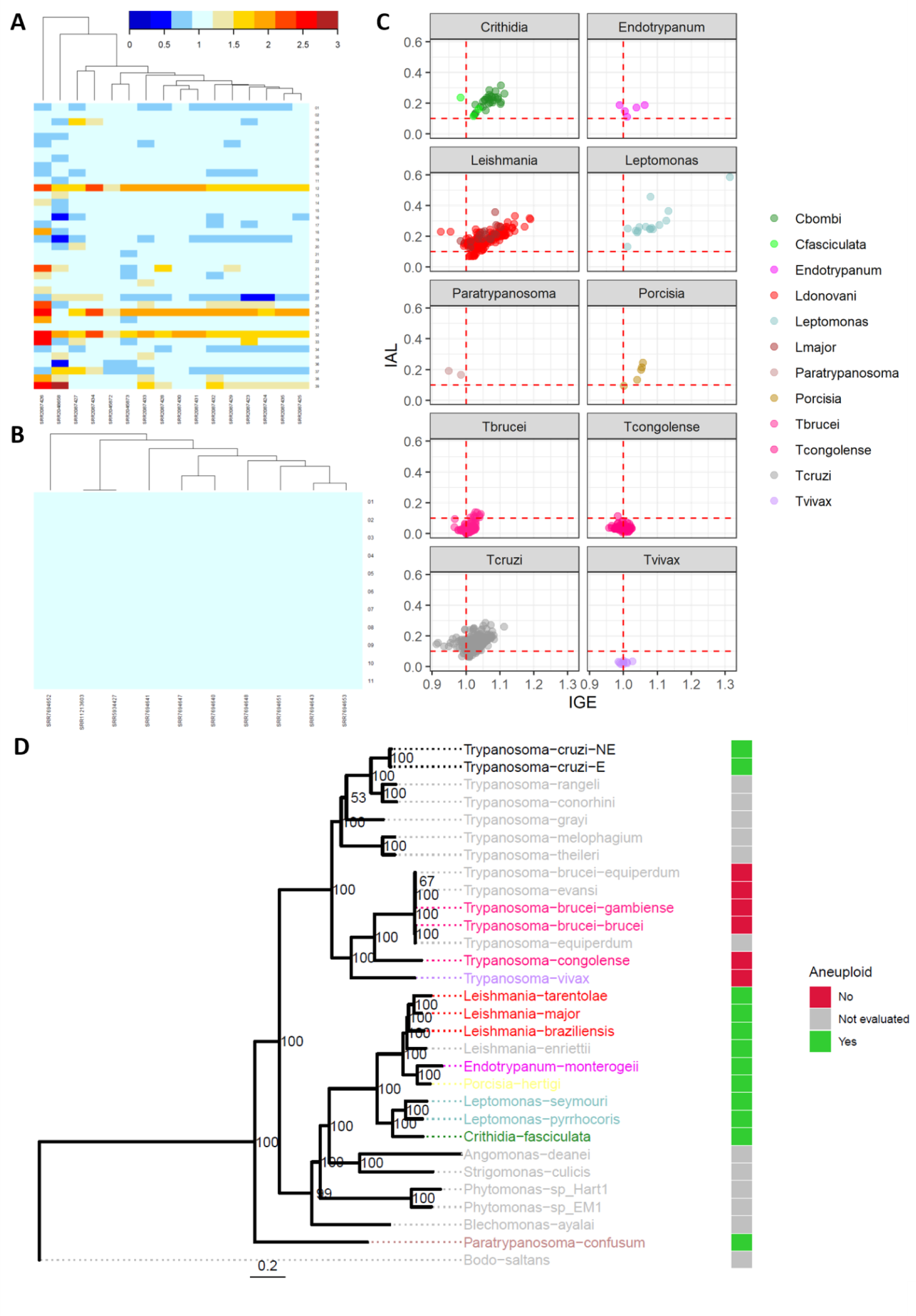
Trypanosomatid tolerances of aneuploidies. Somy estimation for all chrs, exemplified by the aneuploid *Leptomonas* **(A)** and the euploid *T. vivax* **(B)**. Heatmaps for all isolates and all species can be seen in the Supplementary Figure 1. In the heatmaps, each line corresponds to a chr, and each column to an isolate. The colour range represents the chr somy, ranging from blue (low copies -less than 1 copy for each haploid genome copy), light-blue (expected copies for a euploid isolate) to red (polysomic). **(C)** Scatterplot showing the isolate genome expansion IGE (X axis) and isolate aneuploidy level IAL (Y axis) of the chromosomal somy in each isolate. Each dot corresponds to a given isolate. The vertical red dashed line in the X value of “1’’ represents the expected copy number for euploid samples, with one copy for each haploid genome copy. The horizontal red dashed line corresponds to the IAL of 0.1, representing isolates with large CCNV variability. **(D)** Trypanosomatids maximum likelihood phylogeny, highlighting the clades that are aneuploid/mostly euploid and not evaluated. Node numbers in the tree correspond to the percentage of bootstrap support. Clades that were evaluated in this work are coloured, while non-evaluated clades are in grey. The colour strip represents the combined knowledge about presence/absence of aneuploidies in a given clade.

Initially, to assess the level of aneuploidy that was observed in a given isolate, each isolate “aneuploidy content” was measured based on two parameters: **1-** the Isolate Mean Chr Copy Number, representing within-isolate genome expansion (**IGE**), where high values occur when several chrs are duplicated in a single isolate; and **2-** the Isolate Standard Deviation of Chromosomal Copies, which represents within-isolate aneuploidy level (**IAL**), where high values occurs when there is large intra-isolate variation in chr copies (Figure 1 C). High values of IGE indicate that several chrs have extra copies, whereas high values of IAL indicate large *differences* between chr copies within an isolate. All these values were measured relative to haploid copies, where “1” means one copy per haploid genome copy. There was a large variation of aneuploidy levels in a given isolate between and within clades (Figure 1C and Supplementary Table 4). The top three highest IGEs were observed in *Leptomonas* (isolate SRR2087426: 1.313) and *Leishmania* (isolates ERR205781 and ERR205790 with, respectively, 1.189 and 1.187). The low IGE observed in the majority of isolates (min=0.912, mean=1.02, max=1.31) for all groups suggests that variations in chr copy number from the basal ploidy of trypanosomatids occurs in just a few chrs in a given isolate. This restriction of overall chr expansion may be caused by the high fitness cost of maintaining chromosomal-wide gene duplications, as seen in yeast [1].

When the IAL was evaluated, the highest values were once again observed in *Leptomonas* and *Leishmania* (Supplementary Table 4). Some isolates had strikingly high values of both IGE and IAL, such as the highly aneuploid samples “ERR205781” (IGE 1.189 and IAL 0.311) and “ERR205790” (IGE 1.187 and IAL 0.318) from *L. donovani*, and the *Leptomonas* isolate “SRR2087426” (IGE 1.31 and IAL 0.585). These highly aneuploid isolates contrast with others from the same clades that are mainly euploid, such as the *L. donovani* isolate “ERR4678144” (IGE 1.009 and IAL 0.071) and the *Leptomonas* “SRR2045872” sample (IGE 1.011 and IAL 0.132). This suggests that aneuploidy is tolerated, but not a required feature in these protozoans. Some other clades, such as *T. congolense* and *T. vivax* were mainly euploid, where their most variable isolates were, respectively, “ERR1993049” (IGE 1.024 and IAL 0.033) and “SRR7694652” (IGE 1.025 and IAL 0.035). In fact, 80-100% of the *Crithidia, Endotrypanum, Leishmania, Leptomonas, Paratrypanosoma, Porcisia* and *T. cruzi* isolates had an IAL higher than 0.10, while only 0-5% of the *T. brucei, T. congolense* and *T. vivax* isolates were above this cutoff (Table 2). This suggests that there is a lower tolerance of aneuploidy in *T. brucei* and closely related protozoans.

**Table 2:** Isolates with IAL higher than 0.1.

| Isolates | Number of Isolates | Isolates with IAL ≥ 0.1 | Percentage |
| --- | --- | --- | --- |
| *Crithidia bombi* | 29 | 29 | 100 |
| *Crithidia fasciculata* | 3 | 3 | 100 |
| *Endotrypanum* | 5 | 5 | 100 |
| *Leishmania donovani* | 225 | 214 | 95.11 |
| *Leishmania major* | 16 | 16 | 100 |
| *Leptomonas* | 16 | 16 | 100 |
| *Paratrypanosoma* | 2 | 2 | 100 |
| *Porcisia* | 5 | 4 | 80 |
| *T. cruzi* | 298 | 283 | 94.96 |
| *T. brucei* | 187 | 9 | 4.81 |
| *T. congolense* | 70 | 1 | 1.42 |
| *T. vivax* | 10 | 0 | 0 |

Finally, when the total evidence for presence/absence of aneuploidy was evaluated in context of a trypanosomatid phylogenetic tree (Figure 1 D), the presence of CCNV in distant clades (as in *T. cruzi* and *Leishmania*), and in the basal *P. confusum* are strong indicators that aneuploidy is an ancestral characteristic of trypanosomatids, which has later evolved to become largely absent in the *T. brucei* and closely related protozoans clades, that are monophyletic. In this scenario, the reorganisation of *T. brucei* and related protozoan genomes into a reduced number of larger chrs [57] could increase the fitness cost of aneuploidies due to the increased the number of gene products with unbalanced proportions that are generated by each chromosomal duplication, restricting its occurrence in this clade.

### 3.2 Consistent chromosomal expansions are observed in the majority of Trypanosomatid clades

It is well-established that chr 31 is consistently supernumerary in *Leishmania* species (chr 30 in *L. mexicana* complex, due to reordering in chromosome numbers due to 8-29 and 20-36 chromosomal fusions) [23,32,58]. As aneuploidy is frequent in the majority of Trypanosomatid clades, we went on to investigate if there were chrs that consistently vary in copy number and if there were chrs that are consistently expanded in each clade. To that end, two metrics were used to characterise the ploidy of individual chromosomes: **1-** chr mean copy number across isolates, representing chr duplication occurrence (**CDO**) in the population; and **2-** chr copies standard deviation across isolates, which represents chr somy variation (**CSV**): how variable the presence of a given chromosomal copy gain/loss is across the population (Figure 2A, Supplementary Table 5).

**Figure 2.**
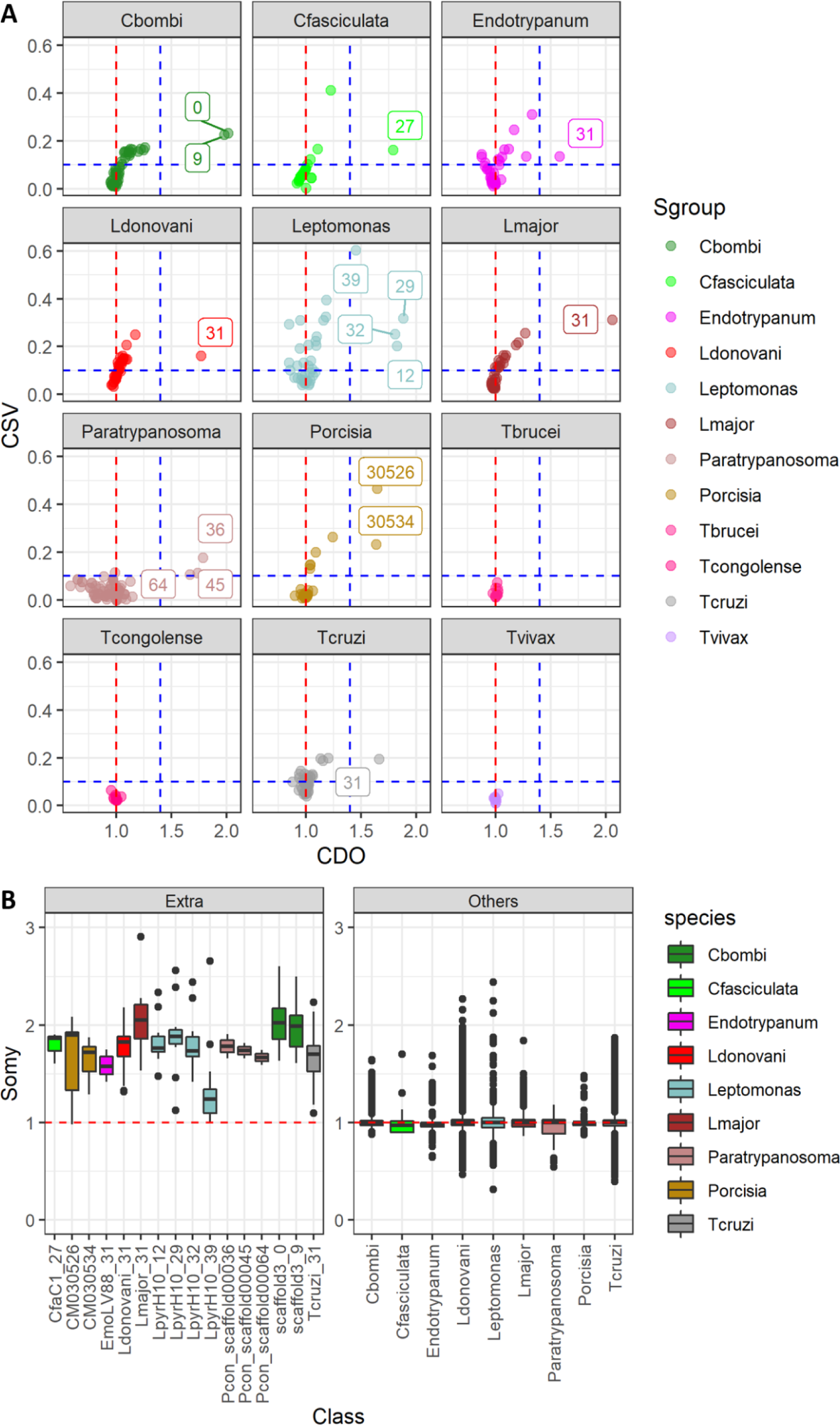
Aneuploidy varies between chrs, with certain chromosomes constitutively expanded in each clade. **(A)** Scatterplot showing the CDO (X axis) and CSV (Y axis) of the chromosomal somy in each chr across each population, representing the somy variability for each chr. Each dot corresponds to a different chr. Chrs with CSV > 0.1 and CDO > 1.4 are identified by their numbers/names, representing the chrs with highest copy number variability. The horizontal red dotted line represents the expected CDO value for euploid chrs. The horizontal and vertical blue dotted line corresponds respectively to CSV 0.1 and CDO 1.4. **B)** Somy of chrs that were consistently expanded in each clade. The panel “extra”, corresponds to the chromosomes that were consistently expanded in each clade, and the panel “others” corresponds to the somy of all chrs that had a population median somy <=1.4. The horizontal red dotted line represents the expected value for euploid chrs.

We calculated these metrics for each chr, for each species. Some chrs had both high CDO and CSV, such as *C. bombi* chrs 0 and 9, *L. donovani* chr 31 and *Leptomonas* chrs 12, 29 and 32, suggesting that there is some variation in their copy number, but that they had a high copy number baseline. Others, such as *C. bombi* chrs/scaffolds 1, 91 and 134, and *L. donovani* chr 08 and and 23, had a high CSV and a low CDO, suggesting that although aneuploid in some isolates, their expansion is not consistent in all isolates. There were also some chrs that were consistently euploid (low CDO and CSV), such as *C. bombi* Chr 6, *L. donovani* chrs 19, 28, 30, 34, 36, and *T. cruzi* chrs 20, 26, 32, 34, 36, 40 (full data can be seen on the Supplementary Table 5). There could be some constraints in allowing extra copies of those mainly euploid chrs, or the conditions where they might be advantageous were not met in the samples used in the present study. There was no strong evidence for consistent CCNV in any *T. brucei, T. congolens*e or *T. vivax* chrs.

To study chrs that consistently had extra copies, we selected only chrs with an CDO higher or equal to 1.4 (close to at least trisomic in a diploid organism) (Figure 2 A), and compared their copy number to the combination of all other chrs (Figure 2 B). This analysis revealed 16 consistently duplicated chrs, comprising two in *C. bombi* (0, 9), one in *C. fasciculata* (27), one in *Endotrypanum* (31), one in *L. donovani* (31), one in *L. major* (31, as described previously) [23], four in *Leptomonas* (12, 29, 32 and 39), three in *Paratrypanosoma* (36, 45 and 64), two in *Porcisia* (30526 and 30534), and one in *T. cruzi* (31 as described previously) [24]. The CDO of these chr had values close to 1.5 to 2, meaning 1.5 to two copies for each haploid genome copy. Taken together, these results suggest that there are some chrs that are consistently expanded in each of these clades, and that these chrs are trisomic or tetrasomic. There were no chrs with an CDO higher or equal to 1.4 in *T. brucei, T. congolens*e or *T. vivax* (Figure 2C), consistent with our previous analysis suggesting aneuploidy is rare in these species.

### 3.3 Chrs that are consistently duplicated in other clades are syntenic to *Leishmania* chr 31

Chr 31 (chr30 in *L. mexicana* complex) is consistently expanded in all *Leishmania* species and isolates evaluated to date [15,23,32]. As we observed similar elevated somy for selected chromosomes in other aneuploid trypanosomatids, we evaluated the orthology among these chrs (Figure 3, Supplementary Figures 2 and 3). This was done by evaluating both the proportion of genes from *L. major* chr31 that had orthologs/paralogs in a given chr, and the reciprocal measure: the proportion of genes in the constantly expanded chrs in each clade that had orthologs/paralogs in *L. major* chr31. This analysis revealed that the majority of consistently expanded chrs shared a large number of orthologs with *L. major* chr31. We observed that between 22 (in *P. confusum*) to 91% (in *L. donovani*) of the 352 *L. major* genes had an ortholog in the consistently expanded chrs from the other clades. Similarly, 52 (*C. fasciculata*) to 91% (*L. donovani*) of genes from the expanded chrs in these other clades had an ortholog/paralog in *L. major* Chr31. These values vary, likely due to differences in gene numbers and potential gene duplications in these chrs among different clades (Supplementary Figure 3). Interestingly, even the typically euploid *T. brucei* and *T. vivax* had Leish Chr31 genes with orthologs simultaneously in two chrs: 04 and 08, as described in [59,60]. The exceptions were *Leptomonas* chr 39 and *Porcisia* CM030526, which were also consistently with extra copies, but had no Leish Chr31 ortholog genes. Differently to the chromosomes that were syntenic to Leish Chr31 and were expanded in all isolates, *Leptomonas* chr 39 was expanded in 10 from the 15 isolates, and *Porcisia* CM030526 was in only 4 of the 5 evaluated isolates (Supplementary Figure 1). *Leptomonas* chr 39 also had the lowest median copy number from the consistently expanded chromosomes (Supplementary Table 5). This suggests a different origin for these expansions in *Leptomonas* and *Porcisia*. Nonetheless, the widespread sharing of orthologous genes suggests the expansion of Leish chr31 is not a recent feature of *Leishmania*, but instead this chr expansion is ancient, predating the separation of the trypanosomatid clades and the genome re-organization seen in *T. brucei*, and persisting across this extensive evolutionary history. Due to this finding, we went on to ask if we could find features of this chr, which will be collectively called “*Duplicated31*”, that reflect this ancient duplication

**Figure 3:**
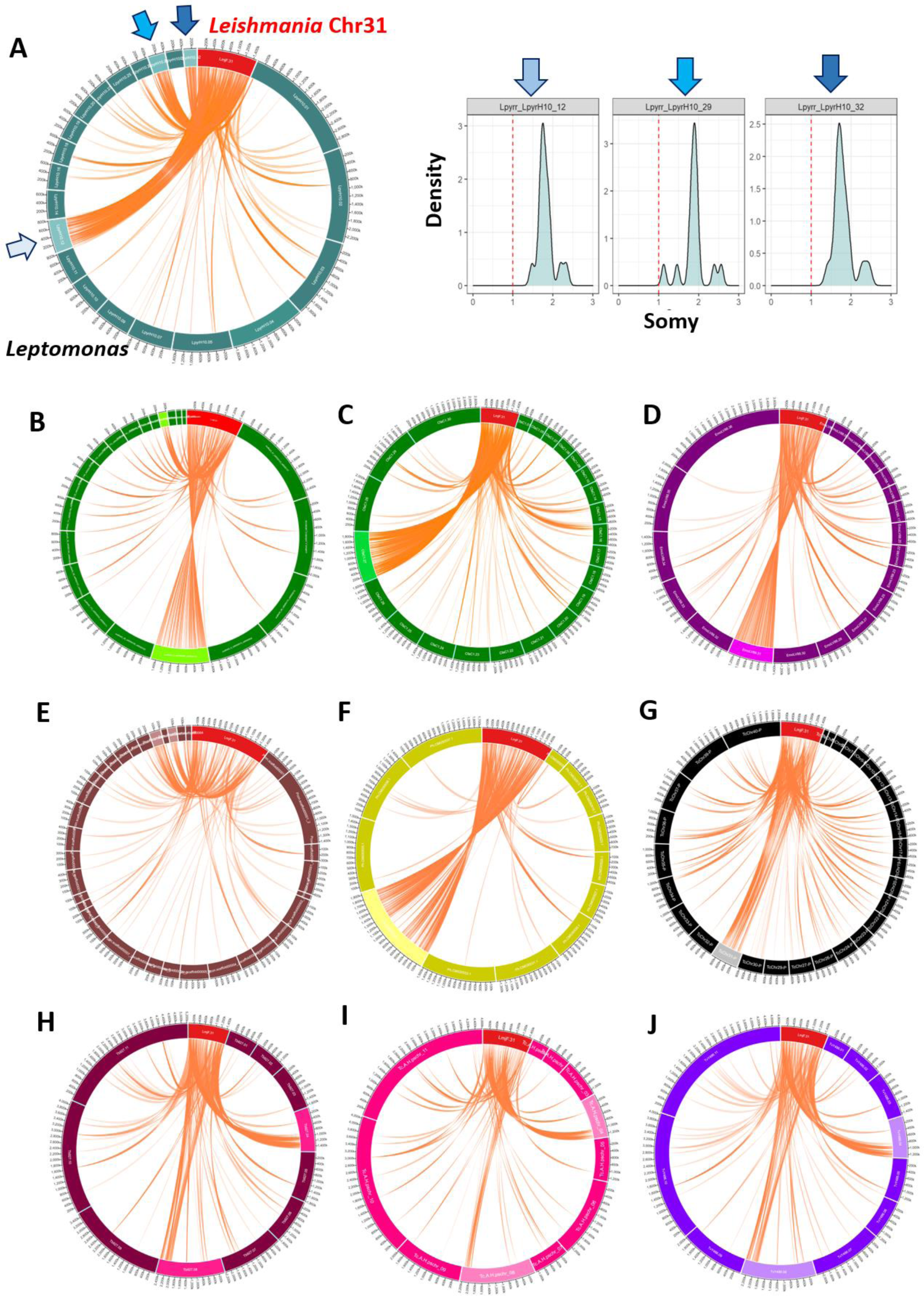
Chrs with a high density of ortholog genes with *Leish* Chr31 also present extra copies in other trypanosomatids. Circa plots representing the orthology between Leish Chr31 (red box) and chromosomes from other species (coloured boxes), drawn in proportion to their sizes. Only chromosomes that shared at least one ortholog gene with Leish Chr31 are shown. Chromosomes that are consistently supernumerary are highlighted by lighter colours. The presence of ortholog genes between Leish Chr31 and a given chr from other species are shown by linking lines. We observed that chrs that were always supernumerary in other trypanosomatids often had a high density of orthologous genes with Leish Chr31. **(A)** *Leptomonas*, showing the circa plot and the distribution density of the somy from chromosomes with syntenic genes to Leish Chr31. Circa plots for **B)** *C. bombi*; **C)** *C. fasciculata* **D)** Endotrypanum; **E)** Paratrypanosoma; **F)** Porcisia; **G)** *T. cruzi*; **H)** *T. brucei*; **I)** *T. congolense*; **J)** *T. vivax*.

### 3.4 *Duplicated31* chr has a peculiar gene structure and an increased nucleotide diversity

We first evaluated structural and functional characteristics of *Duplicated31*, including gene content, nucleotide diversity and gene expression regulation. The majority of genes from Duplicated31 are in the same coding strand, which is unusual for larger chrs in these protozoans, but not exclusive (Figure 4 and Supplementary Figure 4). Of the 352 genes annotated in *L. major* chr31, 349 are in the same coding strand. Similar proportions can be observed in other *Leishmania* species, as well as in other evaluated clades: *C. fasciculata* (518/522), *L. major* (349/352); *L. donovani* (329/332); *Leptomonas* (499/502); *Porcisia* (290/295); *P. confusum* (111/113); *T. cruzi* (234/268). Strand switch regions (SSRs) are important in Trypanosomatids, acting as replication origins and as the transcription start and termination sites for polycistrons [61–64]. Hence, having almost all genes in the same strand could impact DNA replication or gene expression. However, no large difference was observed in the number of polycistronic transcription initiation sites in *L. major* Chr31 when compared with other chromosomes with similar sizes [65].

**Figure 4:**
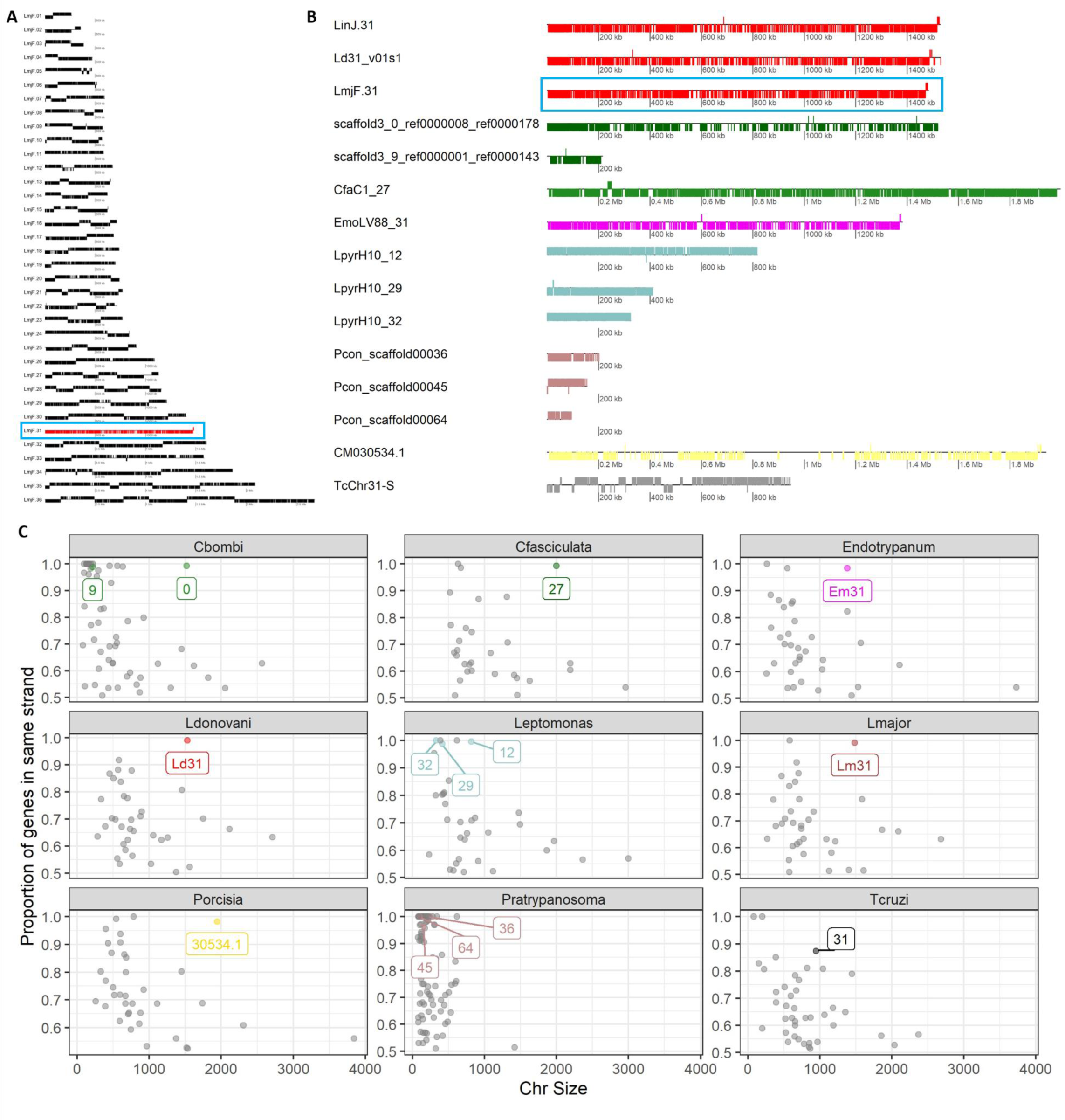
The majority of genes from chrs syntenic to Leish Chr31 are on the same coding strand. **A)** Gene disposition in *L. major* chromosomes. Each box corresponds to a gene, and the position of the box below or above the strand corresponds to the gene orientation. Leish Chr31 is highlighted in red and by a blue box. **B)** Gene distribution of all Duplicated31 chrs. The colours represent the species/clades of origin of each chromosome. All the genes are usually in the same coding strand. **C)** Comparison of the proportion of genes in the same coding strand (Y axis) and chromosomal size in Kb (X axis) for each clade. Each dot corresponds to one chr. The “Duplicated31” chrs are labelled and coloured.

One anticipated consequence of maintaining a duplicated chr long-term is a higher nucleotide diversity (π). Assuming that mutations occur in random positions and in the same frequency across the genome, having twice the copies would result in approximately twice the number of segregating sites (genomic positions that vary in the population) and also increased nucleotide diversity. For this reason, nucleotide diversity was estimated in the three aneuploid lineages with more than 10 isolates, *L. donovani, C. bombi* and *Leptomonas* (Figure 5). Even though *T. cruzi* had a larger dataset, it was not used in the diversity analysis as the massive expansion of multigene repetitive family clusters in this clade could compromise genome-wide diversity estimations. As expected, there were a higher number of segregating sites in chrs 31 *L. donovani*, “0” in *C. bombi* and 12 in *Leptomonas* (Supplementary Table 6), and an increased mean nucleotide diversity in all when compared to the combination of all other chrs, using 10kb sliding windows: *L. donovani* (*p-value* = 1.5 x 10^-47^), *C. bombi* (p-value 3.47 x 10^-39^) and *Leptomonas* (p-value 1.08 x 10^-12^). In fact, the mean Duplicated31 diversity was around twice that observed for other chrs in *L. donovani* (Duplicated31: 0.00221; others: 0.000973); around 1.5x for *C. bombi* (Duplicated31: 0.00331; others: 0.00207); and 1.14x in *Leptomonas* (Duplicated31: 0.00786; others: 0.00692). These findings are in accordance with the proposed long-term maintenance of extra-copies of Duplicated31.

**Figure 5:**
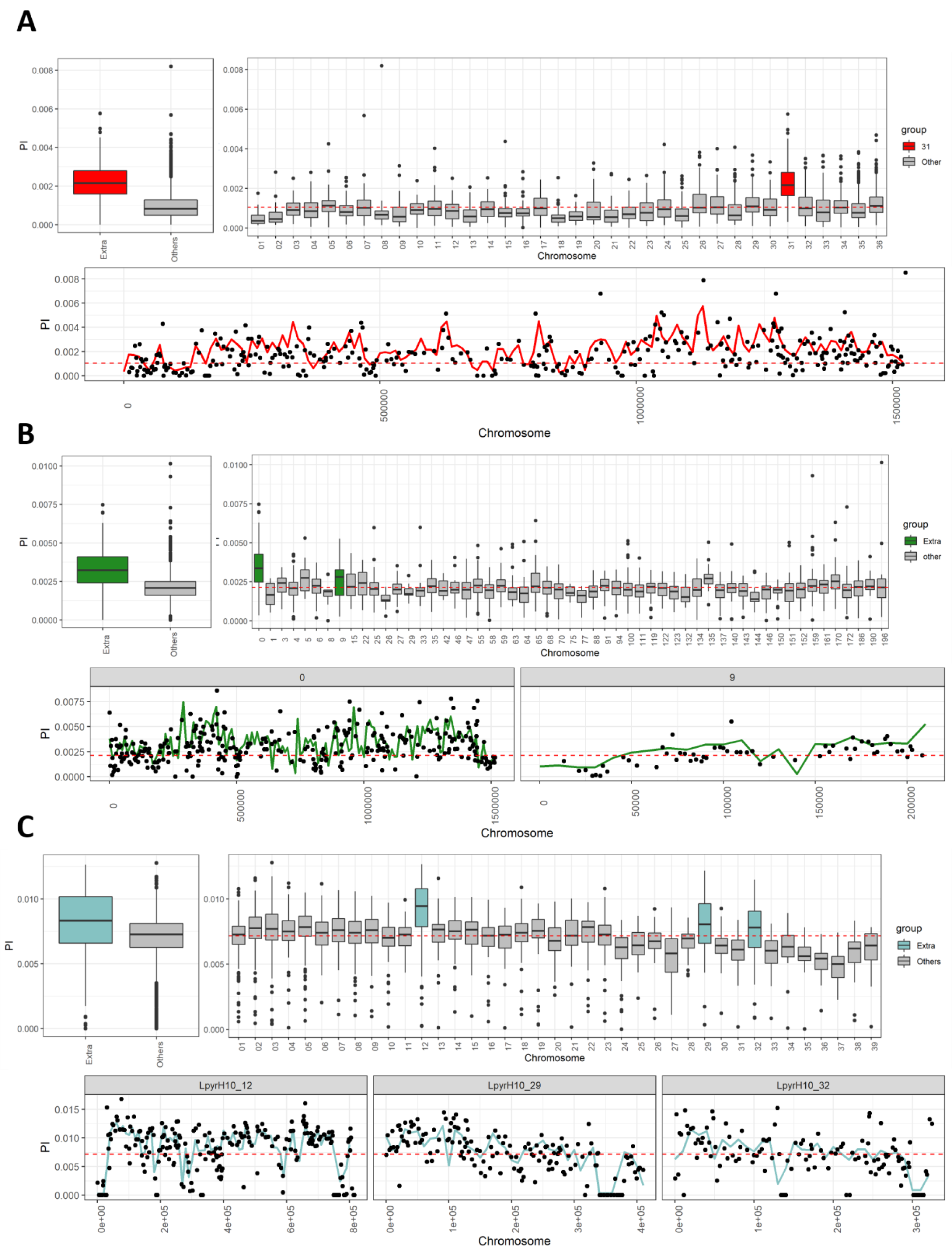
Chrs syntenic to LeishChr31 have a higher nucleotide diversity (π). Top panels are box plots representing 10kb window π values comparing Duplicated31 with others (left panel) or in each chr/scaffold (right panel). Chrs from Duplicated31 are coloured, while other chrs are grey. The bottom panel represents the π along the chr/scaffolds from Duplicated31, where the line represents the 10kb window π, while the dots correspond to the π value in each gene in this chr/scaffold. In both panels, the red dashed line represents the mean π in all chrs. **A)** *L. donovani* EA_HIV set; **B)** *C. bombi*; **C)** *Leptomonas*.

Next, we evaluated gene-by-gene nucleotide diversity, as well as the diversity along the chr in Duplicated 31, to assess if the observed higher diversity was caused by short regions of high diversity or was evenly distributed along the chromosomes. Nucleotide diversity was similar along Duplicated31 sequence for the three clades (Figure 5), suggesting that the observed higher diversity in these chrs was not caused by a few highly polymorphic regions. We identified some genes with a higher nucleotide diversity than expected in Duplicated31 for the three clades (higher than the mean+3SD from the genomic mean π). In *L. donovani*, from the 35 highly diverse genes, 10 are pseudogenes, and 20 are hypothetical proteins (with 5 hypothetical proteins that are pseudogenes). From the 10 annotated non-pseudogenes, two were 2Fe-2S iron-sulphur cluster binding proteins, and the other were proteins enrolled in cellular/vesicular transport, lipases, peptidases, RNA binding proteins and ribosomal proteins (Supplementary Table 7). For *Leptomonas*, from the 20 highly diverse genes, 17 were hypothetical proteins, and the other three were a lipase precursor, an amino acid transporter ATP11 and a ferrochelatase-like protein. The *C. bombi* sequences were not annotated (Supplementary Table 7). It is not surprising that the majority of the most diverse genes are pseudogenes, as those are usually under no or weak purifying selection, accumulating more mutations than other genes. The large number of highly diverse hypothetical proteins identified in Duplicated31 highlights the importance of better characterising Trypanosomatid proteins as these might be important for specific processes in the parasite biology.

We observed a significant positive correlation between the population mean chromosomal copy number for each chr and its nucleotide diversity, when all chrs were evaluated (Figure 6). However, this correlation was lost when the Duplicated31 chrs were removed. This suggests that ploidy variation for the other chrs is transient, and not maintained for a long enough period of time to impact their overall nucleotide diversity.

**Figure 6:**
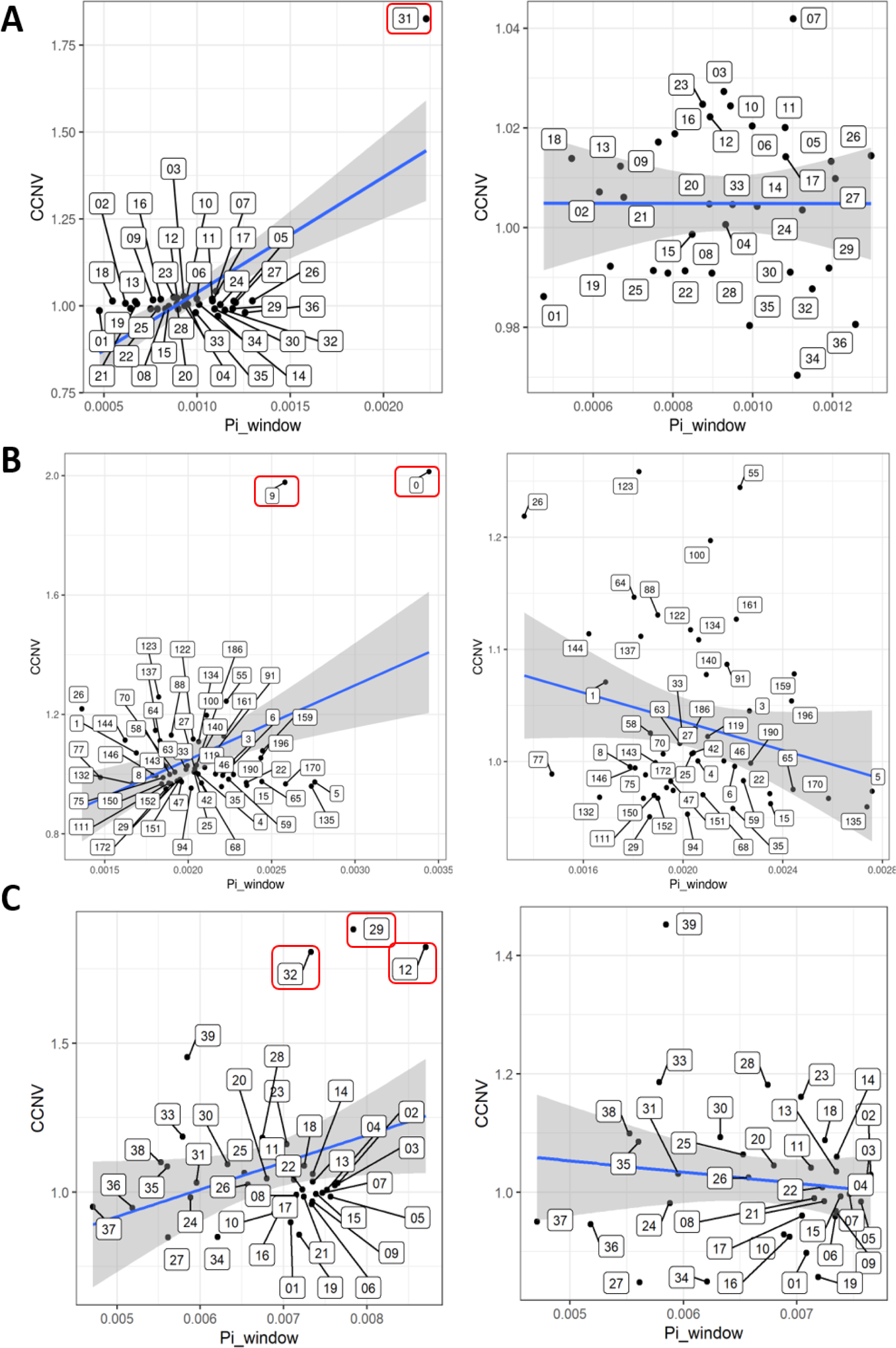
There is no correlation between CCNV and nucleotide diversity (π) without the Duplicated31 chrs. Correlation between CCNV (Y axis) and π (X axis) in all chrs (left panel) and without the Duplicated31 chrs (right panel). The Duplicated31 chrs are highlighted by red boxes. **A)** *L. donovani* EA_HIV set (left panel: r =0.719, p-value 7.58x 10^-7^), (right panel: r = -0.001, p-value = 0.9943); **B)** *C. bombi* (left panel: r =0.429, p-value 0.001082), (right panel: r = -0.229, p-value = 0.09815); **C)** *Leptomonas* (left panel: r =0.309, p-value 0.05542), (right panel: r = -0.133, p-value = 0.4392).

### 3.5 Duplicated 31 has a higher increase in segregating sites in intergenic regions, but also in synonym/non-syonym sites

We next went on to evaluate the biological impact of the segregating sites (genomic positions that vary in the population) in Duplicated 31. We used *L. donovani* for this analysis, as it has the most accurate genome assembly and annotation from the three species used in the nucleotide diversity (**π**) assessment. Segregating sites were separated based on their potential biological impact: missense (amino acid altering), synonymous (no amino acid alterations) and intergenic (occurring outside annotated coding sequences (CDS) regions). In doing so, an increase in SNP counts was found for all three classes in Chr31, which was not impacted by using diploid or tetraploid SNP-calling methods. Chr31 had an increase of SNP counts/kb in all three scenarios when compared to other chrs, which was more prominent in intergenic regions (1.8x), when compared to synonymous (1.41x) and non-synonymous (1.58x) sites (Figure 7A and B and table 3). This is expected, as missense SNPs are usually more deleterious than synonymous or intergenic mutations, and should be rarer in the population.

**Figure 7:**
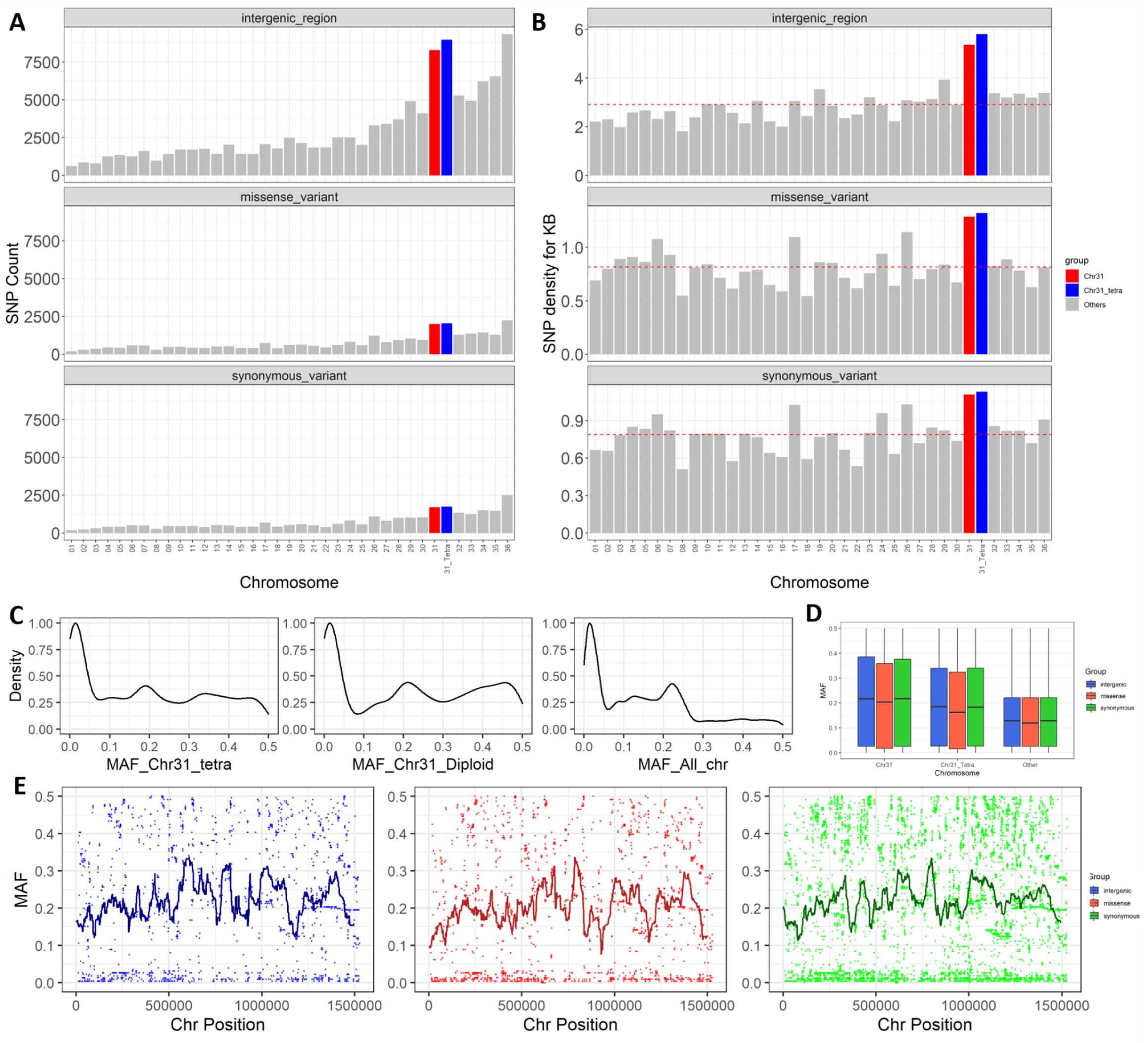
Comparison of MAF in Chr31 in *Leishmania*. **A)** Number of Intergenic non-synonymous and synonymous SNPs across chrs. **B)** SNP density/Kb for each chr. The red dashed line corresponds to the mean density for all chrs. Chr31 SNPs called with diploid or tetraploid SNP-calling methods are respectively in red or blue. **C)** MAF distribution in all SNP positions. The panels from the left to right correspond to: Leish Chr31 using a Tetraploid SNP caller; Leish Chr31 using a diploid SNP caller and all chrs. **D)** Evaluation of MAF in Intergenic (blue), non-Synonym (red) or Synonym (green) positions, comparing all chrs with Leish Chr31 with both using diploid or tetraploid SNP-calling methods. **E)** MAF along Chr31 with diploid SNP-calling method, for Intergenic (blue), non-synonymous (red) or synonymous positions (green). Each dot corresponds to a SNP position. The line corresponds to the average MAF in a sliding window of 50kb, with increments of 1kb.

**Table 3:** Segregating sites per kb.

|  | **31 Diploid SNP calling method** | **31 tetraploid SNP calling methods** | **Mean of all chromosomes** |
| --- | --- | --- | --- |
| Intergenic regions | 5.36 | 5.80 | 2.91 |
| Synonymous SNPs | 1.10 | 1.13 | 0.78 |
| Non-Synonymous SNPs | 1.28 | 1.32 | 0.81 |

To evaluate if having consistent extra copies of a chromosome could impact the retention of rare alleles, as they could be maintained in a lower proportion in a given isolate (3 copies of one allele and 1 copy of the rare allele/cell) we estimated the Minor allele frequency (MAF) in Leish Chr31 both with diploid or tetraploid SNP callers and compared its values with the obtained for other chromosomes. There was a similar distribution of MAF values when the SNP calling was performed with diploid or tetraploid callers, with a peak in low values but a higher tail towards 0.5 MAF when compared with the combination of all other chromosomes (Figure 7C). This finding might suggest that there are more balanced alleles in Chr31, with a higher proportion of alleles close to 0.5 when compared to other chrs, which suggests that there might be sites under balancing selection in this chromosome [66]. There was no clear impact in the MAF estimations of using diploid or tetraploid SNP callers, where the lowest MAF for all was obtained for missense regions (Figure 7D). The overall MAF appears to be similarly distributed along chr 31 (Figure 7E), reinforcing that differences between this chr and others are not caused by spikes of high MAF. There was also no bias in nucleotide diversity using diploid or tetraploid SNP callers which resulted in similar chromosomal π, with values of, respectively, 0.0022 and 0.0021.

### 3.6 Gene expression from *Leishmania* chr31 is controlled by downregulating all chromosomal copies to a similar extent

Previous works from Barja 2017 [19] and Dumetz 2017 [14], have demonstrated that having extra chromosomal copies in *Leishmania* results in an proportional increase in overall gene expression. The only exception to these observations is Leish chr31, which has a gene expression similar to disomic chrs despite having 3 to 4 chromosomal copies. How gene expression control is achieved on Leish chr31 is unknown. We propose two main hypotheses for this regulation. 1: *Leishmania* downregulates the expression of all the ∼four copies of Chr31 to a similar extent; 2: *Leishmania* silences two whole chromosomal copies, leaving two copies normally expressed. To investigate how this silencing occurs, we used the 7 available pairs of WGS and RNAseq data described in Barja 2017 [19], and estimated variation in the proportion of reads in heterozygous positions in both data sets.

We initially confirmed results from Barja and colleagues, supporting that the expression profile of Chr31 is similar to a disomic chromosome despite its supernumerary nature (Figure 8A). Using the same dataset, we evaluated if the Alternate Allele Read Depth (AARD) in heterozygous positions from the WGS data correlates with the AARD from the RNAseq data (Figure 8B and C and Supplementary Table 3). A positive AARD correlation between WGS and RNAseq data is an indication that *Leishmania* down regulates all the copies similarly. In fact, a strong positive correlation (r∼0.88) between paired WGS and RNAseq AARD from 7 *L. donovani* clones was observed, strongly suggesting that all copies are being silenced similarly. This is supported by the evaluation of AARD variation along the chr (Figure 8C). Another, potential scenario is if each individual in a *Leishmania* population (in this case, each cell in this cloned population) silences two chromosomal copies differently and independently of the others. Collectively, this might also result in a positive correlation of AARD between WGS and RNAseq, if the silencing is random. However, for this to happen, the silencing of each chr copy would have to change at least during each cell replication, as this analysis was done with cloned lineages. This regulation does not appear to be caused by codon usage, as there is no significant difference among *Leishmania* chrs (Supplementary Figure 5).

**Figure 8:**
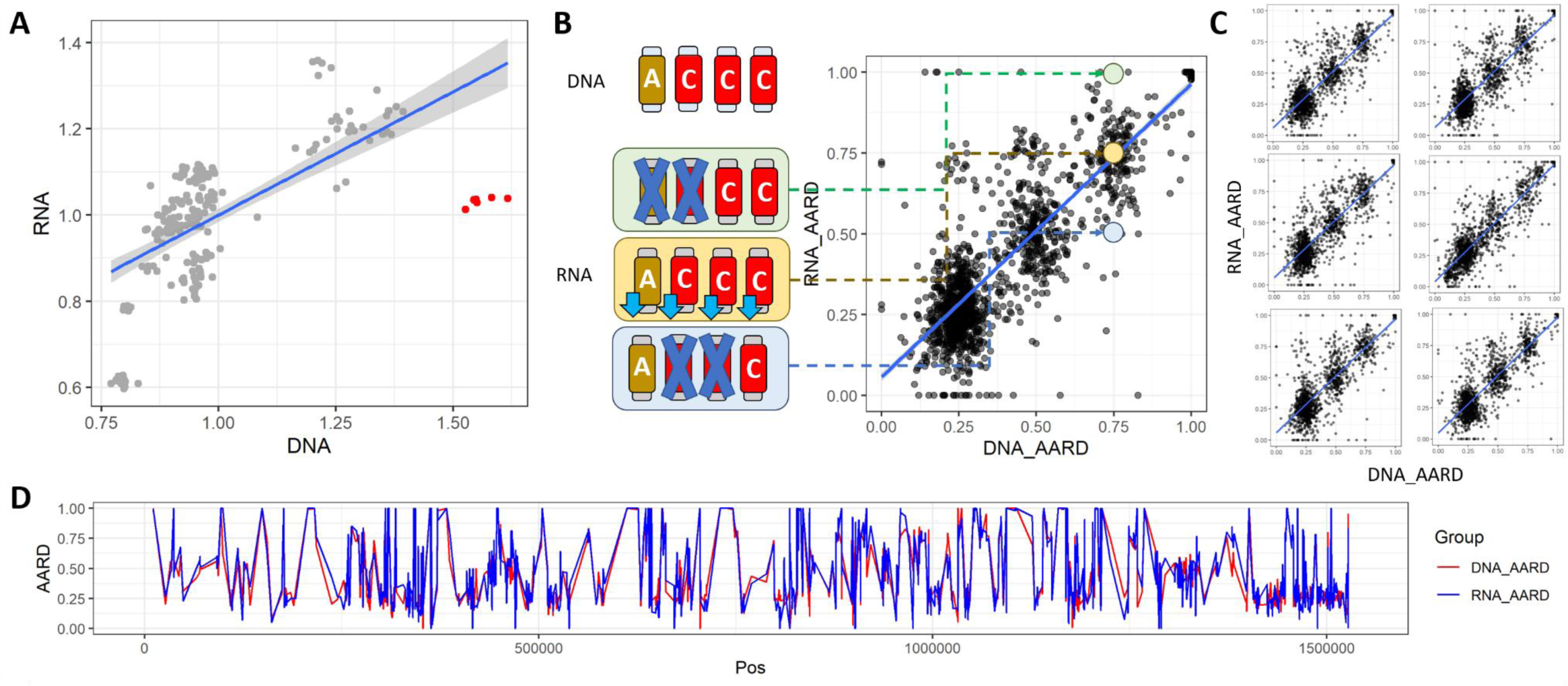
Leish Chr31 gene expression is similar to disomic chrs, and the four copies appear to be downregulated similarly. **A)** Correlation between the DNA copies (X axis) and RNA expression (Y axis) of all 36 *Leishmania* chrs in 7 clones (r=0.647994, p-value 4.14e-27). Leish Chr31 is highlighted in red. This image is a confirmation of the results described in Barja 2017 [19]. **B)** Correlation between DNA AARD (X axis) and RNA AARD (Y axis) for each SNP position in Clone 3 from the tetraploid Leish Chr31. Each dot represents one SNP position from the Leish Chr31 in the Clone 3 (C3) (r=0.872, p-value 0). On the left, a scenario where four alleles are observed “A”, “C”, “C”, “C” and “C” is the alternate allele; therefore WGS AARD = 0.75%. If the gene expression of all chr copies is silenced similarly, the AARD from the RNAseq data for the same position would also be around 0.75 (yellow box and dot). If two chr copies are silenced, this would result in “AC” with AARD of 0.5 or CC with ARDD of 1.0 (green and blue boxes and dots). **C)** AARD DNA-RNA correlation plots for the Leish Chr31 in clones C4 (r=0.875, p-value 0); C6 (r=0.868, p-value 0); C7 (r=0.869, p-value 0); C8 (r=0.905, p-value 0); C9 (r=0.866, p-value 0) and C10 (r=0.887, p-value 0). **D)** DNA (red line) and RNA (blue line) AARD along Leish Chr31 in C3. The other 6 clones can be seen in the Supplementary Figure 6).

### 3.7 Biological functions in genes and regions from Duplicated31

We next evaluated potential biological functions for genes in Duplicated 31, to assess if there is any enriched function in this constantly expanded chr. This analysis was done with *Leishmania donovani* Chr31, *T. cruzi* Chr31 and *T. brucei* (regions from chr04 and chr08 that are syntenic to Duplicated31), as these genomes have curated annotations. We separated the dataset in three main groups: **1: Conserved_all:** genes that were present in orthogroups containing the three clades; **2:Conserved_pair:** genes in orthogroups containing only two clades; and **3: Exclusive:** genes that were exclusive for each clade. We compared the gene ontology from these datasets with a dataset containing all genes for the three species.

When the genes from the Conserved_all dataset were evaluated, the biological functions that were enriched were from basal cellular metabolism, such as: “inner dynein arm assembly”, “negative regulation of transcription by RNA polymerase III”, “histone H3-S10 phosphorylation”; “mitochondrial respiratory chain complex III”, “endoplasmic reticulum lumen”, “glucosidase activity” and “ubiquinol-cytochrome-c reductase activity”, among others (Supplementary Table 8).

For group 2 “Conserved_pair”, there was an enrichment of GO terms related to amino-acid transport for the pair *T. cruzi* and *L. donovani*, which might be important for their intracellular survival. But the most striking enrichment was the “GPI anchor biosynthetic process” (∼25% of the genes), “protein glycosylation” (∼25% of the genes) and “UDP-glycosyltransferase activity” (∼43% of the genes), obtained for the pair *T. brucei* and *T. cruzi*. The GPI anchor is a crucial structure in these protozoans, and anchor to the protozoan outer membrane proteins encoded by multigene families that are virulence factors, such as the Variant Surface Glycoprotein (VSG) from *T. brucei*; and Mucin (TcMUC), Mucin Associated Surface Proteins (MASP) and Trans-Sialidases (TS) in *T. cruz*i. Some of these proteins, such as Mucins and VSGs are known to be glycosylated. This suggests that genes in Duplicated31 could be important for the GPI assembly and glycosylation in these protozoans. There was no relevant function enriched in the *T. brucei* - *L. major* pair.

Finally, for the group 3 “Exclusive”, there were only a few functional enrichments, such as: “phosphatidylcholine biosynthetic process” for *L. major*; “cAMP biosynthetic process” for *T. brucei*; and “arginine biosynthetic process” and “deacetylase activity” for *T. cruzi* (Supplementary Table 8). It is worth mentioning that some important drug resistance targets in *Leishmania* are also found in this chr, such as the Miltefosine Susceptibility Locus [67] and the Aquaglyceroporin gene [68], and they are all a result of gene/segmental loss.

Taken together, these results suggest that the genes in Duplicated31 that were shared among species are enrolled in cellular basal metabolism, and that some species, such as *T. cruzi* and *T. brucei* have transferred genes related to their parasitic lifestyle to this chr.

## 4. Discussion

By evaluating chr ploidy across genera that span the Trypanosomatid grouping, we have revealed three features of aneuploidy in these single celled parasites. First, aneuploidy is an ancestral characteristic of trypanosomatids, suggesting it is central to their genome functionality. Second, *T. brucei* and related African trypanosomes have more recently evolved to largely dispense with aneuploidy, perhaps reflecting genome reorganisation. Third, we have identified the presence of an ancestral chromosomal duplication, named collectively as “Duplicated31”, which has been maintained throughout Trypanosomatid evolution either as a >diploid chr or a syntenic duplication in two chrs in African trypanosomes, resulting in higher nucleotide diversity, unusual gene organisation and expression regulation.

Previous works have described aneuploidy to be found in distant trypanosomatid clades, mainly represented by several *Leishmania* species [23]), and *T. cruz*i subgroups and field isolates [24,25]. Taking advantage of near complete genomes of clades across the trypanosomatids, representing both dixenous and monoxenous species, and including the basal protozoan *P. confusum* [28], we reveal that aneuploidy is widespread throughout these protozoans. This is the most comprehensive evaluation of aneuploidies across trypanosomatids to date. Our analysis shows that the mosaicism of chr variation is largest in *Leishmania* and *Leptomonas*, where some of their isolates had the highest level and variation in chr expansions among the evaluated groups. Beyond these clades, isolate-specific data indicates that the overall level and variation in chr expansions is low, which suggests that while aneuploidy is present, it is normally limited to only a few chrs with extra copies in the same isolate. Tolerance to aneuploidy is a balance of the fitness cost of chromosomal duplication contrasted with the fitness advantage from increased dosage of specific genes [1]. Hence, the low number of chrs with extra copies in a given isolate suggests that the increased fitness cost that they incur usually outweighs the fitness gain of having specific genes in extra copies [1]. What aspects of *Leishmania* and *Leptomonas* biology tip this balance towards greater tolerance of increased ploidy is unclear. We have also observed that, in the aneuploid clades, monosomies are less common than chromosomal gains. This could be caused by haploinsufficiency of genes and unmasking of recessive mutations, which make monosomies highly deleterious [69,70]. Lower IGE and IAL were observed in isolates from *T. brucei* and closely related clades, which indicates that these clades are unusual amongst trypanosomatids, by being mostly euploid (discussed further below) [26,71]. However, some aneuploid and/or polyploid isolates from these species have been described in the literature [27,72]. The widespread presence of aneuploidies across distantly related clades, as well as the restricted but not absent occurrence of aneuploidy in *T. brucei*, strongly supports that aneuploidy is an ancestral characteristic, controlled in the *T. brucei* and closely related protozoans.

Even though aneuploidy is observed in the majority of trypanosomatid clades, the number and level of chromosomal duplications vary between species and even among individuals from the same species. This suggests that there might be different levels of tolerance to aneuploidies in these parasites, where *Leishmania* and *Leptomonas* are among the most permissive clades, and *T. brucei* and related parasites the most restrictive. The within species difference in tolerance to aneuploidies could be caused mainly by environmental pressures, where more permissive environments (as cell culture or the insect vector) allows for the expansion of different aneuploidy patterns generating a mosaic population, and more selective environments as the mammalian host, selects for specific chromosomal expansion patterns. This is in agreement with what was shown by Dumtez and colleagues, where there were less chromosomes with extra copies in *L. donovani* in the mammalian host than in the cell culture [14]; and with Negreira 2022, which showed that there are variations in ploidy patterns in single-cells that were originated from clonal samples [34]. However the strong reduction of aneuploidies in *T. brucei* and closely related parasite clades suggests that specific genomic modifications in this group are incompatible or at least restrict chromosomal duplications. There are several factors that might be related to the difference in tolerance to aneuploidy, such as: **1-Megabase chr sizes:** *T. brucei* and closely related protozoans have their genome organised in 11 large ∼mega-based size chrs, while *T. cruzi* and *Leishmania* have divided in respectively ∼41-47 and 34-36 smaller chrs [50,52,57,73–77]. Hence, the presence of one chr with extra copies in *T. brucei* might, on average, amplify more genes than one in *T. cruzi* or *Leishmania*. In this scenario, the fitness cost of having unwanted genes amplified could outweigh the benefits of having just few beneficial genes with extra copies, as proposed in yeast [1]. Why *T. brucei* and close relatives might have evolved a smaller repertoire of larger chrs is unclear, but our data shows that, as part of the reorganisation, parts of the ancestral Duplicated31, (see below), was maintained in increased copy number through duplication in chrs 4 and 8. **2-Minichromosomes and intermediate chromosomes**: A mitigating factor for the lack of aneuploidies in *T. brucei* might be presence of “*minichromosomes*”, which are short sequences between 30-150 kb, with an estimated number of ∼100 copies per cell. Intermediate chromosomes are larger than minichromosomes, but observed in lower copy numbers (reviewed in [78]). These short chrs usually contain one or two genes coding for Variant Surface Glycoproteins (VSG), that are important in the antigenic variation process in these parasites (Reviewed in [79]). These minichromosomes could potentially vary in copy and content among *T. brucei* cells, mitigating the need of whole chr aneuploidies. However, to date, only VSG genes and VSG transcription sites have been found to be present in minichromosomes, so how this would contribute to adaptive gene expression is unclear. **3-Differences in DNA replication regulation**: There are several differences in the DNA replications between *T. brucei* and *Leishmania* (reviewed in *[17]*). MFA-seq experiments have shown that while *T. brucei* have several replication origins in each chr, *Leishmania* appears to have only one preferential site during the S phase, which is less than the number of origins needed to timely replicate long chrs [59]. To mitigate this, while the duplication of the core genome is confined to the S phase in *Leishmania*, the duplication of the sub-telomeres occurs in late S and G2/M and G1 stages of the cycle [80]. Replication outside S phase is common in aneuploid cells, being also observed in aneuploid cancer [81,82]. Short nascent DNA strand sequencing (SNS-seq) has suggested the presence of several alternative origins of replication in *Leishmania* [83], which could be differently activated in a cell population [17]. This might result in diversity of replication and potentially mosaic aneuploidy, which could explain the within clone variation described in [34]. Hence, the differences in the susceptibility to aneuploidy could be directly related to the protozoan genome organisation and replication processes.

By evaluating the chromosomal duplications, we have shown that the majority of chrs with consistent extra copies in all evaluated species are syntenic to Leish Chr31, now collectively called Duplicated31. This suggests that the expansion in the copy number of this chr predates the speciation of the evaluated clades, and was maintained across their evolution. The biological relevance of this chromosomal duplication is further supported by the presence, even in the mainly euploid *T. brucei*, of regions of Leish Chr31 duplicated in two different chrs. This ancient duplication is further confirmed by the increased nucleotide diversity in Duplicated31, evaluated in *C. bombi, L. donovani* and *Leptomonas*. Even though the presence of extra copies of Chr31 and the synteny with regions from *T. brucei* were already known [19,23,59,60], our data reveal for the first time that this stable increased ploidy is a universal feature of trypanosomatids. There appears to be no equivalent stable increased ploidy in any other chr in any trypanosomatid we have tested, which suggests that, at least for the evaluated populations, the presence of extra copies from other chrs is transient and potentially context dependent. When discriminated by biological impact, the highest increase of segregating site counts in Leish Chr31 when compared with other chrs was in intergenic regions and pseudogenes. This is in accordance with what is expected in long term duplications, as mutations in intergenic regions and pseudogenes are usually less impactful and more frequently retained than the ones in protein coding regions [84,85].

One striking feature of Duplicated31 consistent tetraploidy is that it appears to have impacted both the gene structure and the gene expression control. Although consistently having extra copies, Leish Ch31 is usually expressed as if it is a disomic chr [14,19]. As this chr duplication is ancient, these parasites might have developed a mechanism to control its gene expression, potentially counteracting the deleterious effects of chr-wide consistent unbalanced gene copies. This expression control could be generated by fully silencing chromosomal copies, as seen in the human X chr silencing, mediated by the ncRNA XIST and irreversible heterochromatization [86–88]; or by downregulating similarly different chromosomal copies, by a still unknown mechanism. By comparing the AARD in heterozygous positions in DNA and RNAseq along Leish Chr31 in clones (data from [19]), we have shown that, at least at the population level, *L. donovani* potentially regulates the gene expression by similarly downregulating all gene copies; and that this appears to be independent of codon usage variation [89]. This control could be mediated by chr packing, transcription rate, or by rapid mRNA decay. With the current data, we cannot completely rule out that within a culture flask, cells might silence Chr 31 copies differently, resulting in a global similar level of expression of all chromosomal copies. If that is the case, the rate in which the chr silencing changes in *L. donovani* would have to be high, to match the WGS data, as this analysis was done in cloned isolates. Another potential point of control of Leish Chr31 expression could be its nuclear compartmentalization, which has been already shown to be relevant for gene expression in *T. brucei* [90,91], where expressed chromosomal copies could be in different compartments when compared to non-expressed copies. Another peculiarity of Leish Chr31 (and in all Duplicated31) that could impact expression levels is the majority of its genes being in the same coding strand. Strand switch regions are important in Trypanosomatids, being relevant in defining the end of polycistronic units and DNA being replication origins [61,62]. Hence, having practically all the genes in the same coding strand could ease the gene expression regulation control of this chr. However, no bias was observed in polycistron numbers in *L. major* Chr31 chr [65]. Nevertheless, regardless of the mechanism of expression control, at least at the population level, the gene expression of Leish Chr 31 is consistent with a similar downregulation of all their genome copies.

As usually chromosomal duplications are linked to increased gene expression [1,14], the apparently paradoxical finding in Leish Chr31 where it has consistently extra copies but regulates gene expression to dissomic levels is intriguing, and requires further studies. One possible explanation is that the peculiar gene structure in this chromosome (all genes in the same coding strand) somewhat complicates the gene expression, requiring four copies to achieve a similar to diploid expression. Careful evaluation of gene expression mechanisms in this chromosome, as well as the presence of expression-blocking structures such as R-loops might help in elucidating if this control occurs at the transcription level. Another possibility is that, although the overall chromosomal expression is regulated, Leish Chr31 might keep some key gene expressions in its tetrasomic level. A careful analysis of RNAseq data not only from several *Leishmania* species/isolates, but also other trypanosomatids could show if there are genes that are consistently overexpressed in Duplicated31 across clades. These genes might be crucial for trypanosomatid basal functions. On the other hand, the evolutionary pressure to keep extra copies of Leish Chr31 might not be related to the need of overexpressing genes at all. Instead, it could be related to increasing sequence variability and allowing neofunctionalization in paralogous genes, as proposed for the *T. brucei* Chr04-08 duplication by Jackson and colleagues [60]. In this scenario, having extra copies could allow mutations to accumulate differentially in each haplotype, and more suitable variants could be expanded by haplotype selection [19]. As there are three to four copies of this chromosome in each cell, it could have a higher recombination rate than other chromosomes, shuffling these variants in combinations with better fitness. In this scenario, the gene expression control could mitigate the deleterious effects of some of the accumulated mutations, as its expression levels would be low. In accordance with this, several important drug resistance mutations are observed in Leish Chr31 (see below).

Finally, we evaluated the biological functions that were enriched in the shared core, or in species-specific regions from Duplicated31, focusing on the ‘TriTryps’, *Leishmania* (chr31), *T. brucei* (Chr4 1mb:1.5mb and Chr8 2mb:2.5mb) and *T. cruzi* (Chr31). The majority of the biological functions that were enriched in the shared content are from basal cellular processes, such as “regulation of gene expression”, “histone phosphorylation” and “glucosidase activity”. One of the striking findings was in the functions that were enriched only in *T. brucei* and *T. cruzi*, which were linked to glycosylation and surface protein anchoring, such as: “GPI anchor biosynthetic process”, “protein glycosylation” and “UDP-glycosyltransferase activity”. In these two protozoans, GPI-anchored proteins play a pivotal role in host-pathogen interactions, including immune evasion and cellular invasion processes [79,92]. UDP-glycosyltransferases catalyse the transfer of N-acetylglucosamine residues from UDP-GlcNAc to phosphatidylinositol, which is crucial not only in the initial steps of GPI anchor synthesis, but also in the synthesis of several glycans in the surface of trypanosomatids [93–95]. In *T. cruzi*, the proteins encoded by the multigene families MASP, Mucin and Trans-Sialidase, which are enrolled in cellular adhesion, invasion and immune evasion processes (reviewed in [96]), are anchored to the parasite surface by GPI-anchors, where Mucins are also highly glycosylated [97]. In *T. brucei*, Variant Surface Glycoproteins (VSG), the protein enrolled in immune evasion by antigenic variation, is also anchored to the parasite surface by GPI [92,98]. Even though not highlighted in the gene function by enrichment analysis in *Leishmania*, several drug resistance genes or regions are found in Leish chr31, such as the Miltefosine Susceptibility Locus (MSL) [67] and the Aquaglyceroporin gene [68], and both a result of gene inactivation or deletion. It is tempting to speculate that chrs with extra copies might be more prone to gene inactivation without loss of function. In this scenario, by having consistent extra copies, the loss of function of some genes in one or two chr copies could be compensated at least partially by the functional gene in the other copies, in natural conditions. In drug pressure or other stressors, if the deleted regions are advantageous, they might be selected and even expanded by mechanisms as the haplotype selection [19]. In fact, if these deletions/frameshifts result in lower fitness, they might be kept in lower dosages (i.e. 1 of the 4 copies for example) in some isolates, mitigating their detrimental effects in natural condition, and increasing in number if the need arises. Aneuploidies are commonly associated with drug-resistance emergence in cancer, where resistant cells harbour recurrent aneuploidies [10], and in yeast [5,99]. Taken together, these results suggest that the expansion of Duplicated 31 impacted shared housekeeping genes, but also host-parasite interaction and drug resistance genes/loci. Moreover, different parasite lineages could have selected or transferred different genes to Duplicated31 depending upon their specific evolutionary needs.

## 5. Conclusions

In the present work, we have shown that aneuploidy is an ancestral characteristic of the trypanosomatid clade, which is limited in the *T. brucei* and closely related parasites, potentially due to genome re-organization and DNA replication mechanisms. In addition, we have also shown that stable ∼tetrasomy of the chrs that are syntenic to Leish Chr31 also predates these clades separation and resulted in a differential gene structure, increased nucleotide diversity and downregulation of gene expression of the four chr copies in a similar proportion, at the population level. Finally, we have shown that while the core shared content of this chr is enrolled in housekeeping functions, some species have transferred genes to this chr that relate to their parasitic lifestyles, such as GPI anchor and glycosylation relevant genes in *T. brucei* and *T. cruzi*. The consistent extra copies of Leish Chr31 could also be relevant for the development of drug resistance, especially by gene function loss, as seen in the MSL locus and the aquaglyceroporin gene. We still do not fully comprehend the processes that govern aneuploidy in these protozoans, nor the molecular mechanisms that regulate the gene expression Leish Chr31. New studies investigating these modifications at the single-cell level will be important to address these unknowns. Moreover, a complete genome assembly of the protozoan *Bodo saltans* could also shed light on the range of aneuploidy presence and Duplicated31 expansion in the wider Kinetoplastid grouping.

## Supplementary Material Legends

**Supplementary Figure 1:** Heatmap (top panel) and density plot (bottom panel) of chromosomal copies in all isolates from all clades. The heatmaps show the sample-by-sample CCNV. Each line corresponds to a different chromosome/scaffold and each column to a different isolate. The chromosome copies are represented by a colour scale from dark-blue (low copy number) to dark-red (high copy number), where euploid chromosomes (∼1 copy for each haploid genome copy) are in light blue. The dendrogram on top clusters the samples by UPGMA, based on the Manhattan distance of their chromosome copy numbers. The density plots represent the deviation from euploidy for each chromosome. Each box corresponds to a given chromosome/scaffold, the X axis to the chromosome somy and the Y axis to the frequency of observations. The red dotted line represents the expected value for euploid chromosomes. Each page corresponds to one clade, on the order: *C. bombi, C. fasciculata, Endotrypanum, L. donovani, L. major, Leptomonas, Paratrypanosoma, Porcisia, T. brucei, T. congolense, T. cruz*i, *T. vivax*.

**Supplementary Figure 2: Proportion of gene sharing between LeishChr31 and the chromosomes in other species.** Each panel corresponds to a different species. The X axis corresponds to the number of shared Orthologs + Paralogs in a given chromosome and the L. major LeishChr31. Chromosomes with consistent extra copies and syntenic to L. major LeishChr31 are highlighted in gold, while other chromosomes are represented in blue. Chromosomes/scaffolds with less than 10 sharings were “grouped” in the “others” column, in grey.

**Supplementary Figure 3: Proportion of gene sharing between LeishChr31 and the chromosomes in other species. A)** Table representing the number of genes (total genes in *L major* chr 31 = 352) in each LeishChr31 syntenic chromosome and how many of these genes have orthologs/Paralogs in L. major LeishChr31. **Chr_genes**: Number of genes in a given chromosome (or combination of chromosomes, as in Leptomonas). **Orthologs**: How many of these genes have orthologs/Paralogs in *L. major* LeishChr31. ***L. major* Orthologs**: How many genes in *L. major* LeishChr31 have orthologs/paralogs with genes in the other species selected chromosomes. **Prop_Lmajor**: The proportion of genes in *L. major* LeishChr31 have orthologs/paralogs with genes in the other species selected chromosomes, **Prop_Other 31**: The proportion of genes in a given chromosome that have orthologs/Paralogs in *L. major* LeishChr31.

**Supplementary Figure 4: Gene disposition and orientation in Trypanosomatids chrs.** For each clade, chromosomes are drawn in scale. Genes are represented by black boxes and their orientation by the presence above (plus strand) or below (minus strand) of the chromosomal line.

**Supplementary Figure 5: Codon usage in *L. donovani* genes in each chr.** Comparison of Measure Independent of Length and Composition (MILC) in Chr 31 (red) and all other chromosomes combined (black), represented as **A)** density plots, **B)** boxplot and violin plots. **C)** Comparison of MILC (Y axis) and gene length (X axis), **D)** Log10 MILC in each chromosome. **E)** Heatmap of the codon relative presence in genes from each chromosome.

**Supplementary Figure 6: LeishChr31 gene expression and copies in all seven clones.** Top panel: Correlation between DNA AARD (X axis) and RNA AARD (Y axis) for each SNP position from the tetraploid LeishChr31. Each dot represents one SNP position from the LeishChr31. Correlation in clone 3 (C3) (r=0.872, p-value 0); C4 (r=0.875, p-value 0); C6 (r=0.868, p-value 0); C7 (r=0.869, p-value 0); C8 (r=0.905, p-value 0); C9 (r=0.866, p-value 0) and C10 (r=0.887, p-value 0). Bottom panel: DNA (red line) and RNA (blue line) AARD along LeishChr31 in C3.

**Supplementary Table 1:** Overview of the WGS read libraries used in the CCNV estimations. Including read counts, proportion of used reads, genome coverage (columns 1-7) and the estimate of somy for each chr/scaffold (columns 8 - last column, numbers differ based on the number of chromosomes in the clade). Each clade is represented in a different tab.

**Supplementary Table 2:** Origin of each predicted proteome used in the phylogeny estimations.

**Supplementary Table 3:** Comparison of the WGS and RNAseq coverage of each chromosome for each clone/replicate. Sample= Clone ID; SRA_DNA= SRA ID from the WGS sample; SRA_RNA = SRA ID from the RNAseq sample; Chr=Chromosome; DNA = Mean chr coverage using WGS data; RNA = mean chr coverage based on RNAseq data.

**Supplementary Table 4:** Comparison of population wide CCNV, represented by IGE and IAL.

**Supplementary Table 5:** Comparison of population wide CCNV in each individual chromosome, using CDO, Median Chr copies and CSV.

**Supplementary Table 6:** Number and density of segregating sites in each chr in *Leishmania, C. bombi* and *Leptomonas* populations.

**Supplementary Table 7:** Genes with highest nucleotide diversity in Chr31. Each clade is represented in a different tab.

**Supplementary Table 8:** Summary of GO enrichment results. Each dataset is represented in a different tab. Proportion = Proportion of the genes with a given GO that were present in the Duplicated31 in the clade.

## Acknowledgments

This project was undertaken on the Viking Cluster, which is a high-performance compute facility provided by the University of York. We are grateful for computational support from the University of York High Performance Computing service, Viking, and the Research Computing team. We state that no conflicts of interest exist.

We thank funding agencies that provided funds for this study, Medical Research Council (MRC), Marie Sklodowska-Curie, Wellcome Trust, Newton UK:Brazil Joint Centre Partnership in Leishmaniasis, Brazilian Federal Agency for Support of Graduate Education (CAPES), Brazilian Council for Scientific and Technological Development (CNPq), Minas Gerais State Agency for Research and Development (FAPEMIG), São Paulo State Agency for Research and Development (FAPESP), National Institute for Science and Technology in Vaccines (INCTV) and Pró-reitoria de Pesquisa, Universidade Federal de Minas Gerais. J.L.R.-C. and D.C.J. are supported by a MRC New Investigator Research Grant (MR/T016019/1) and by MRC Newton as a component of the UK:Brazil Joint Centre Partnership in Leishmaniasis. A.C.-d.-S., S.A.P.C., and L.A.V. received scholarships from CNPq. JB is supported by a FAPESP post-doctoral fellowship (20/01883-7). R. M., J.D and C. A. M are supported by the Wellcome Trust (Investigator Award (224501/Z/21/Z) to R.M and by the BBSRC (BB/N016165/1 and BB/R017166/1 to R.M, and to BB/W001101/1 to R.M. and C.A.M). J.D was supported by a Marie Sklodowska-Curie Individual Fellowship, RECREPEMLE.

