## Supplementary figure 1 for "Aneuploidies are an ancestral feature of trypanosomatids, and an ancient chromosome duplication is maintained in extant species"

**Supplementary Figure 1:** Heatmap (top panel) and density plot (bottom panel) of chromosomal copies in all isolates from all clades. The heatmaps show the sample-by-sample CCNV. Each line corresponds to a different chromosome/scaffold and each column to a different isolate. The chromosome copies are represented by a colour scale from dark-blue (low copy number) to dark-red (high copy number), where euploid chromosomes (~1 copy for each haploid genome copy) are in light blue. The dendrogram on top clusters the samples by UPGMA, based on the Manhattan distance of their chromosome copy numbers. The density plots represent the deviation from euploidy for each chromosome. Each box corresponds to a given chromosome/scaffold, the X axis to the chromosome somy and the Y axis to the frequency of observations. The red dotted line represents the expected value for euploid chromosomes. Each page corresponds to one clade, on the order: *C. bombi*, *C. fasciculata*, *Endotrypanum*, *L. donovani*, *L. major*, *Leptomonas*, *Paratrypanosoma*, *Porcisia*, *T. brucei*, *T. congolense*, *T. cruzi*, *T. vivax*.

### *C. bombi*

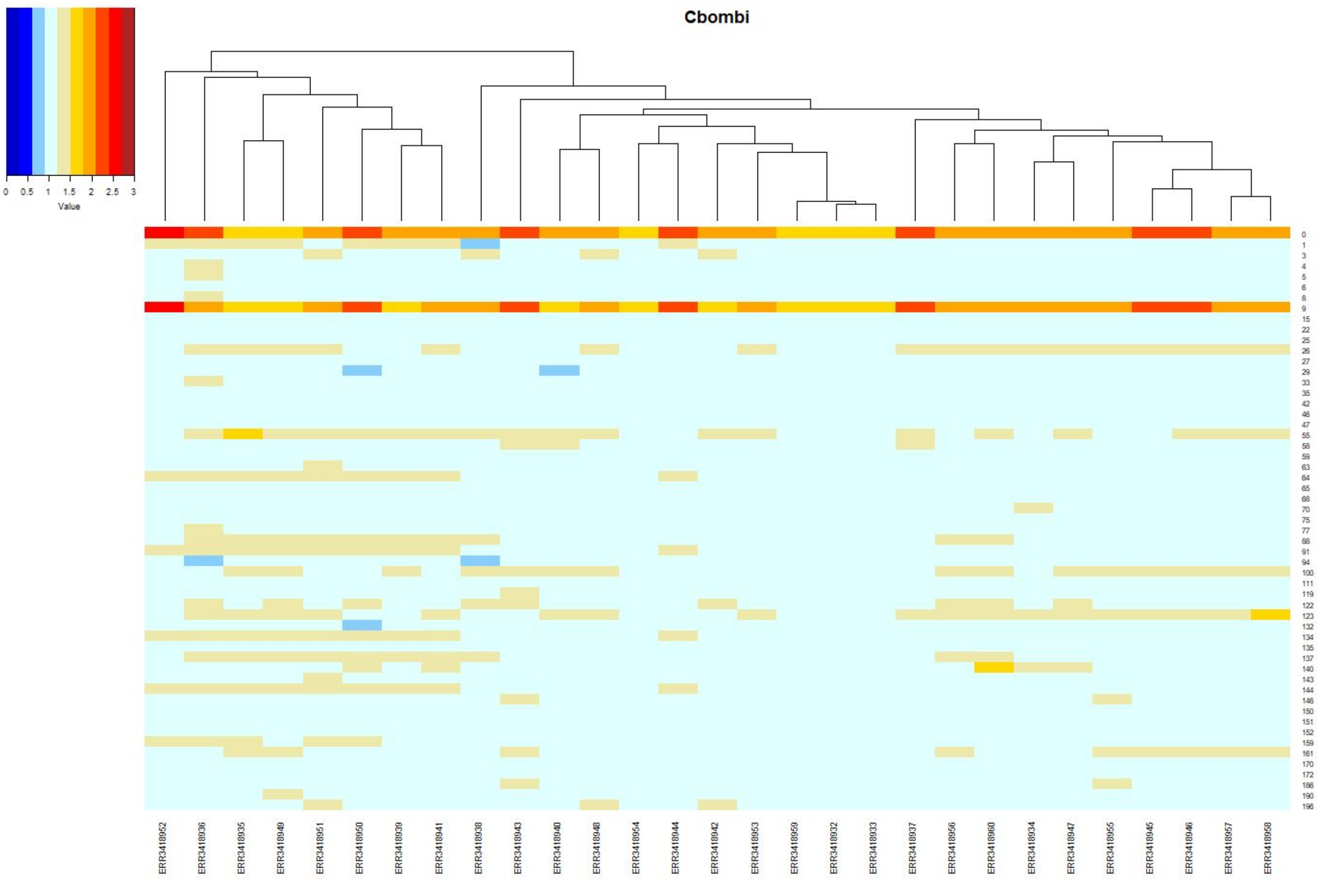

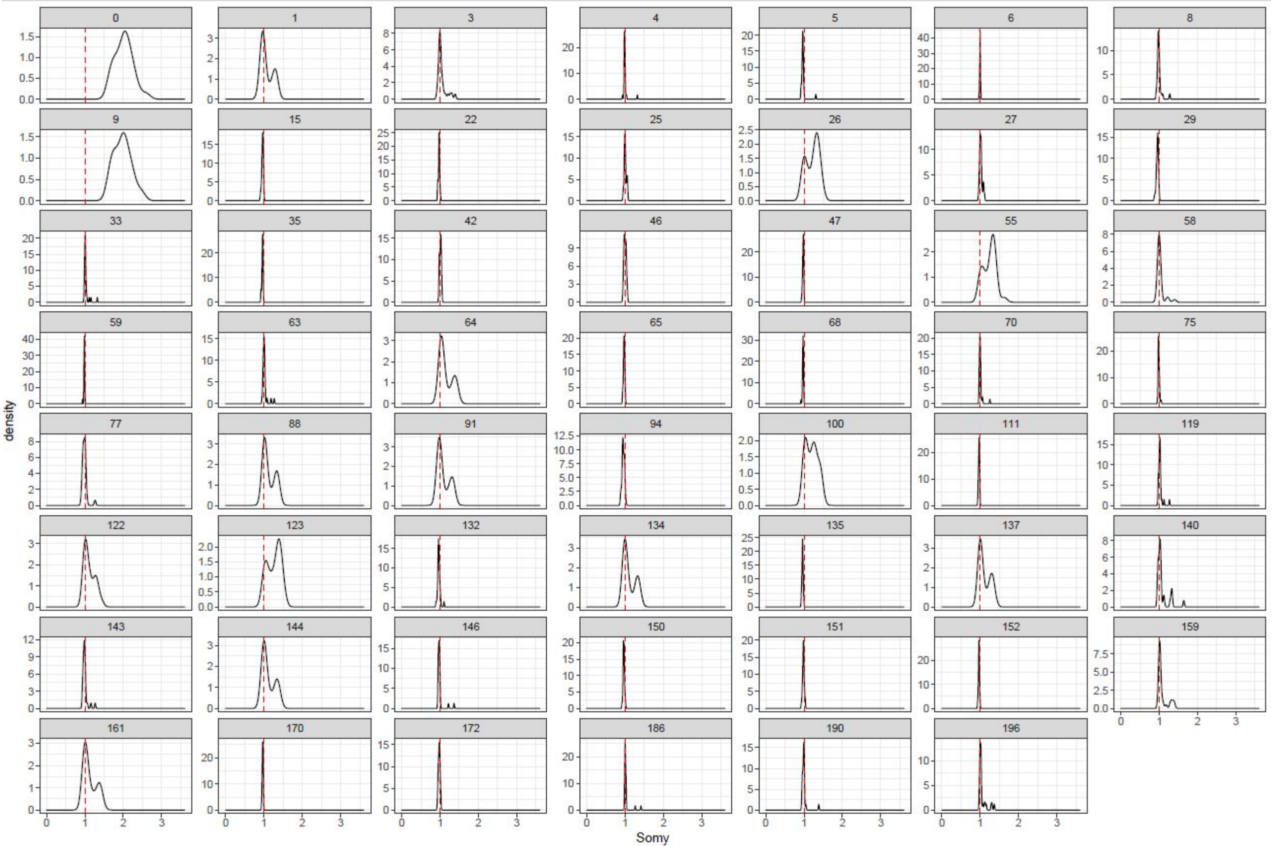

### *C. fasciculata*

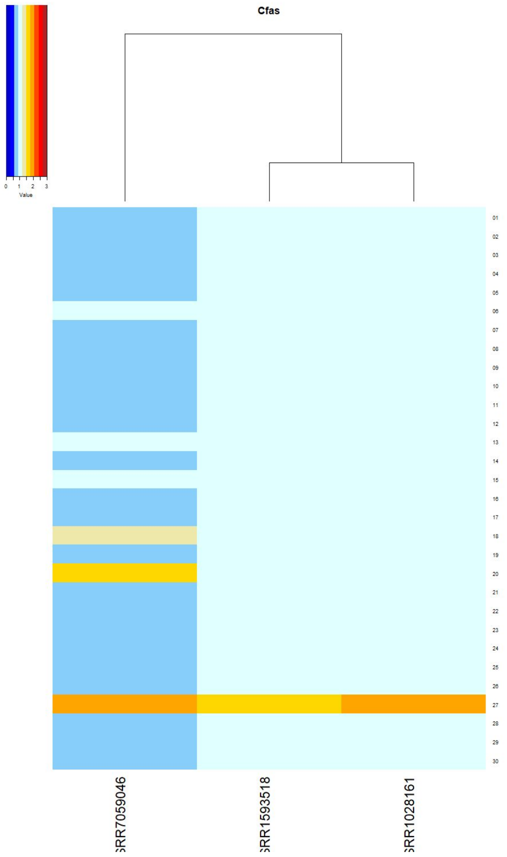

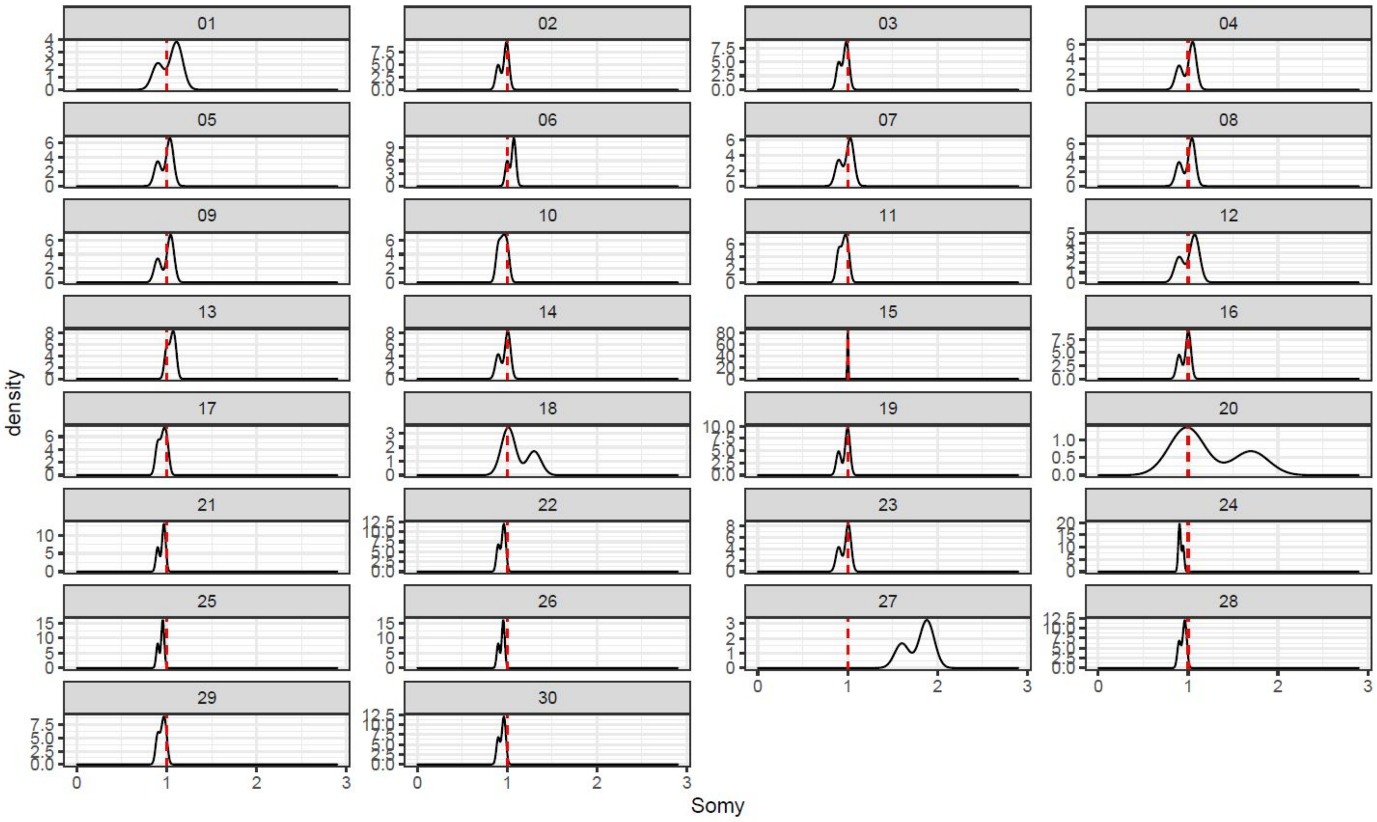

*Endotrypanum*

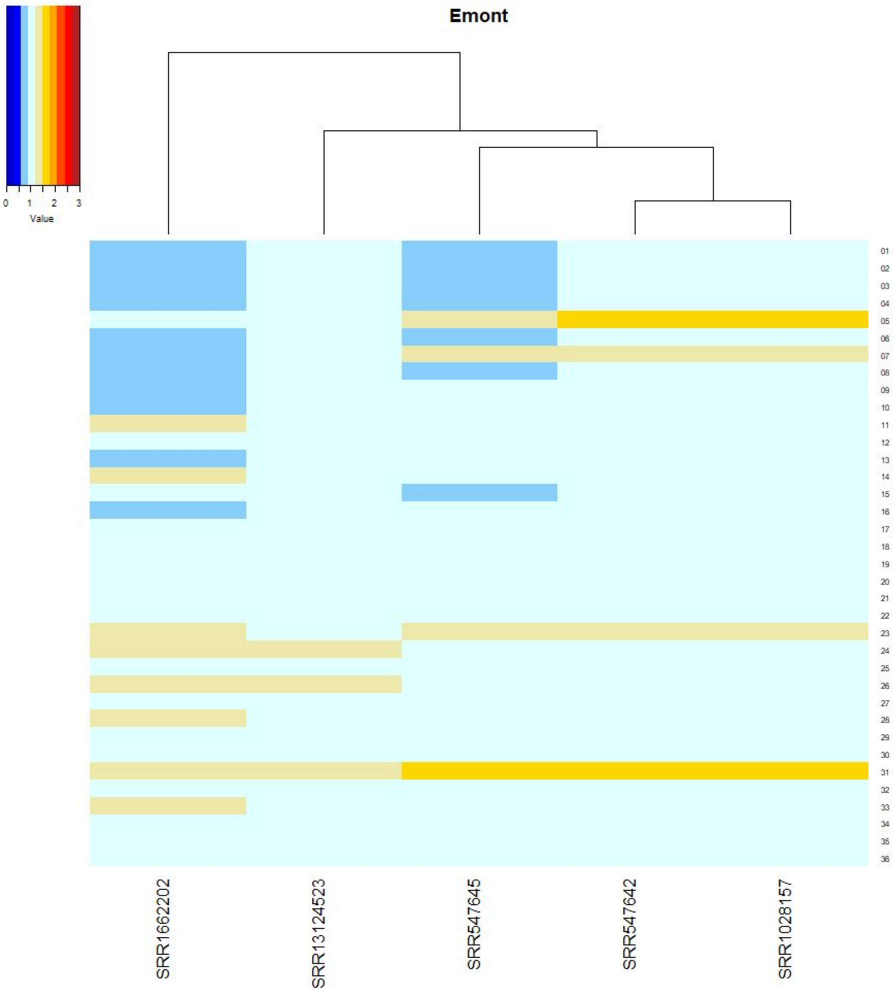

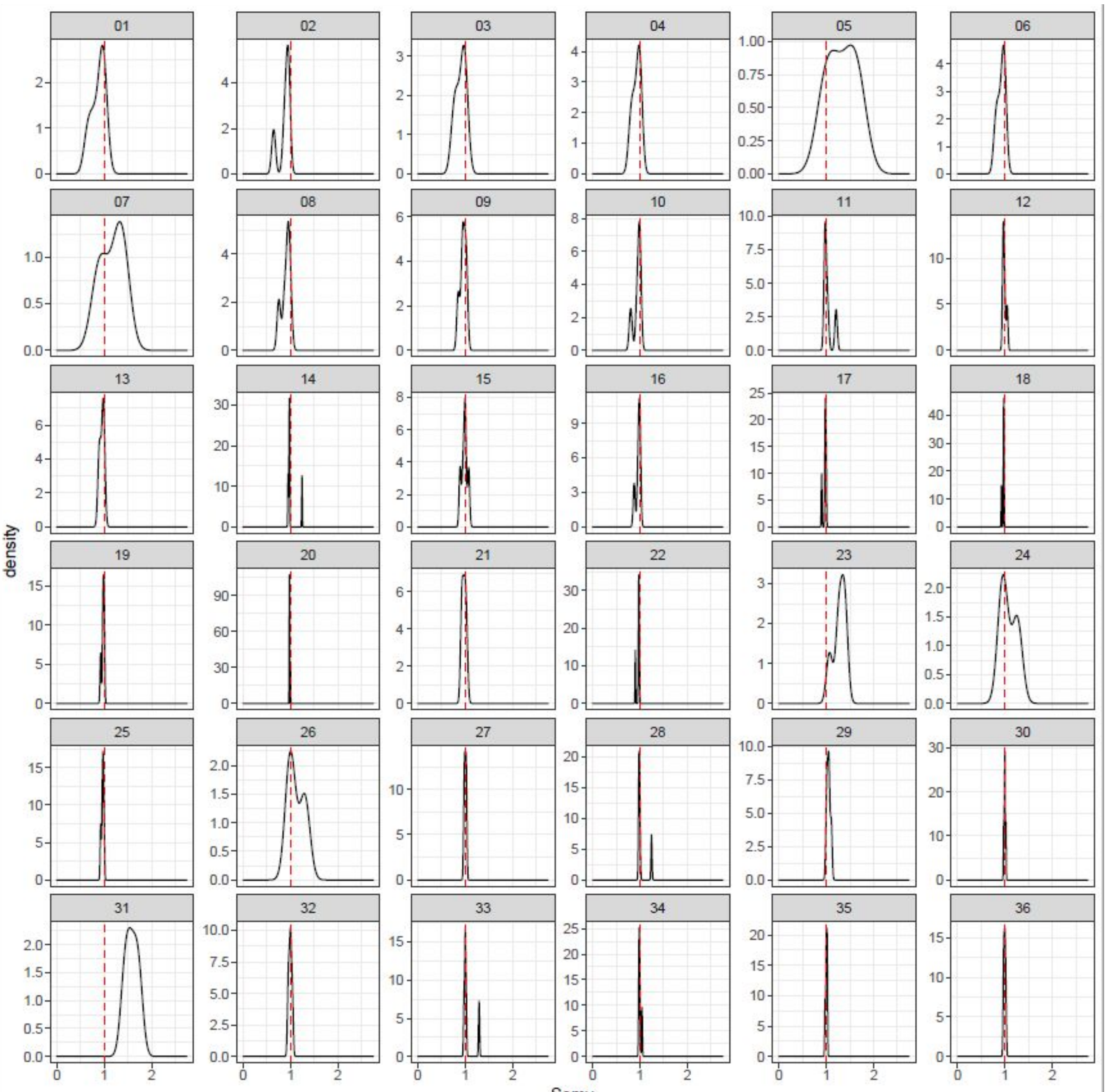

*L. donovani*

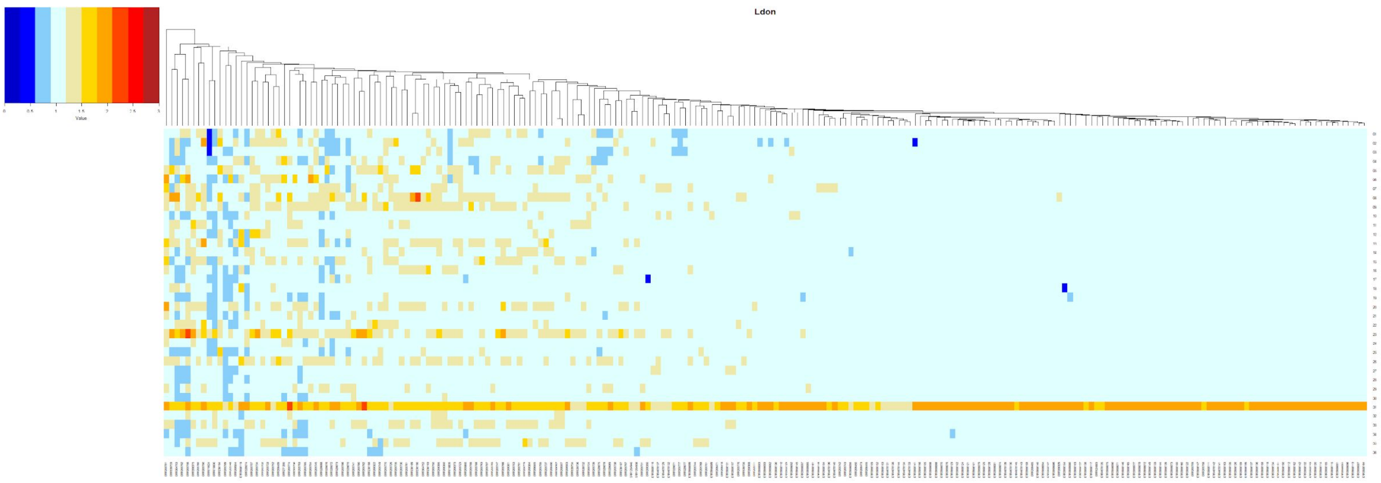
Ldon

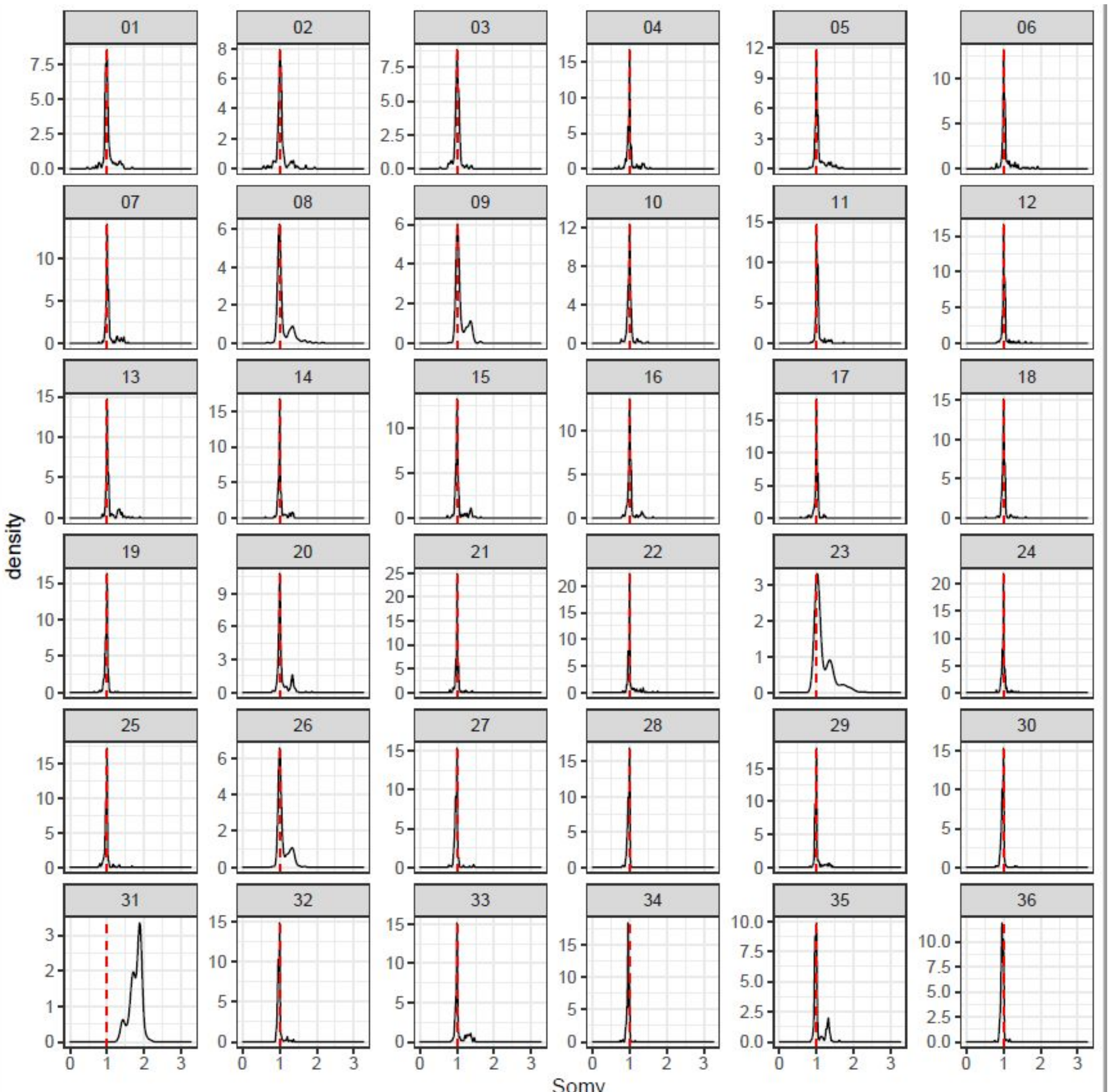
density
Somy

*L. major*

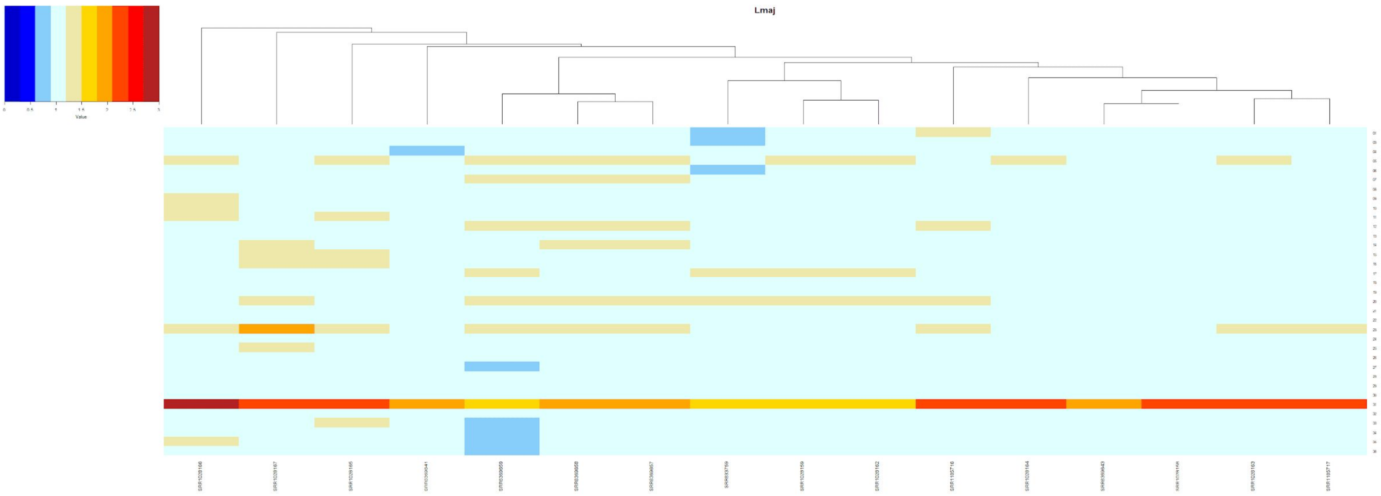
Lmaj
Value

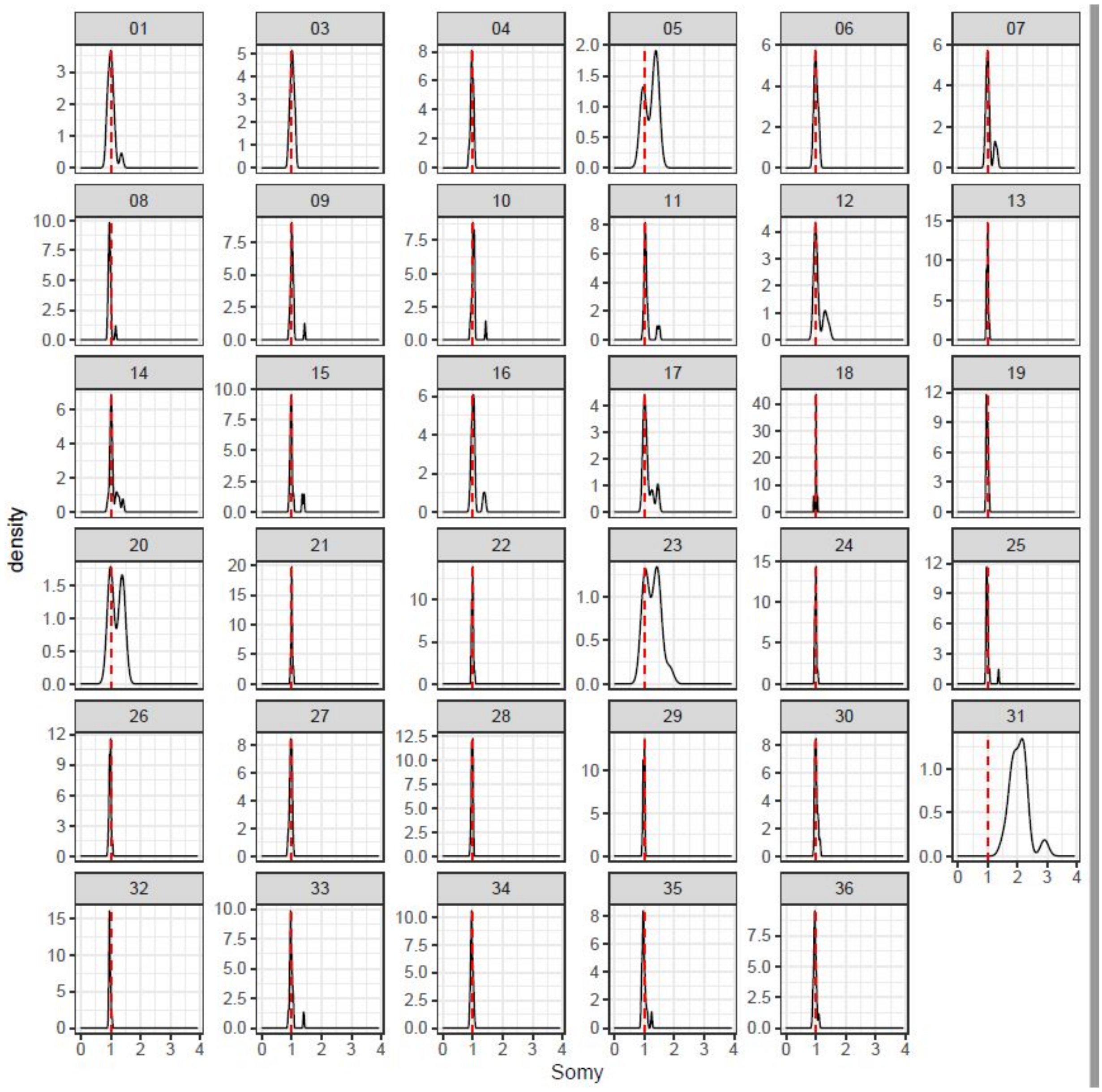
01
density
Somy

### *Leptomonas*

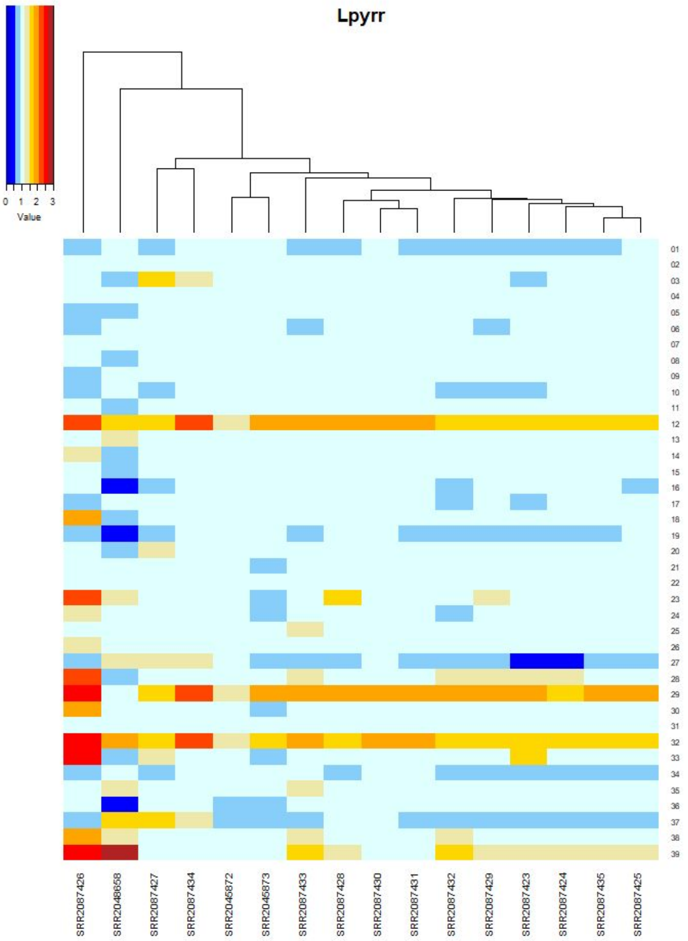
Lpyrr
0 1 2 3
Value
01 02 03 04 05 06 07 08 09 10 11 12 13 14 15 16 17 18 19 20 21 22 23 24 25 26 27 28 29 30 31 32 33 34 35 36 37 38 39
SRR2087426
SRR2048658
SRR2087427
SRR2087434
SRR2045872
SRR2045873
SRR2087433
SRR2087428
SRR2087430
SRR2087431
SRR2087432
SRR2087429
SRR2087423
SRR2087424
SRR2087435
SRR2087425

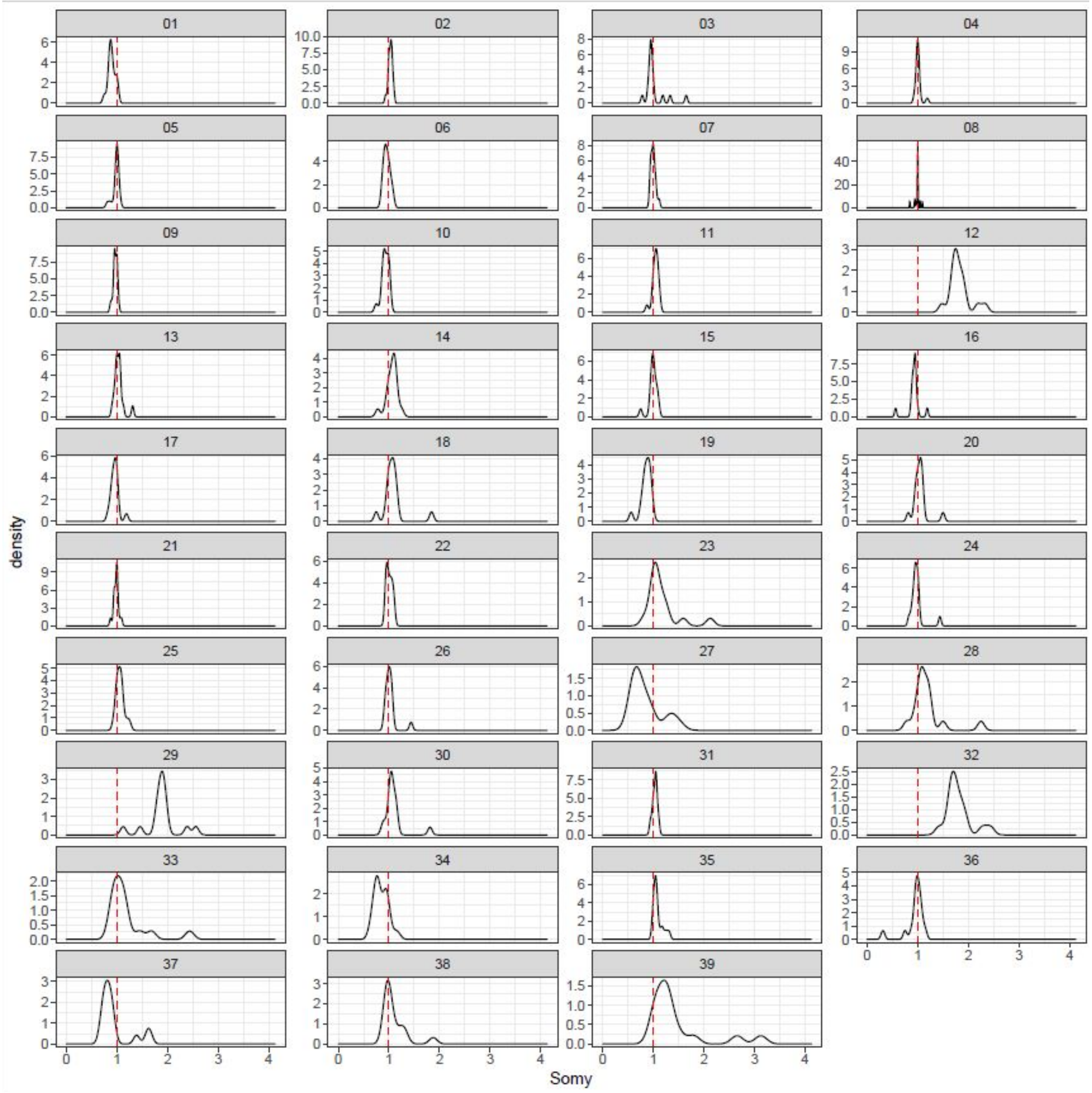
01 02 03 04 05 06 07 08 09 10 11 12 13 14 15 16 17 18 19 20 21 22 23 24 25 26 27 28 29 30 31 32 33 34 35 36 37 38 39
density
Somy
0 1 2 3 4

*Paratrypanosoma*

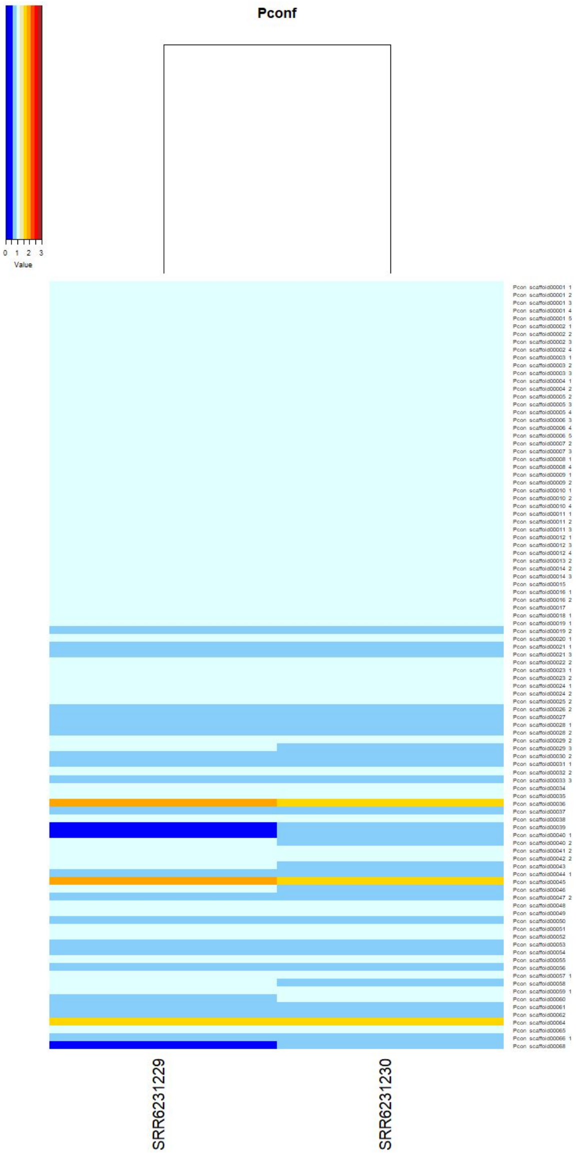

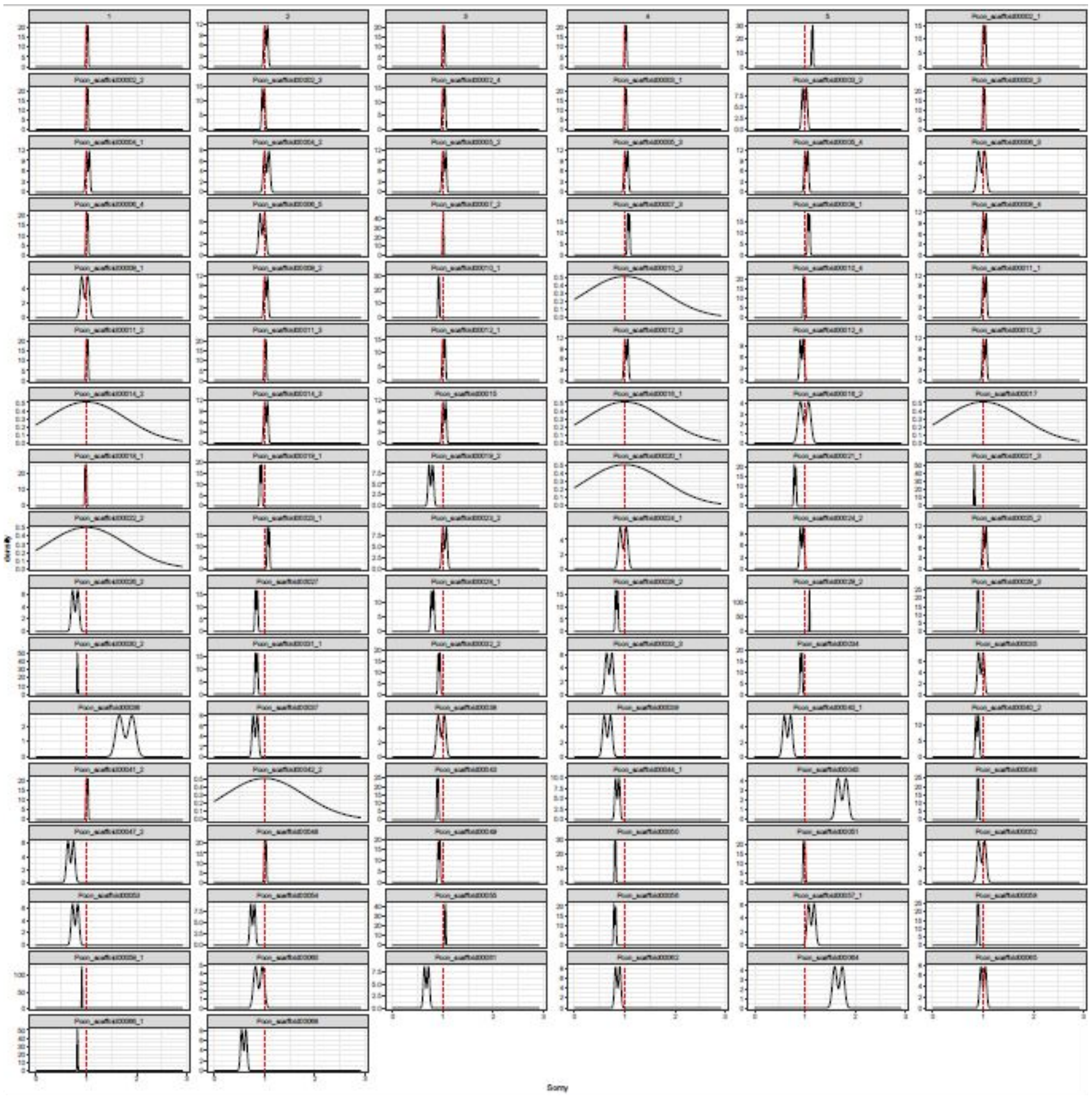

*Porcisia*

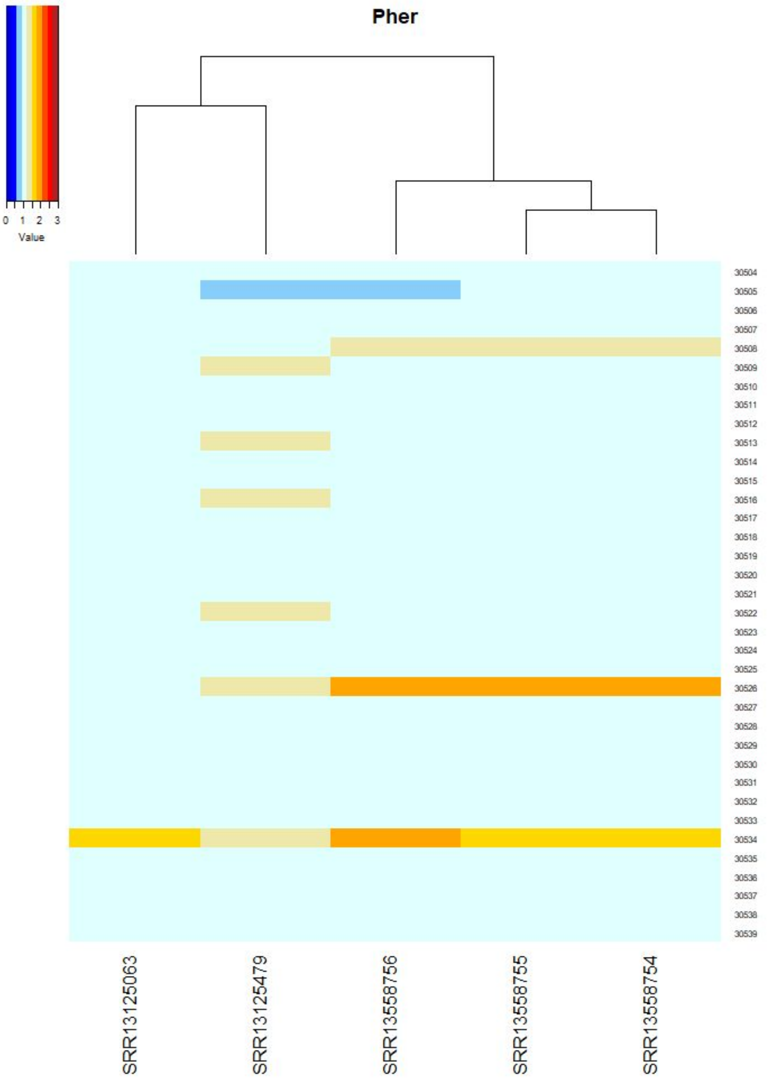
Pher
Value
0 1 2 3
SRR13125063
SRR13125479
SRR13558756
SRR13558755
SRR13558754

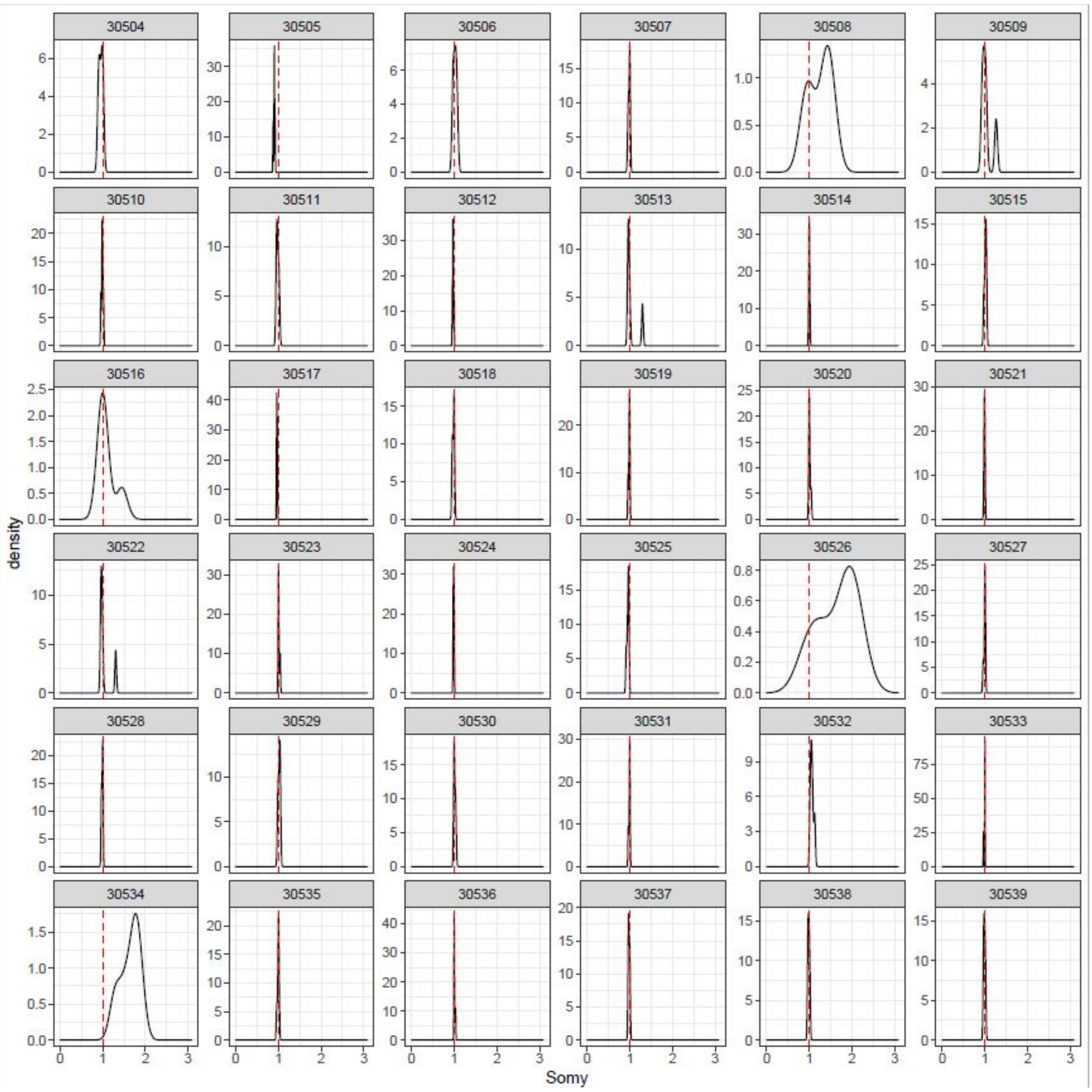
density
Somy

*T. brucei*

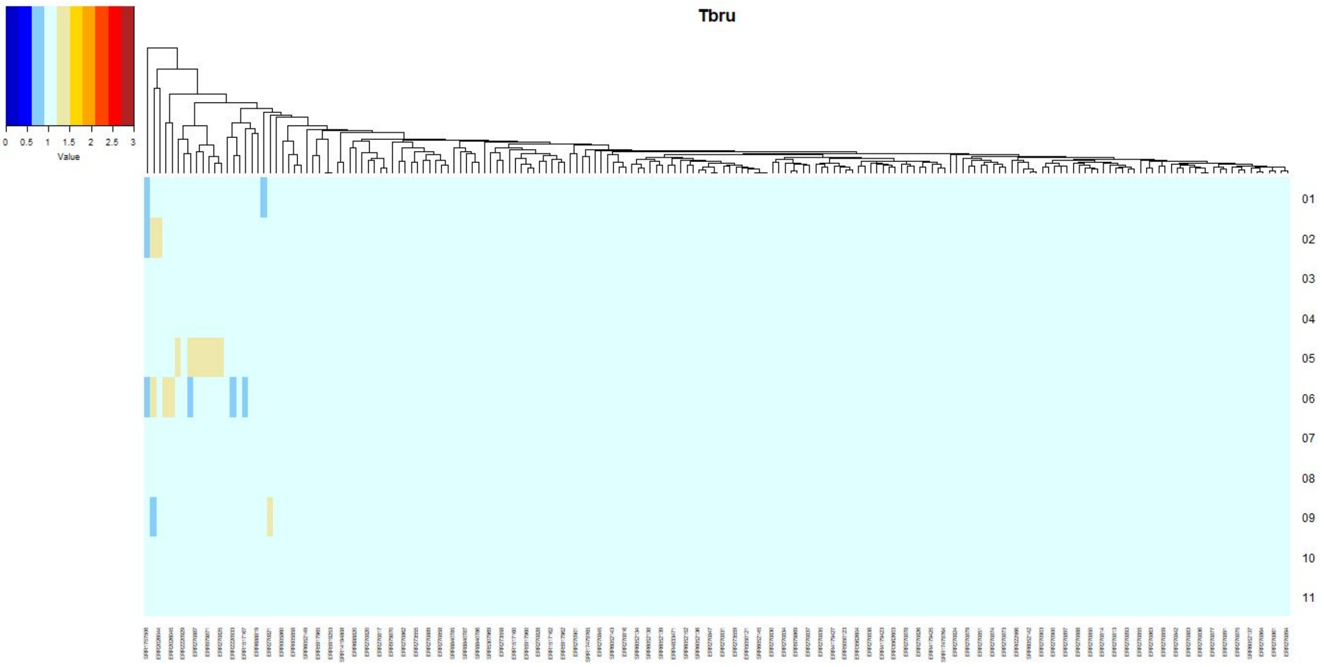
Tbru
0 0.5 1 1.5 2 2.5 3
Value
01
02
03
04
05
06
07
08
09
10
11

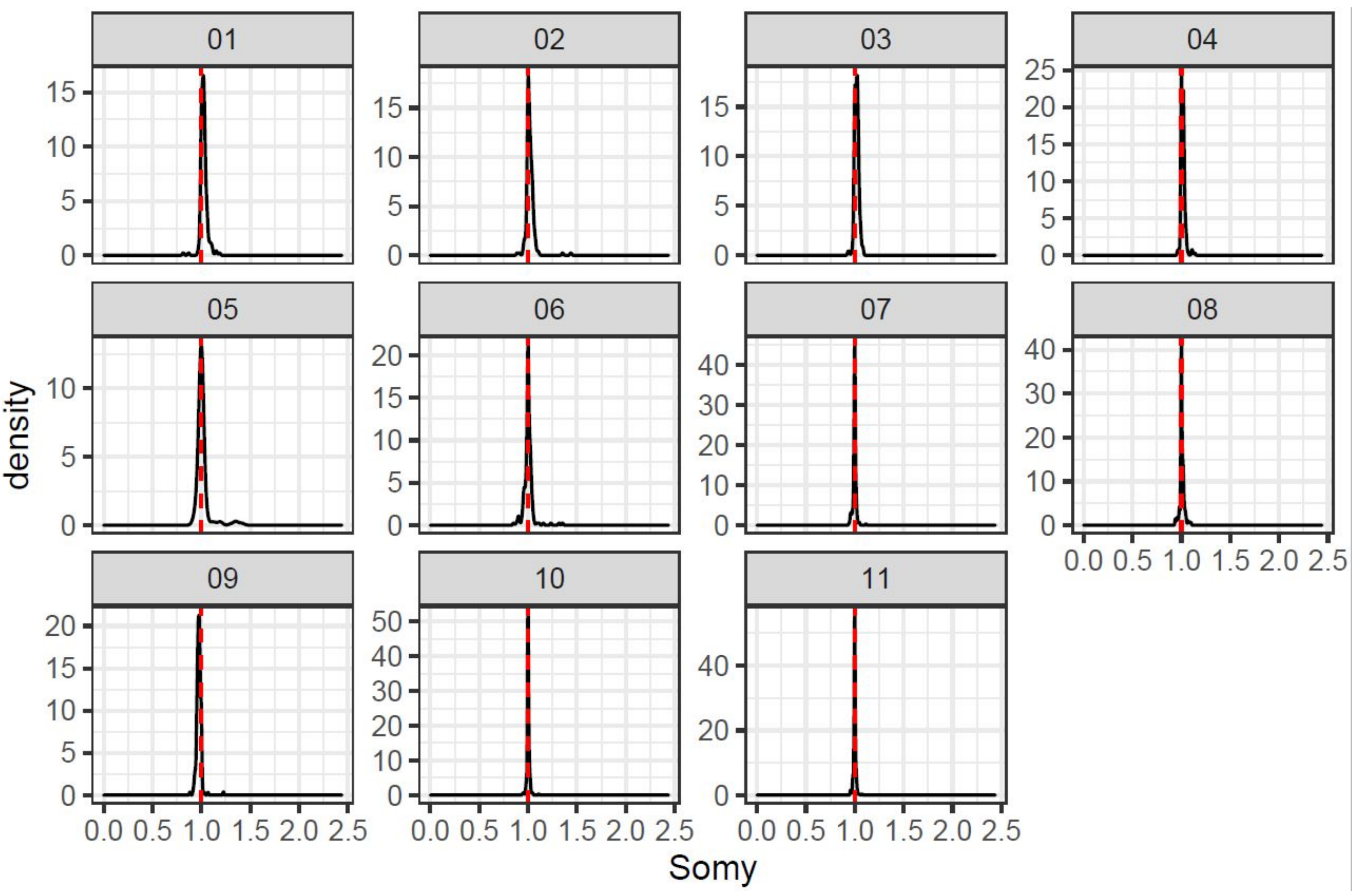
01
02
03
04
05
06
07
08
09
10
11
density
Somy
0.0 0.5 1.0 1.5 2.0 2.5

*T. congolense*

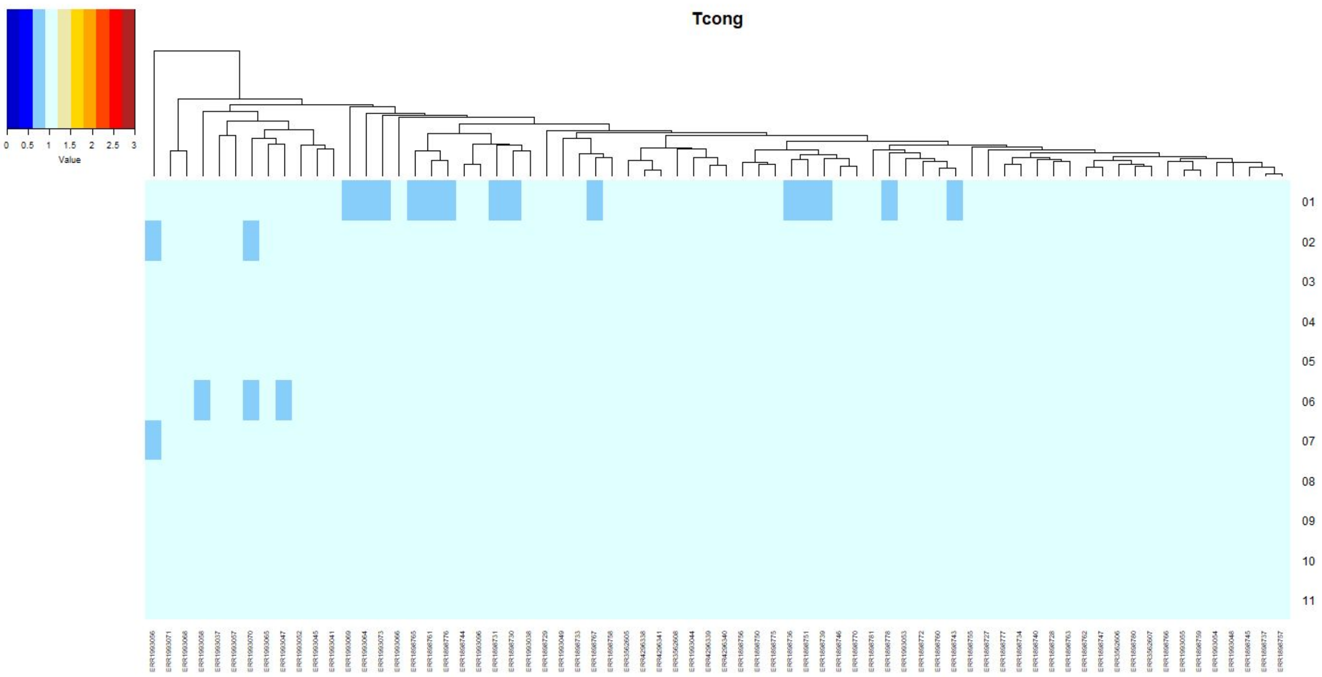

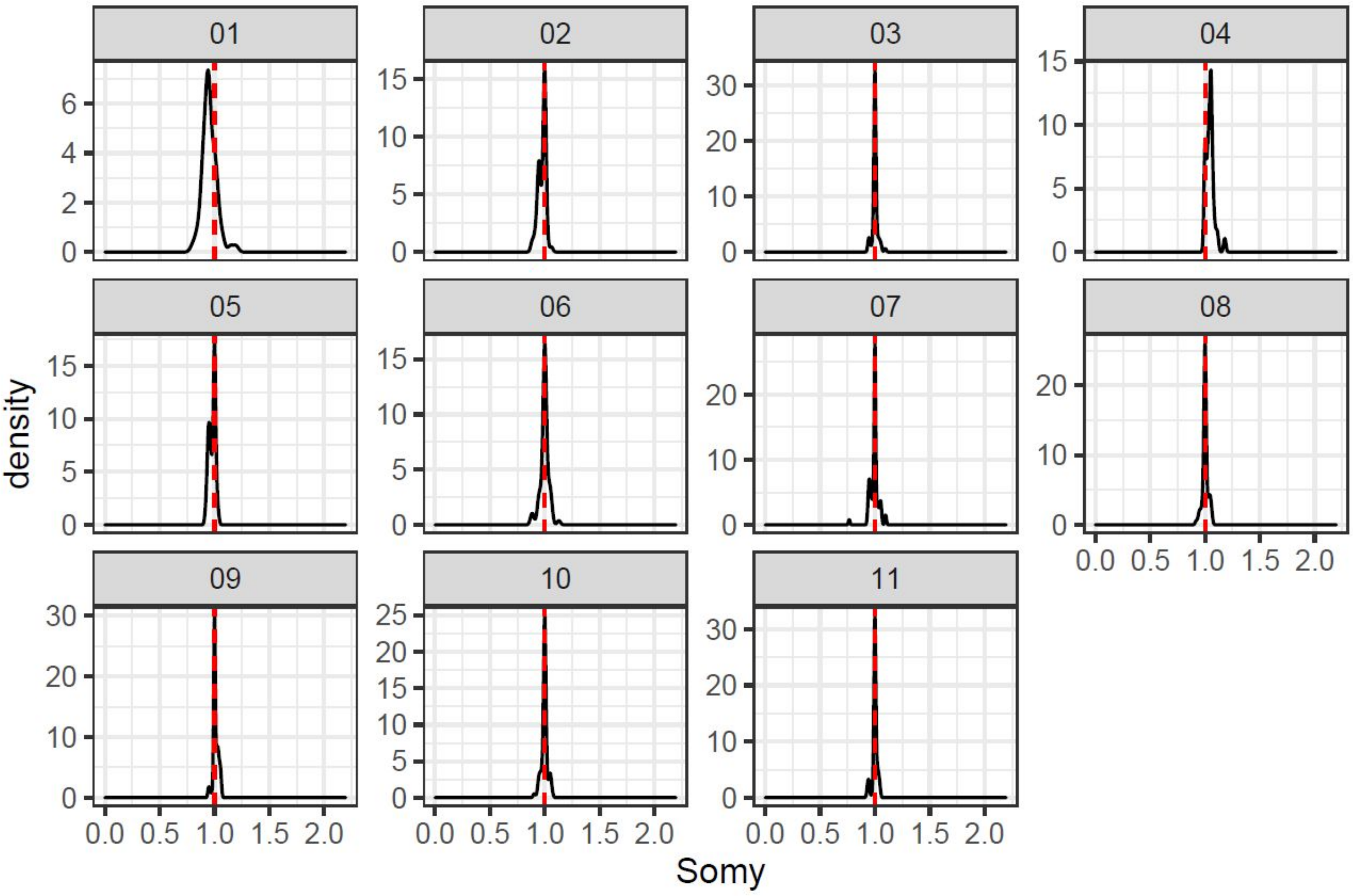

*T. cruzi*

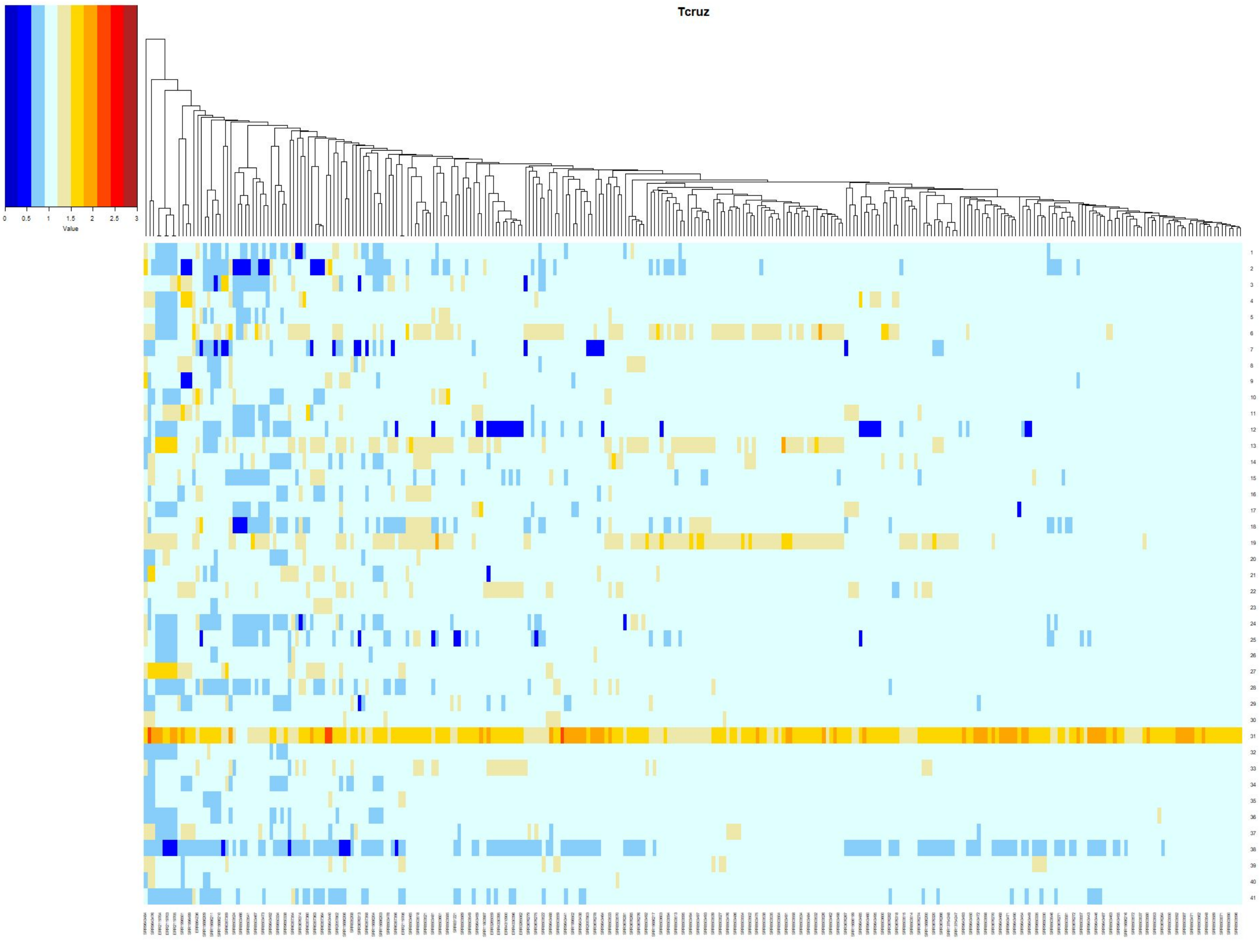

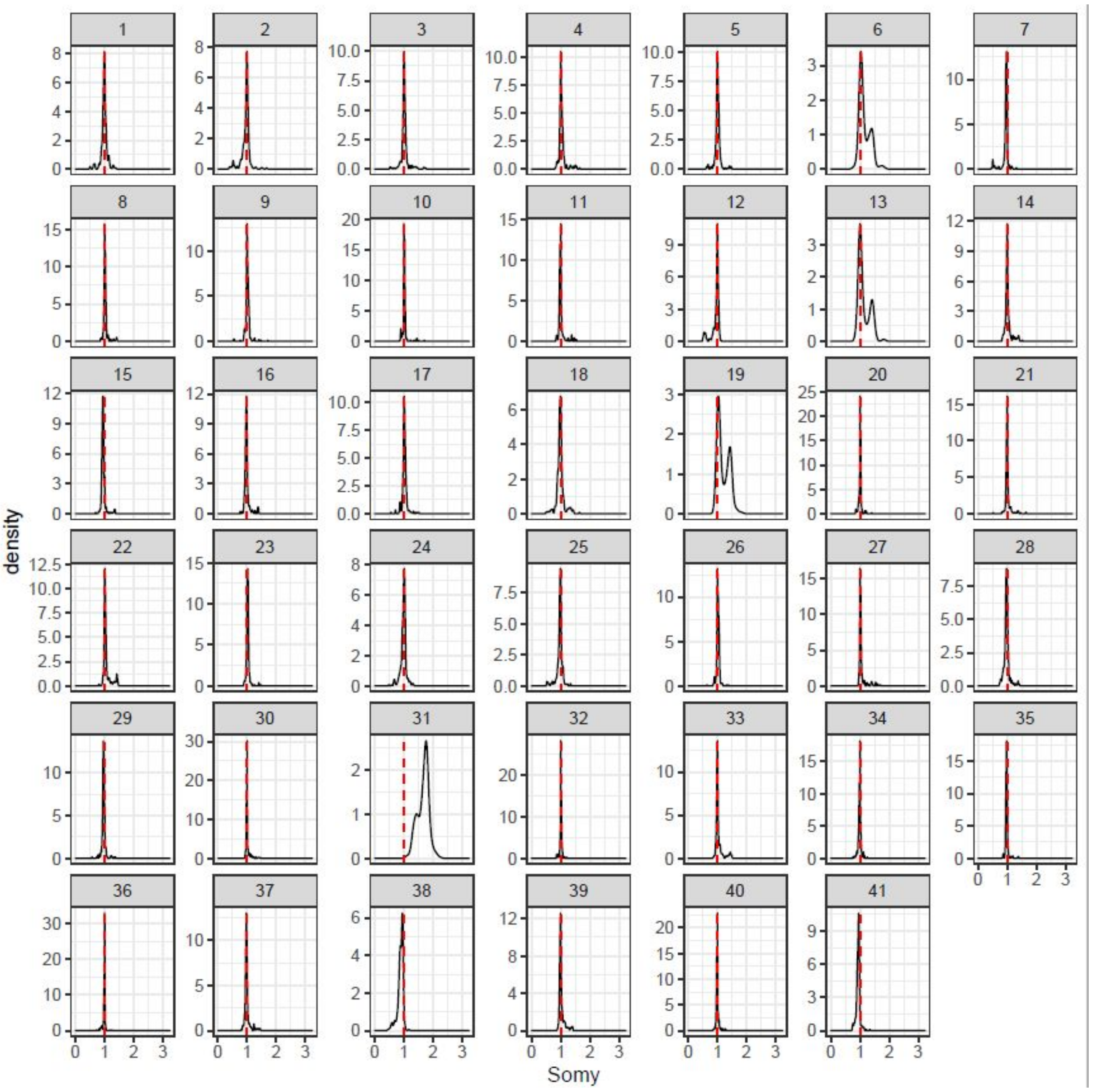

*T. vivax*

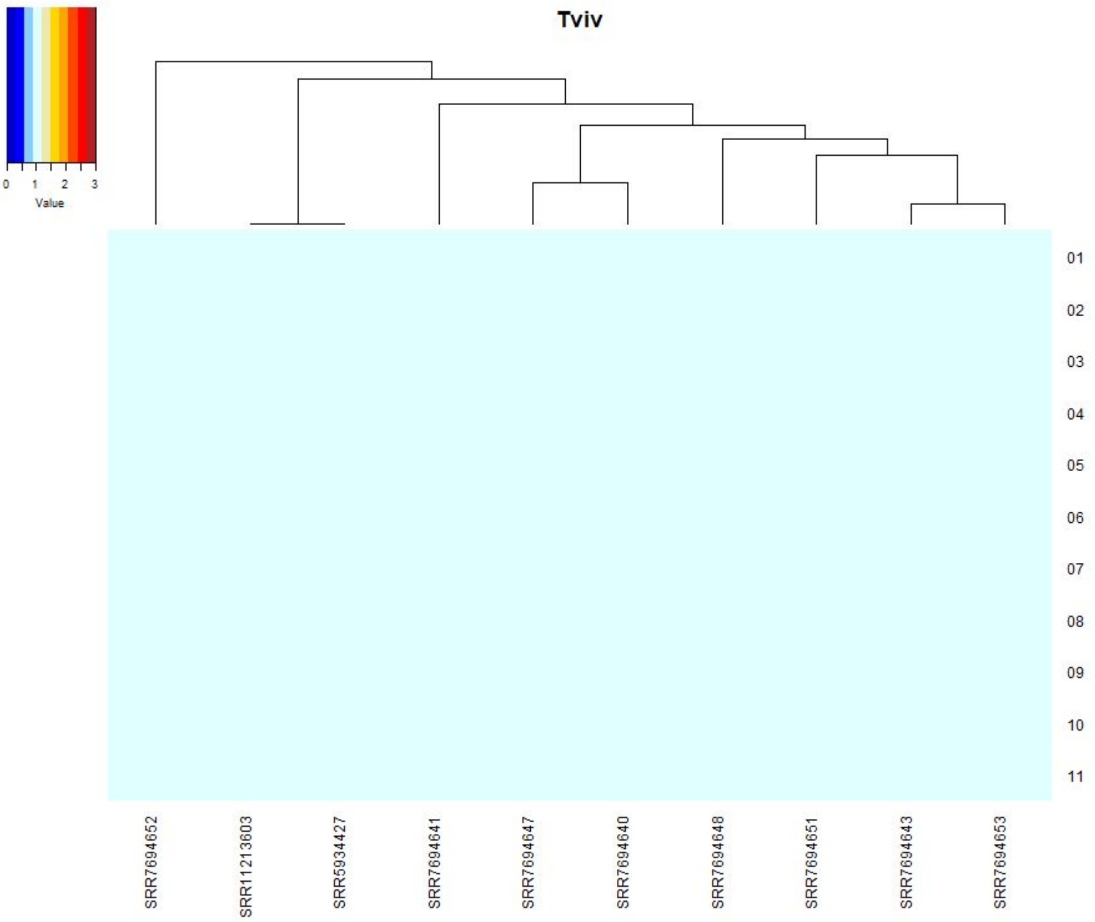
Tviv
0 1 2 3
Value
01
02
03
04
05
06
07
08
09
10
11
SRR7694652
SRR11213603
SRR5934427
SRR7694641
SRR7694647
SRR7694640
SRR7694648
SRR7694651
SRR7694643
SRR7694653

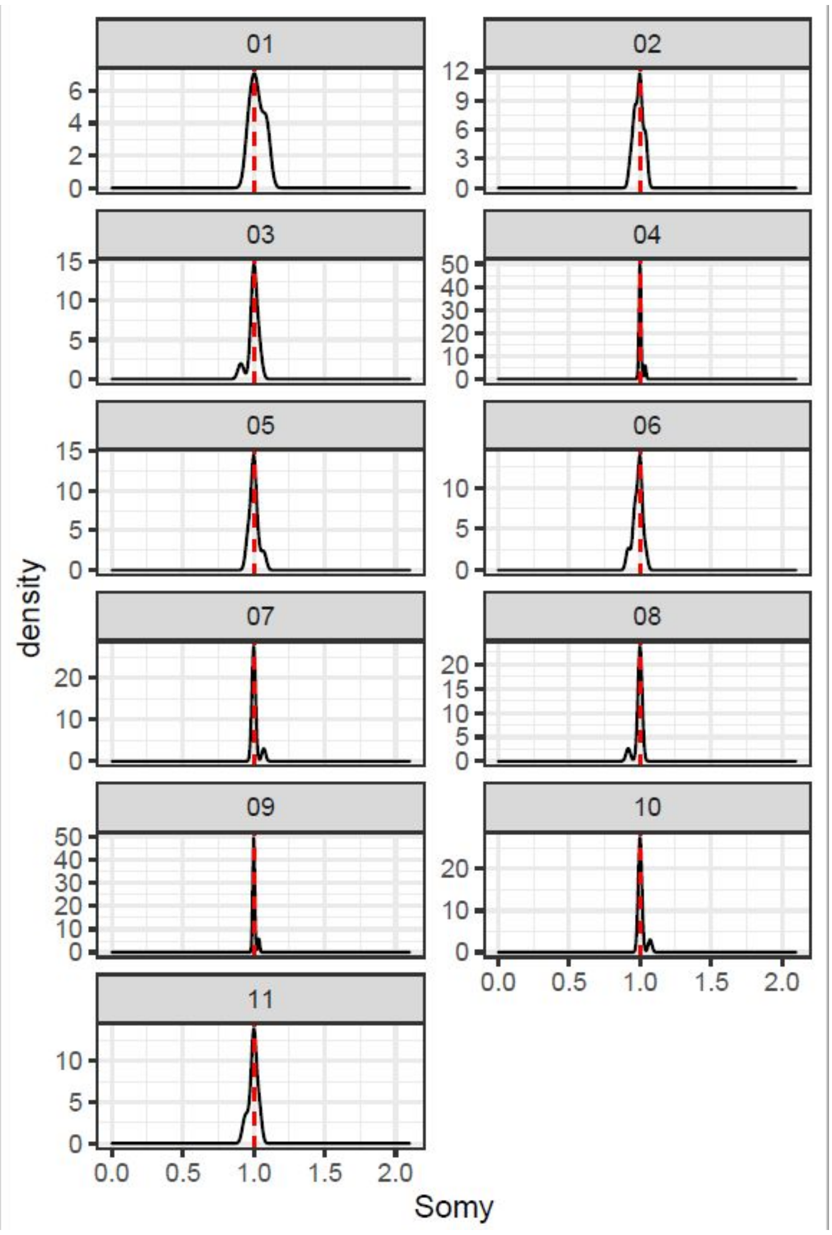
01
02
03
04
05
06
07
08
09
10
11
density
0.0 0.5 1.0 1.5 2.0
Somy
