## Supplementary figure 2 for "Aneuploidies are an ancestral feature of trypanosomatids, and an ancient chromosome duplication is maintained in extant species"

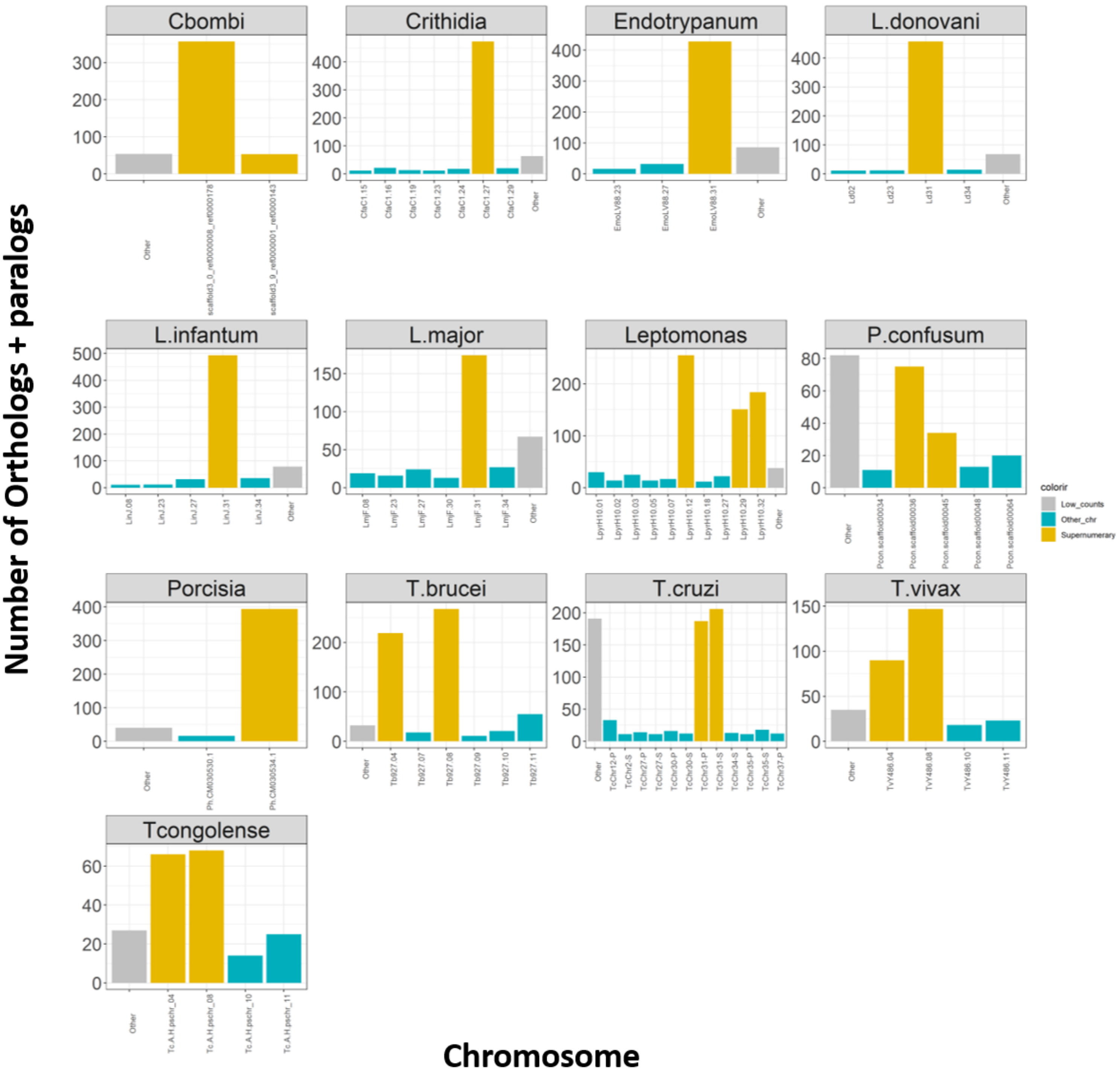


**Supplementary Figure 2: Proportion of gene sharing between LeishChr31 and the chromosomes in other species.** Each panel corresponds to a different species. The X axis corresponds to the number of shared Orthologs + Paralogs in a given chromosome and the *L. major* LeishChr31. Chromosomes with consistent extra copies and syntenic to L. major LeishChr31 are highlighted in gold, while other chromosomes are represented in blue. Chromosomes/scaffolds with less than 10 sharings were “grouped” in the “others” column, in grey.
