## Supplementary figure 3 for "Aneuploidies are an ancestral feature of trypanosomatids, and an ancient chromosome duplication is maintained in extant species"

**A**

| Species | Chromosome | Chr_genes | Ortologs | Lmaj_ortologs | Prop_Lmajor31 | Prop_Other31 |
|---|---|---|---|---|---|---|
| ***L.donovani*** | Ld31 | 332 | 303 | 321 | 0.911931818 | 0.912650602 |
| ***L.pyrrhocoris*** | LpyrH10.12 - LpyrH10.29 - LpyrH10.32 | 502 | 353 | 312 | 0.886363636 | 0.703187251 |
| ***P.hertigi*** | CM030534.1 | 295 | 284 | 282 | 0.801136364 | 0.962711864 |
| ***E.monterogeii*** | EmoLV88.31 | 303 | 283 | 282 | 0.801136364 | 0.933993399 |
| ***Cbombi*** | 0 - 9 | 490 | 295 | 282 | 0.801136364 | 0.602040816 |
| ***C.fasciculata*** | CfaC1.27 | 522 | 284 | 253 | 0.71875 | 0.544061303 |
| ***T.cruzi*** | TcChr31-S | 268 | 140 | 128 | 0.363636364 | 0.52238806 |
| ***P.confusun*** | Pcon.scaffold00036 - Pcon.scaffold00045 | 113 | 74 | 79 | 0.224431818 | 0.654867257 |
| ***T.brucei*** | Tb927.04 AND Tb927.08 | 1440 | 141 | 66 | 0.1875 | 0.097916667 |
| ***T.vivax*** | TvY486.04 AND TvY486.08 | 1247 | 49 | 31 | 0.088068182 | 0.039294306 |
| ***T.congolense*** | Tc.A.H.pschr_04 AND Tc.A.H.pschr_08 | 1421 | 12 | 4 | 0.011363636 | 0.008444757 |

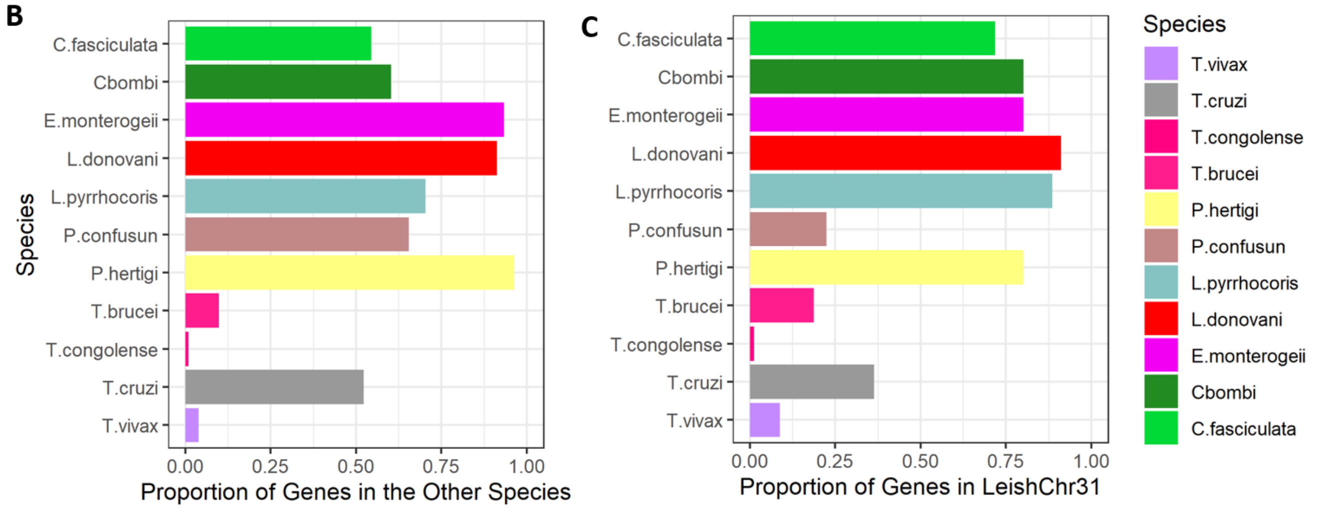


**Figure Supporting 3: Proportion of gene sharing between LeishChr31 and the chromosomes in other species. A)** Table representing the number of genes (total genes in *L major* chr 31 = 352) in each LeishChr31 syntenic chromosome and how many of these genes have orthologs/Paralogs in L. major LeishChr31. **Chr_genes**: Number of genes in a given chromosome (or combination of chromosomes, as in Leptomonas). **Orthologs**: How many of these genes have orthologs/Paralogs in *L. major* LeishChr31. **L. major Orthologs**: How many genes in *L. major* LeishChr31 have orthologs/paralogs with genes in the other species selected chromosomes. **Prop_Lmajor**: The proportion of genes in *L. major* LeishChr31 have orthologs/paralogs with genes in the other species selected chromosomes, **Prop_Other 31**: The proportion of genes in a given chromosome that have orthologs/Paralogs in *L. major* LeishChr31.
