## Supplementary figure 4 for "Aneuploidies are an ancestral feature of trypanosomatids, and an ancient chromosome duplication is maintained in extant species"

C. fasciculata

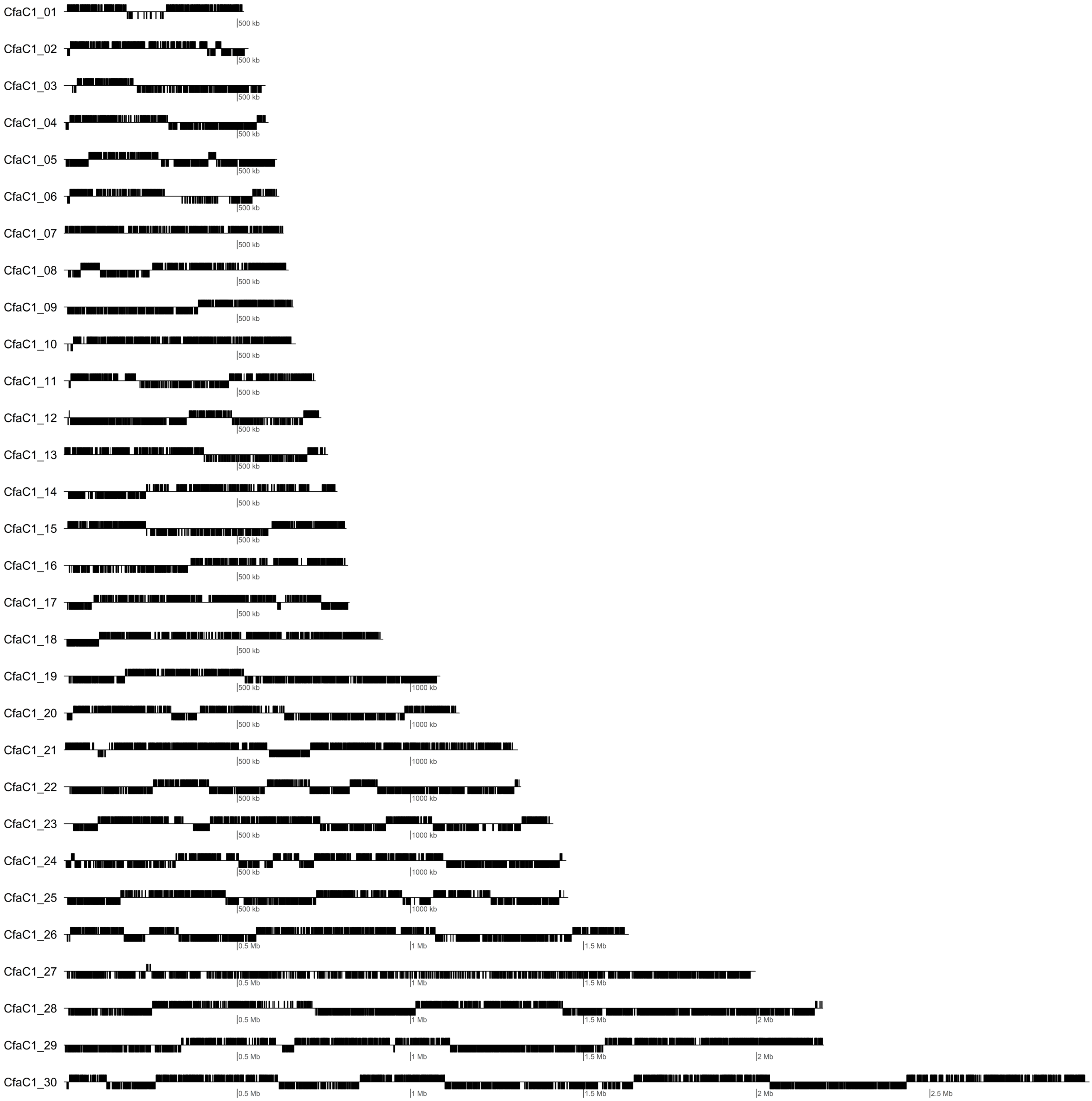

C. bombi

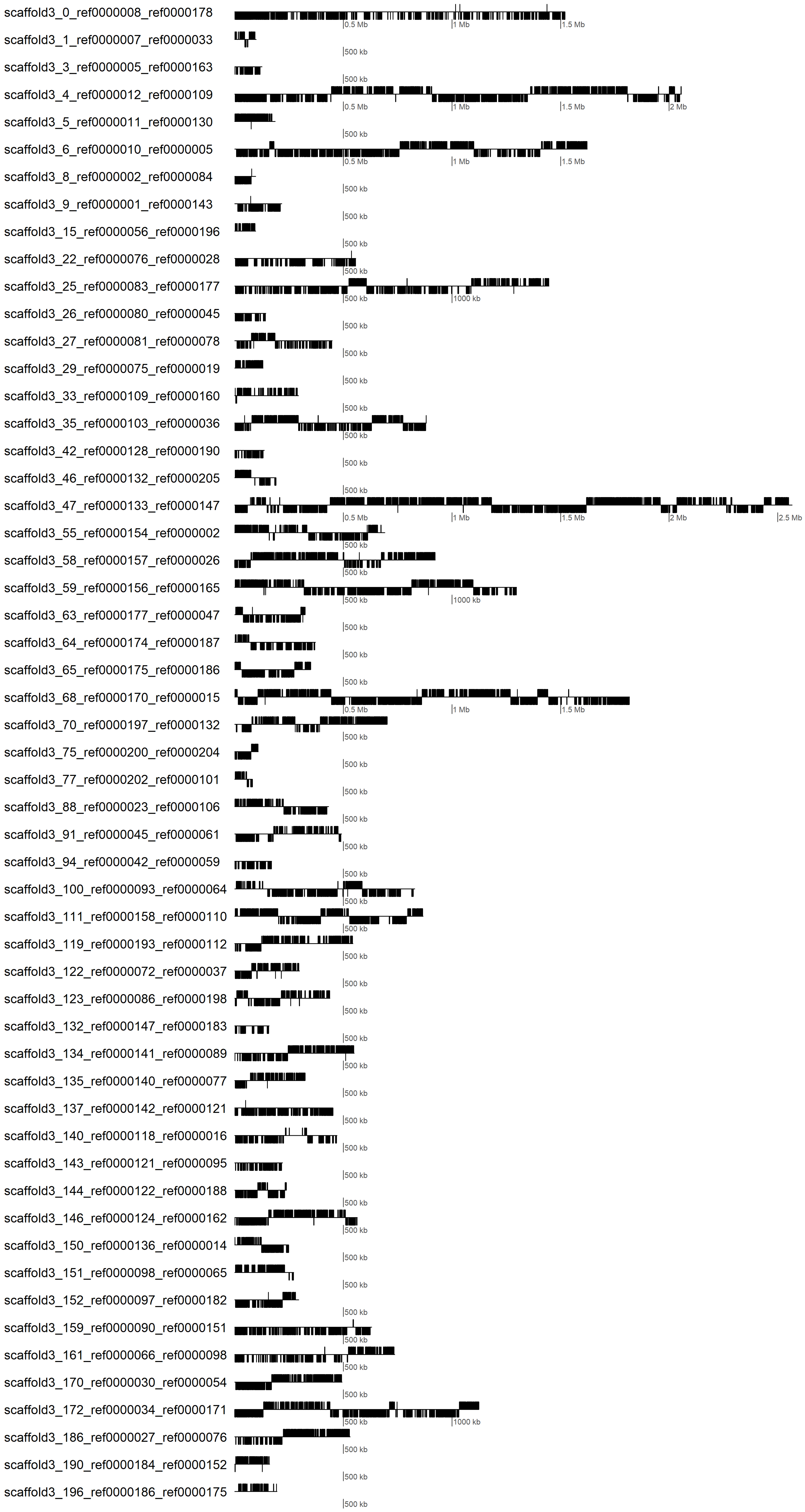
scaffold3_0_ref0000008_ref0000178
scaffold3_1_ref0000007_ref0000033
scaffold3_3_ref0000005_ref0000163
scaffold3_4_ref0000012_ref0000109
scaffold3_5_ref0000011_ref0000130
scaffold3_6_ref0000010_ref0000005
scaffold3_8_ref0000002_ref0000084
scaffold3_9_ref0000001_ref0000143
scaffold3_15_ref0000056_ref0000196
scaffold3_22_ref0000076_ref0000028
scaffold3_25_ref0000083_ref0000177
scaffold3_26_ref0000080_ref0000045
scaffold3_27_ref0000081_ref0000078
scaffold3_29_ref0000075_ref0000019
scaffold3_33_ref0000109_ref0000160
scaffold3_35_ref0000103_ref0000036
scaffold3_42_ref0000128_ref0000190
scaffold3_46_ref0000132_ref0000205
scaffold3_47_ref0000133_ref0000147
scaffold3_55_ref0000154_ref0000002
scaffold3_58_ref0000157_ref0000026
scaffold3_59_ref0000156_ref0000165
scaffold3_63_ref0000177_ref0000047
scaffold3_64_ref0000174_ref0000187
scaffold3_65_ref0000175_ref0000186
scaffold3_68_ref0000170_ref0000015
scaffold3_70_ref0000197_ref0000132
scaffold3_75_ref0000200_ref0000204
scaffold3_77_ref0000202_ref0000101
scaffold3_88_ref0000023_ref0000106
scaffold3_91_ref0000045_ref0000061
scaffold3_94_ref0000042_ref0000059
scaffold3_100_ref0000093_ref0000064
scaffold3_111_ref0000158_ref0000110
scaffold3_119_ref0000193_ref0000112
scaffold3_122_ref0000072_ref0000037
scaffold3_123_ref0000086_ref0000198
scaffold3_132_ref0000147_ref0000183
scaffold3_134_ref0000141_ref0000089
scaffold3_135_ref0000140_ref0000077
scaffold3_137_ref0000142_ref0000121
scaffold3_140_ref0000118_ref0000016
scaffold3_143_ref0000121_ref0000095
scaffold3_144_ref0000122_ref0000188
scaffold3_146_ref0000124_ref0000162
scaffold3_150_ref0000136_ref0000014
scaffold3_151_ref0000098_ref0000065
scaffold3_152_ref0000097_ref0000182
scaffold3_159_ref0000090_ref0000151
scaffold3_161_ref0000066_ref0000098
scaffold3_170_ref0000030_ref0000054
scaffold3_172_ref0000034_ref0000171
scaffold3_186_ref0000027_ref0000076
scaffold3_190_ref0000184_ref0000152
scaffold3_196_ref0000186_ref0000175
0.5 Mb
1 Mb
1.5 Mb
2 Mb
2.5 Mb
500 kb
1000 kb

Endotrypanum

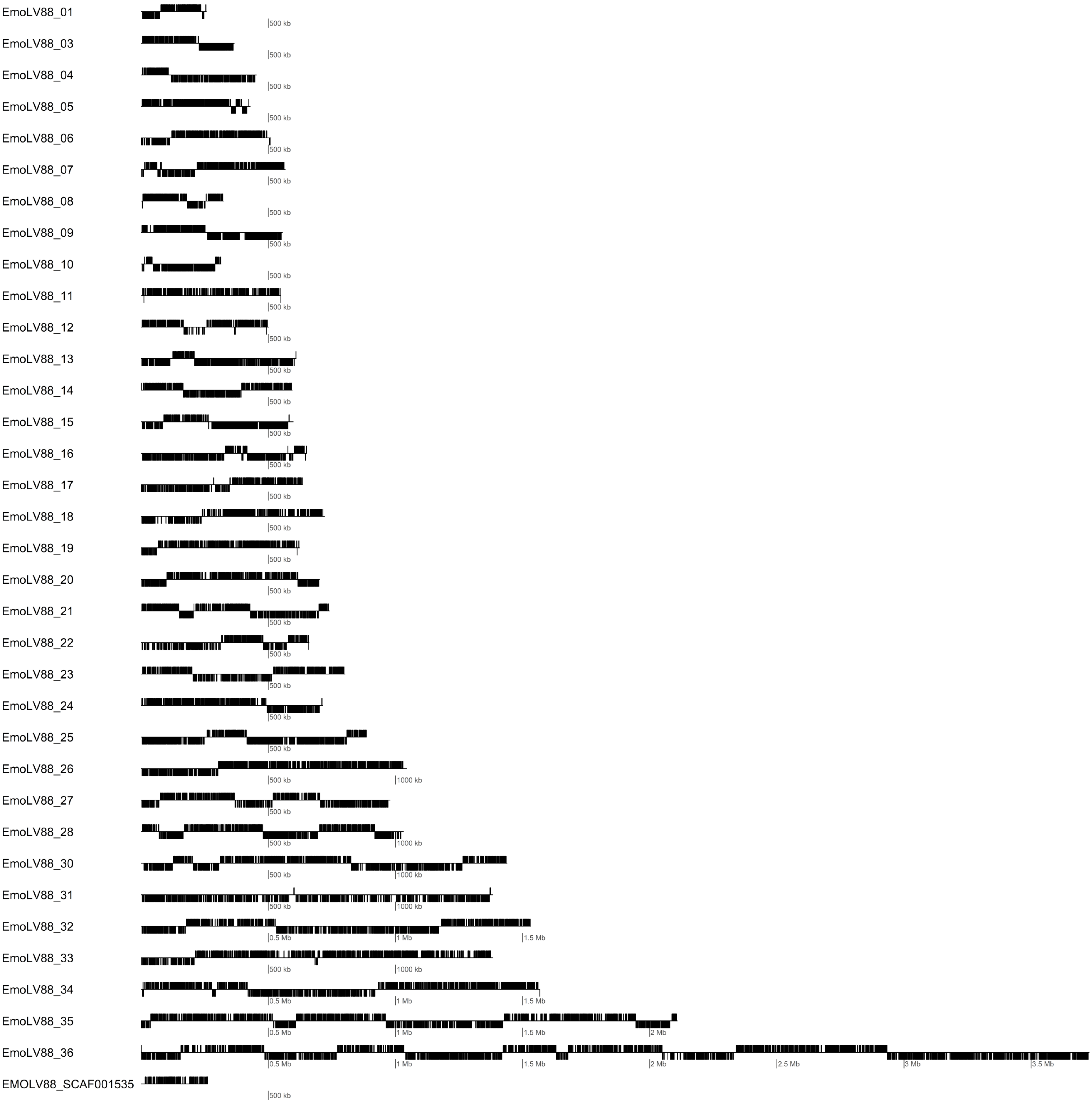

L. major

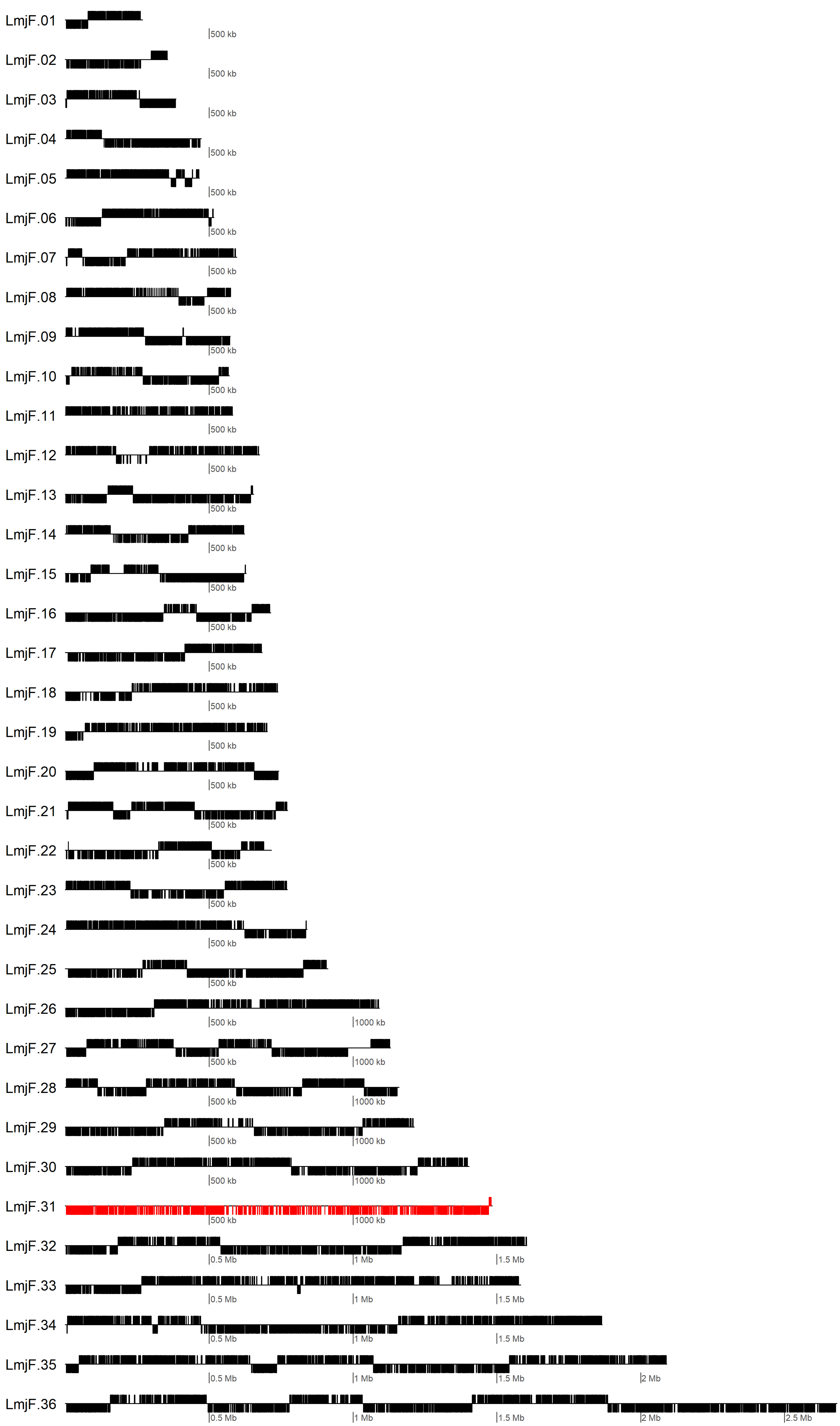
LmjF.01
500 kb
LmjF.02
500 kb
LmjF.03
500 kb
LmjF.04
500 kb
LmjF.05
500 kb
LmjF.06
500 kb
LmjF.07
500 kb
LmjF.08
500 kb
LmjF.09
500 kb
LmjF.10
500 kb
LmjF.11
500 kb
LmjF.12
500 kb
LmjF.13
500 kb
LmjF.14
500 kb
LmjF.15
500 kb
LmjF.16
500 kb
LmjF.17
500 kb
LmjF.18
500 kb
LmjF.19
500 kb
LmjF.20
500 kb
LmjF.21
500 kb
LmjF.22
500 kb
LmjF.23
500 kb
LmjF.24
500 kb
LmjF.25
500 kb
LmjF.26
500 kb
1000 kb
LmjF.27
500 kb
1000 kb
LmjF.28
500 kb
1000 kb
LmjF.29
500 kb
1000 kb
LmjF.30
500 kb
1000 kb
LmjF.31
500 kb
1000 kb
LmjF.32
0.5 Mb
1 Mb
1.5 Mb
LmjF.33
0.5 Mb
1 Mb
1.5 Mb
LmjF.34
0.5 Mb
1 Mb
1.5 Mb
LmjF.35
0.5 Mb
1 Mb
1.5 Mb
2 Mb
LmjF.36
0.5 Mb
1 Mb
1.5 Mb
2 Mb
2.5 Mb

L. donovani

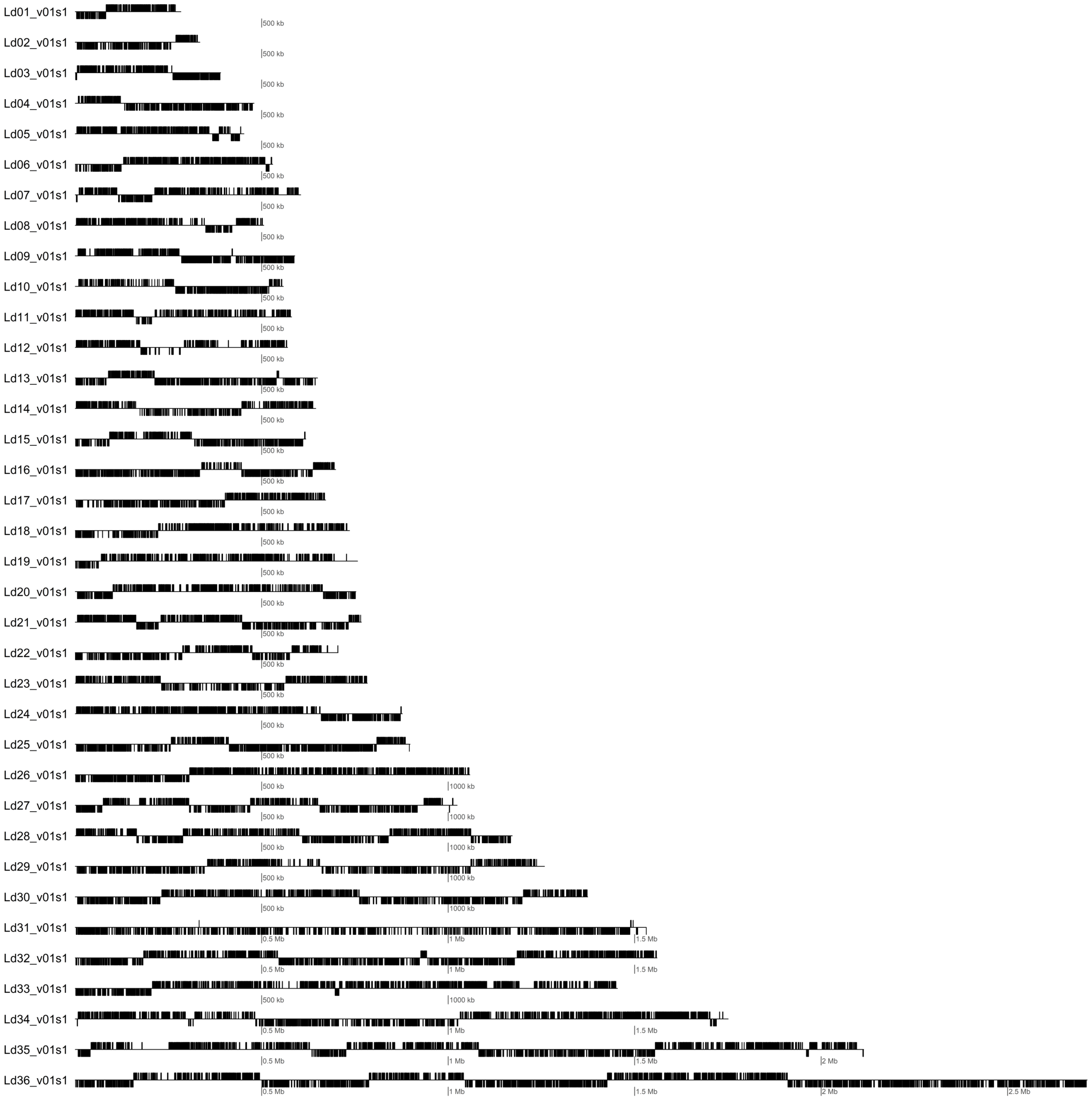
Ld01_v01s1
Ld02_v01s1
Ld03_v01s1
Ld04_v01s1
Ld05_v01s1
Ld06_v01s1
Ld07_v01s1
Ld08_v01s1
Ld09_v01s1
Ld10_v01s1
Ld11_v01s1
Ld12_v01s1
Ld13_v01s1
Ld14_v01s1
Ld15_v01s1
Ld16_v01s1
Ld17_v01s1
Ld18_v01s1
Ld19_v01s1
Ld20_v01s1
Ld21_v01s1
Ld22_v01s1
Ld23_v01s1
Ld24_v01s1
Ld25_v01s1
Ld26_v01s1
Ld27_v01s1
Ld28_v01s1
Ld29_v01s1
Ld30_v01s1
Ld31_v01s1
Ld32_v01s1
Ld33_v01s1
Ld34_v01s1
Ld35_v01s1
Ld36_v01s1
500 kb
1000 kb
0.5 Mb
1 Mb
1.5 Mb
2 Mb
2.5 Mb

L. infantum

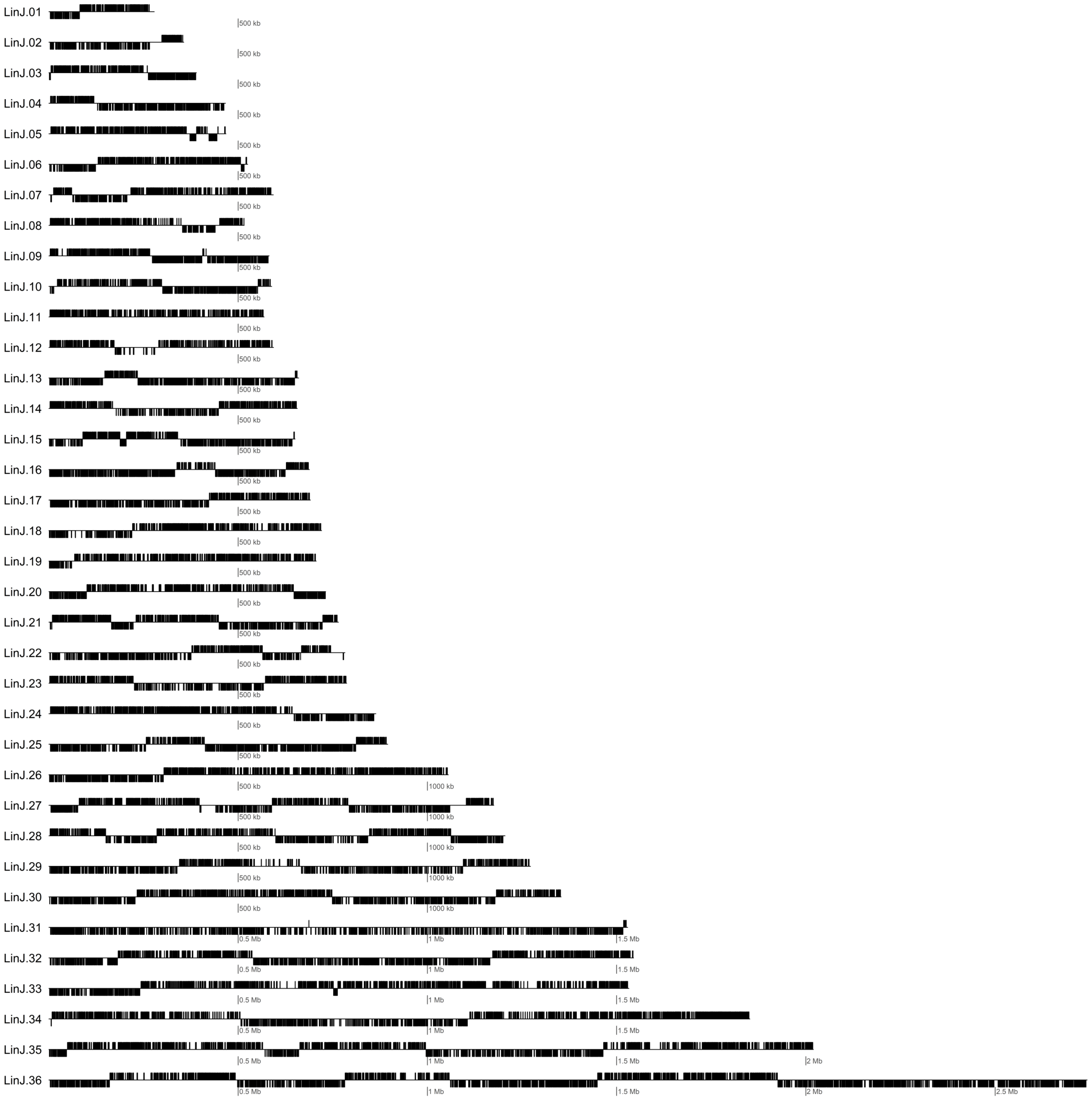
LinJ.01
LinJ.02
LinJ.03
LinJ.04
LinJ.05
LinJ.06
LinJ.07
LinJ.08
LinJ.09
LinJ.10
LinJ.11
LinJ.12
LinJ.13
LinJ.14
LinJ.15
LinJ.16
LinJ.17
LinJ.18
LinJ.19
LinJ.20
LinJ.21
LinJ.22
LinJ.23
LinJ.24
LinJ.25
LinJ.26
LinJ.27
LinJ.28
LinJ.29
LinJ.30
LinJ.31
LinJ.32
LinJ.33
LinJ.34
LinJ.35
LinJ.36
500 kb
1000 kb
0.5 Mb
1 Mb
1.5 Mb
2 Mb
2.5 Mb

Leptomonas

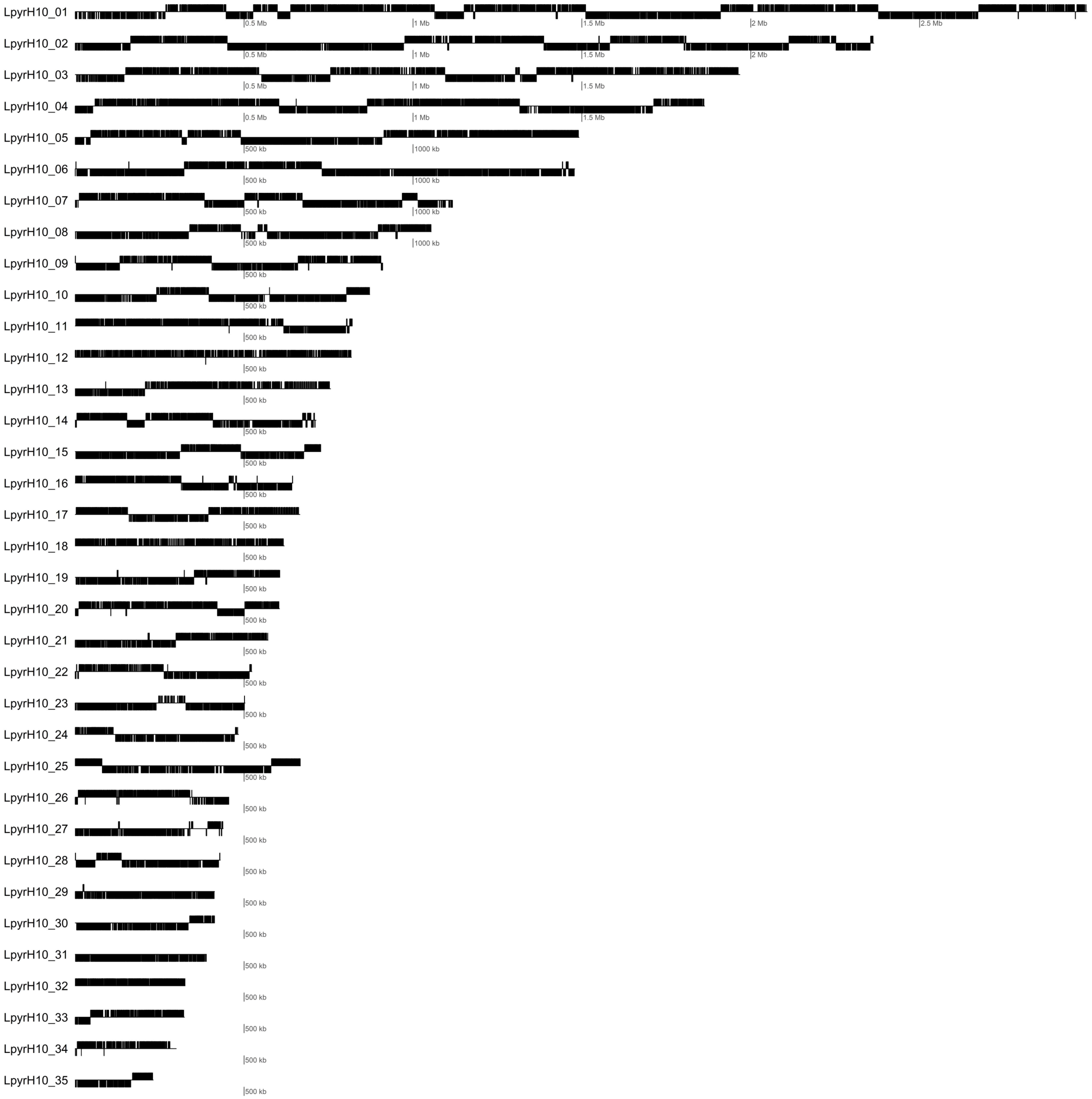
LpyrH10_01
LpyrH10_02
LpyrH10_03
LpyrH10_04
LpyrH10_05
LpyrH10_06
LpyrH10_07
LpyrH10_08
LpyrH10_09
LpyrH10_10
LpyrH10_11
LpyrH10_12
LpyrH10_13
LpyrH10_14
LpyrH10_15
LpyrH10_16
LpyrH10_17
LpyrH10_18
LpyrH10_19
LpyrH10_20
LpyrH10_21
LpyrH10_22
LpyrH10_23
LpyrH10_24
LpyrH10_25
LpyrH10_26
LpyrH10_27
LpyrH10_28
LpyrH10_29
LpyrH10_30
LpyrH10_31
LpyrH10_32
LpyrH10_33
LpyrH10_34
LpyrH10_35
0.5 Mb
1 Mb
1.5 Mb
2 Mb
2.5 Mb
500 kb
1000 kb

P. confusum

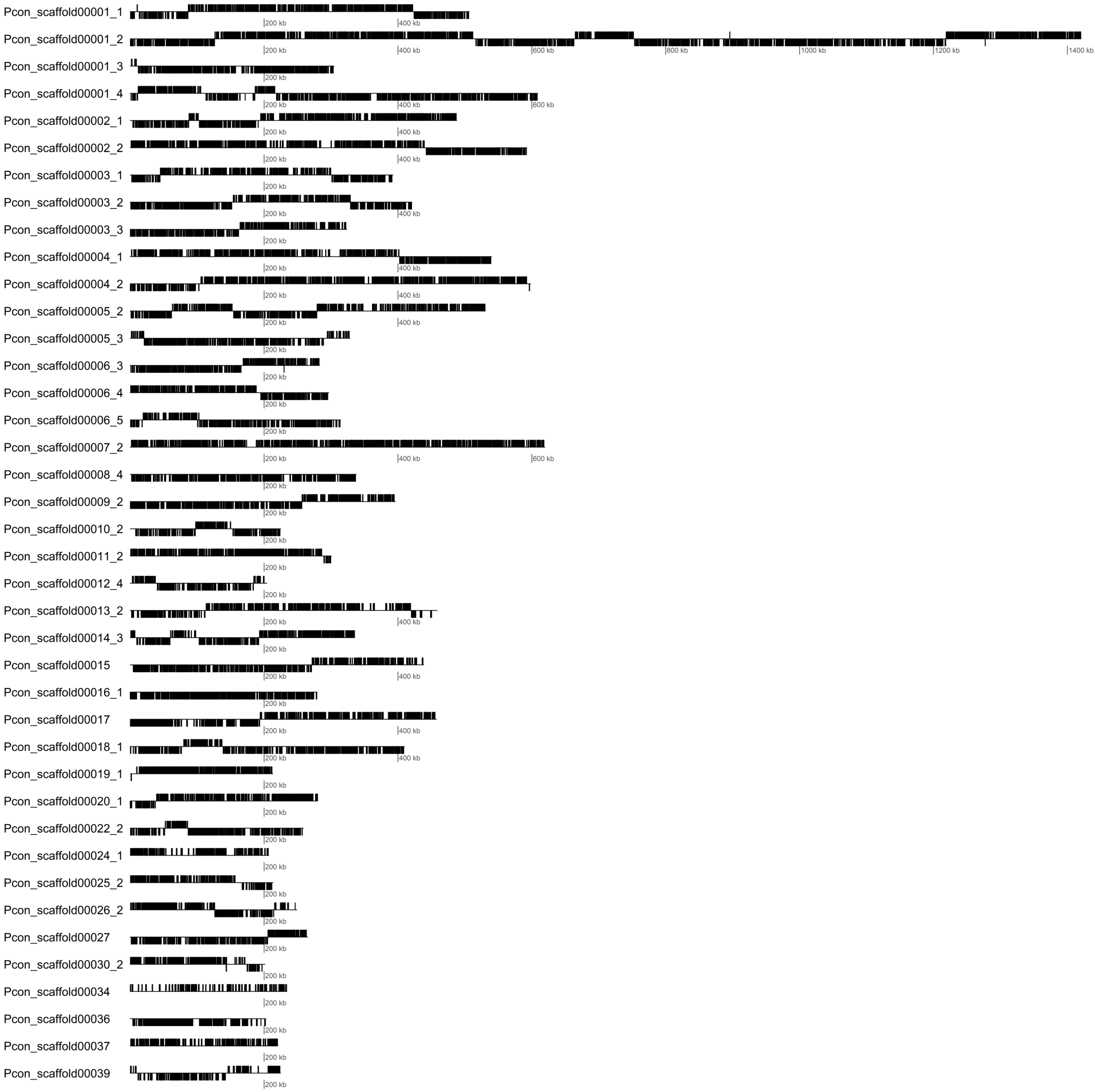
Pcon_scaffold00001_1
Pcon_scaffold00001_2
Pcon_scaffold00001_3
Pcon_scaffold00001_4
Pcon_scaffold00002_1
Pcon_scaffold00002_2
Pcon_scaffold00003_1
Pcon_scaffold00003_2
Pcon_scaffold00003_3
Pcon_scaffold00004_1
Pcon_scaffold00004_2
Pcon_scaffold00005_2
Pcon_scaffold00005_3
Pcon_scaffold00006_3
Pcon_scaffold00006_4
Pcon_scaffold00006_5
Pcon_scaffold00007_2
Pcon_scaffold00008_4
Pcon_scaffold00009_2
Pcon_scaffold00010_2
Pcon_scaffold00011_2
Pcon_scaffold00012_4
Pcon_scaffold00013_2
Pcon_scaffold00014_3
Pcon_scaffold00015
Pcon_scaffold00016_1
Pcon_scaffold00017
Pcon_scaffold00018_1
Pcon_scaffold00019_1
Pcon_scaffold00020_1
Pcon_scaffold00022_2
Pcon_scaffold00024_1
Pcon_scaffold00025_2
Pcon_scaffold00026_2
Pcon_scaffold00027
Pcon_scaffold00030_2
Pcon_scaffold00034
Pcon_scaffold00036
Pcon_scaffold00037
Pcon_scaffold00039
200 kb
400 kb
600 kb
800 kb
1000 kb
1200 kb
1400 kb

Porcisia

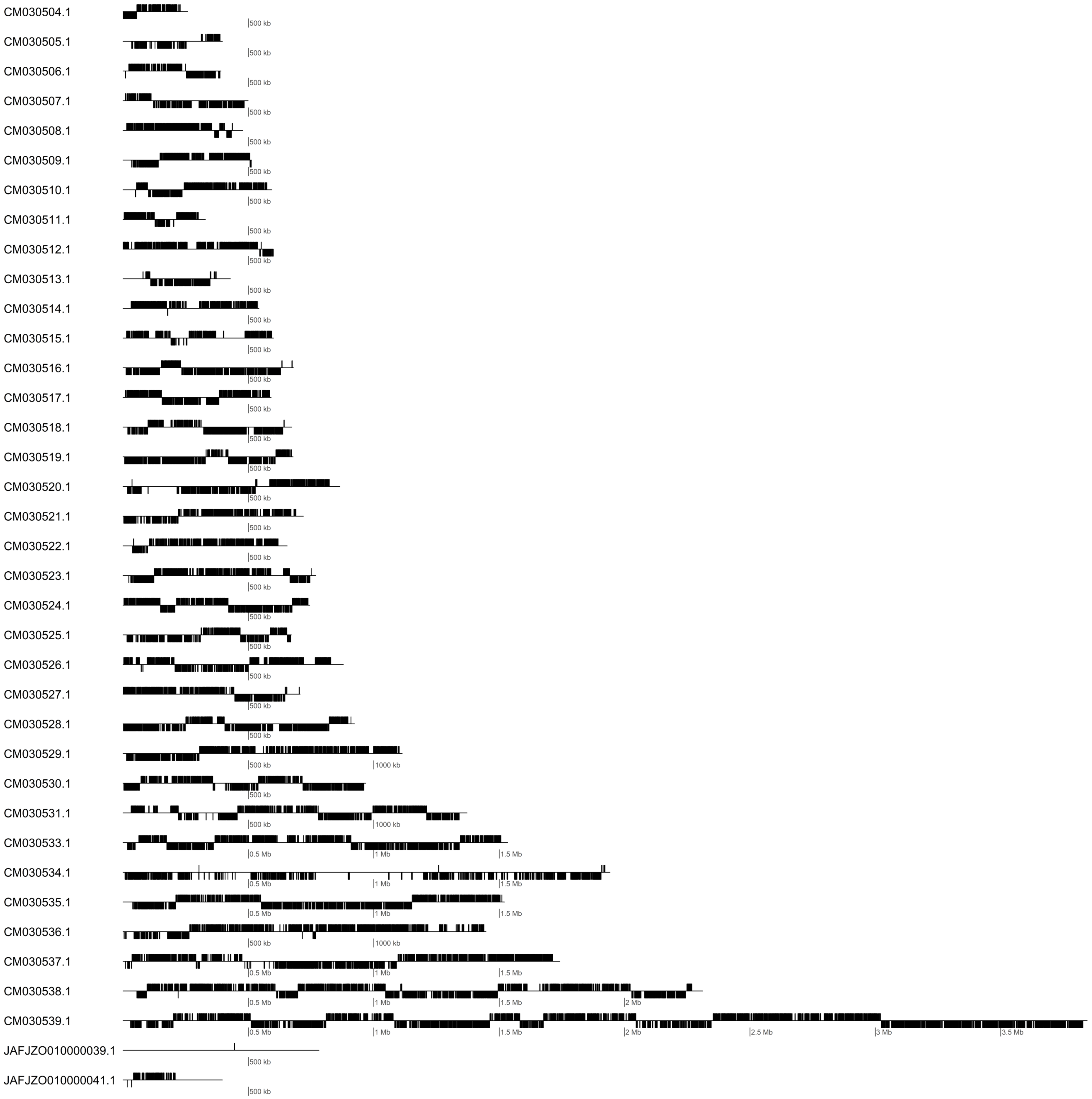
CM030504.1
CM030505.1
CM030506.1
CM030507.1
CM030508.1
CM030509.1
CM030510.1
CM030511.1
CM030512.1
CM030513.1
CM030514.1
CM030515.1
CM030516.1
CM030517.1
CM030518.1
CM030519.1
CM030520.1
CM030521.1
CM030522.1
CM030523.1
CM030524.1
CM030525.1
CM030526.1
CM030527.1
CM030528.1
CM030529.1
CM030530.1
CM030531.1
CM030533.1
CM030534.1
CM030535.1
CM030536.1
CM030537.1
CM030538.1
CM030539.1
JAFJZO010000039.1
JAFJZO010000041.1
500 kb
1000 kb
0.5 Mb
1 Mb
1.5 Mb
2 Mb
2.5 Mb
3 Mb
3.5 Mb

### T. brucei

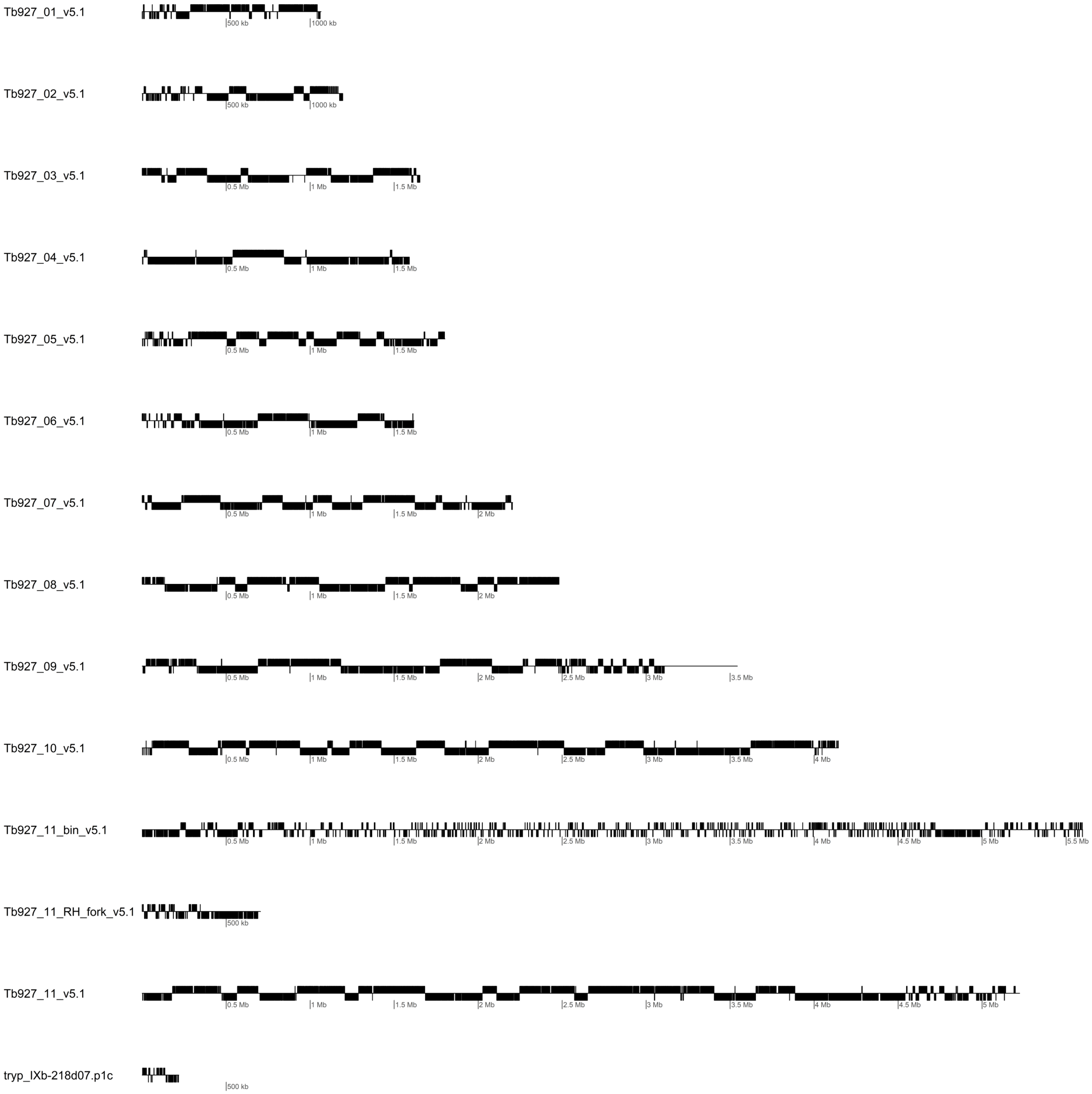
Tb927_01_v5.1
500 kb
1000 kb
Tb927_02_v5.1
500 kb
1000 kb
Tb927_03_v5.1
0.5 Mb
1 Mb
1.5 Mb
Tb927_04_v5.1
0.5 Mb
1 Mb
1.5 Mb
Tb927_05_v5.1
0.5 Mb
1 Mb
1.5 Mb
Tb927_06_v5.1
0.5 Mb
1 Mb
1.5 Mb
Tb927_07_v5.1
0.5 Mb
1 Mb
1.5 Mb
2 Mb
Tb927_08_v5.1
0.5 Mb
1 Mb
1.5 Mb
2 Mb
Tb927_09_v5.1
0.5 Mb
1 Mb
1.5 Mb
2 Mb
2.5 Mb
3 Mb
3.5 Mb
Tb927_10_v5.1
0.5 Mb
1 Mb
1.5 Mb
2 Mb
2.5 Mb
3 Mb
3.5 Mb
4 Mb
Tb927_11_bin_v5.1
0.5 Mb
1 Mb
1.5 Mb
2 Mb
2.5 Mb
3 Mb
3.5 Mb
4 Mb
4.5 Mb
5 Mb
5.5 Mb
Tb927_11_RH_fork_v5.1
500 kb
Tb927_11_v5.1
0.5 Mb
1 Mb
1.5 Mb
2 Mb
2.5 Mb
3 Mb
3.5 Mb
4 Mb
4.5 Mb
5 Mb
tryp_IXb-218d07.p1c
500 kb

T. cruzi

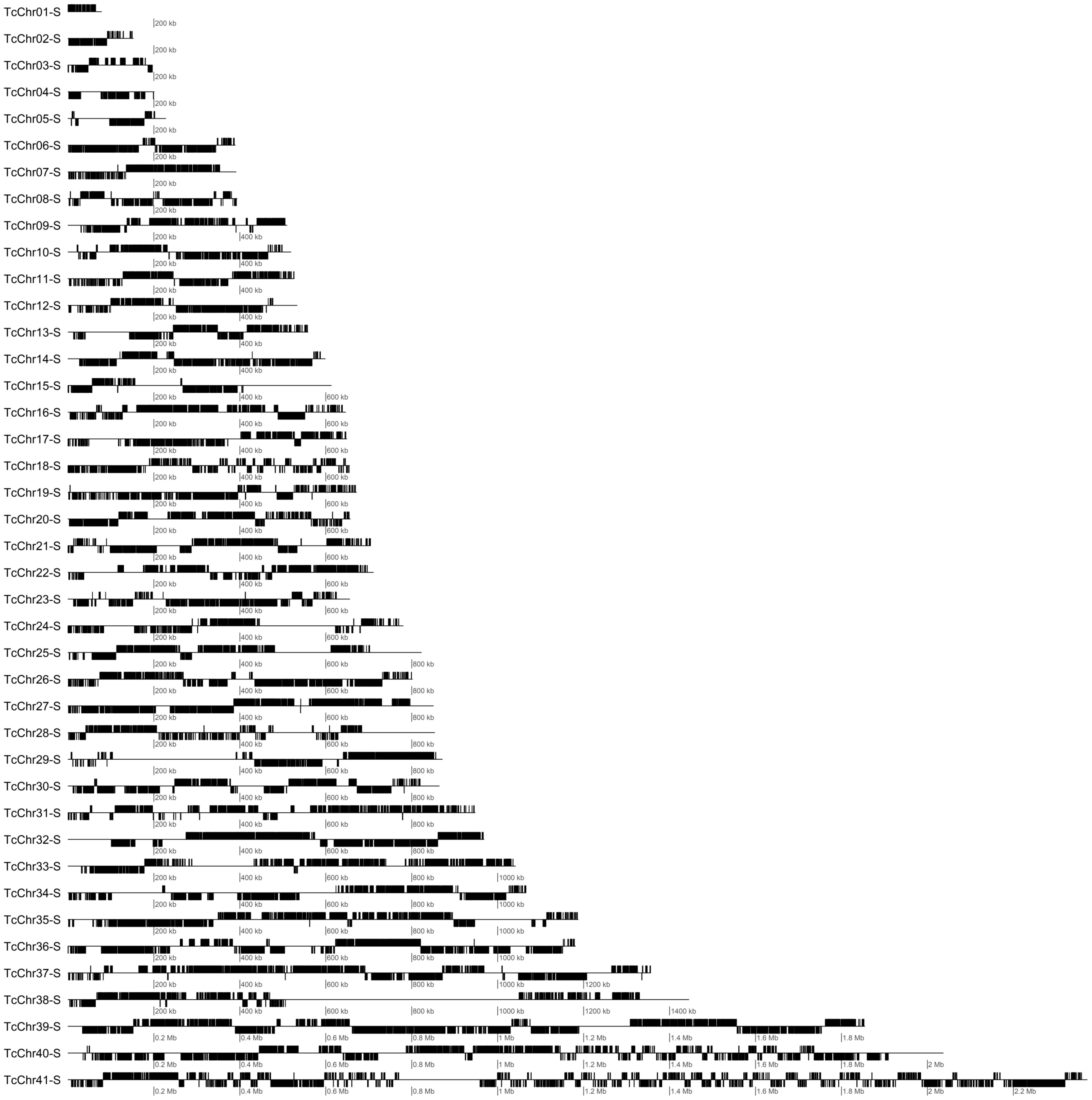
TcChr01-S
TcChr02-S
TcChr03-S
TcChr04-S
TcChr05-S
TcChr06-S
TcChr07-S
TcChr08-S
TcChr09-S
TcChr10-S
TcChr11-S
TcChr12-S
TcChr13-S
TcChr14-S
TcChr15-S
TcChr16-S
TcChr17-S
TcChr18-S
TcChr19-S
TcChr20-S
TcChr21-S
TcChr22-S
TcChr23-S
TcChr24-S
TcChr25-S
TcChr26-S
TcChr27-S
TcChr28-S
TcChr29-S
TcChr30-S
TcChr31-S
TcChr32-S
TcChr33-S
TcChr34-S
TcChr35-S
TcChr36-S
TcChr37-S
TcChr38-S
TcChr39-S
TcChr40-S
TcChr41-S
200 kb
400 kb
600 kb
800 kb
1000 kb
1200 kb
1400 kb
0.2 Mb
0.4 Mb
0.6 Mb
0.8 Mb
1 Mb
1.2 Mb
1.4 Mb
1.6 Mb
1.8 Mb
2 Mb
2.2 Mb

T. vivax

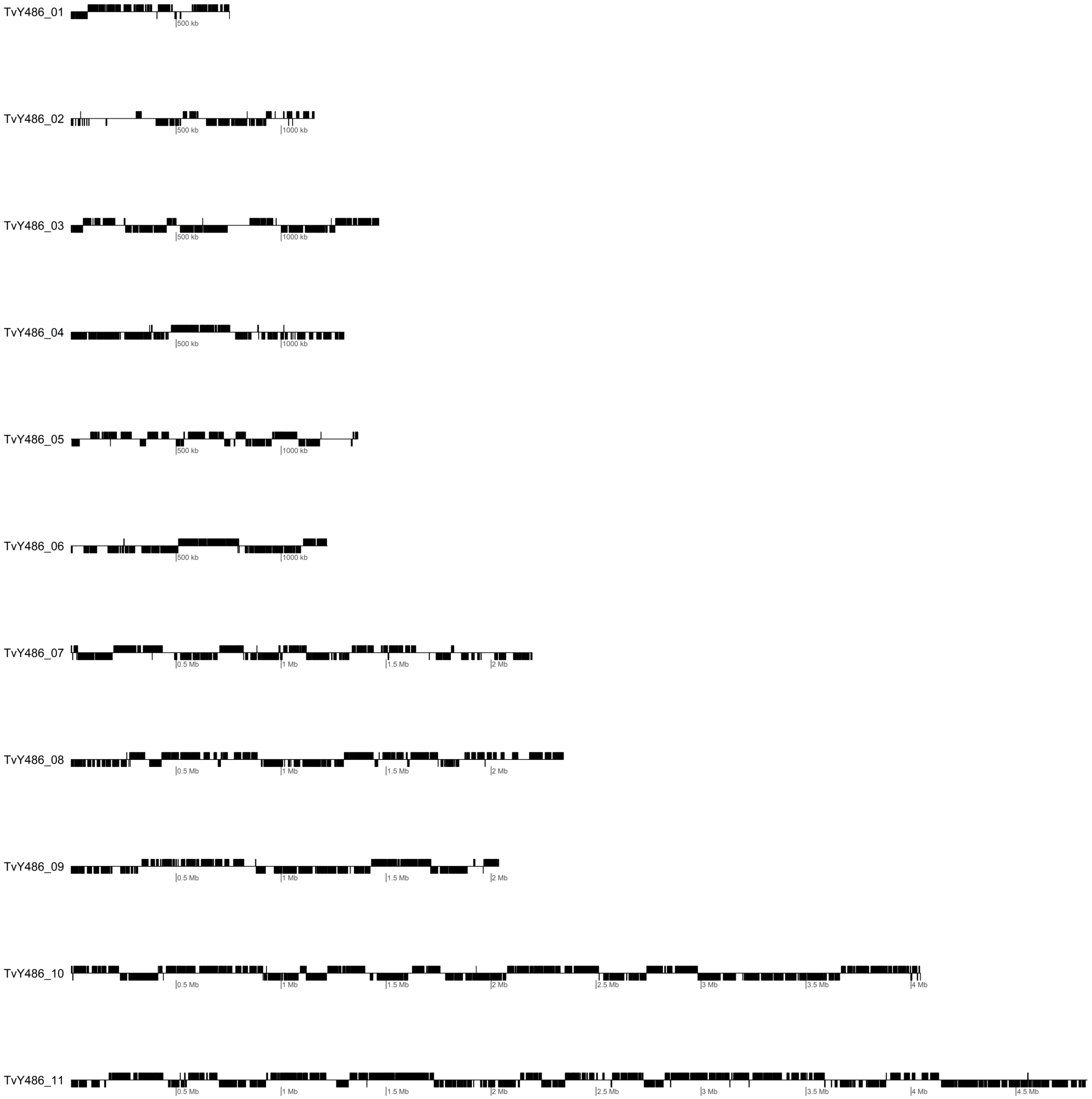
TvY486_01
500 kb
TvY486_02
500 kb
1000 kb
TvY486_03
500 kb
1000 kb
TvY486_04
500 kb
1000 kb
TvY486_05
500 kb
1000 kb
TvY486_06
500 kb
1000 kb
TvY486_07
0.5 Mb
1 Mb
1.5 Mb
2 Mb
TvY486_08
0.5 Mb
1 Mb
1.5 Mb
2 Mb
TvY486_09
0.5 Mb
1 Mb
1.5 Mb
2 Mb
TvY486_10
0.5 Mb
1 Mb
1.5 Mb
2 Mb
2.5 Mb
3 Mb
3.5 Mb
4 Mb
TvY486_11
0.5 Mb
1 Mb
1.5 Mb
2 Mb
2.5 Mb
3 Mb
3.5 Mb
4 Mb
4.5 Mb
