## Supplementary figure 5 for "Aneuploidies are an ancestral feature of trypanosomatids, and an ancient chromosome duplication is maintained in extant species"

**Supplementary Figure 5:** Codon usage in *L. donovani* genes in each chr. Comparison of Measure Independent of Length and Composition (MILC) in Chr 31 (red) and all other chromosomes combined (black), represented as **A)** density plots, **B)** boxplot and violin plots. **C)** Comparison of MILC (Y axis) and gene length (X axis), **D)** Log10 MILC in each chromosome. **E)** Heatmap of the codon relative presence in genes from each chromosome.

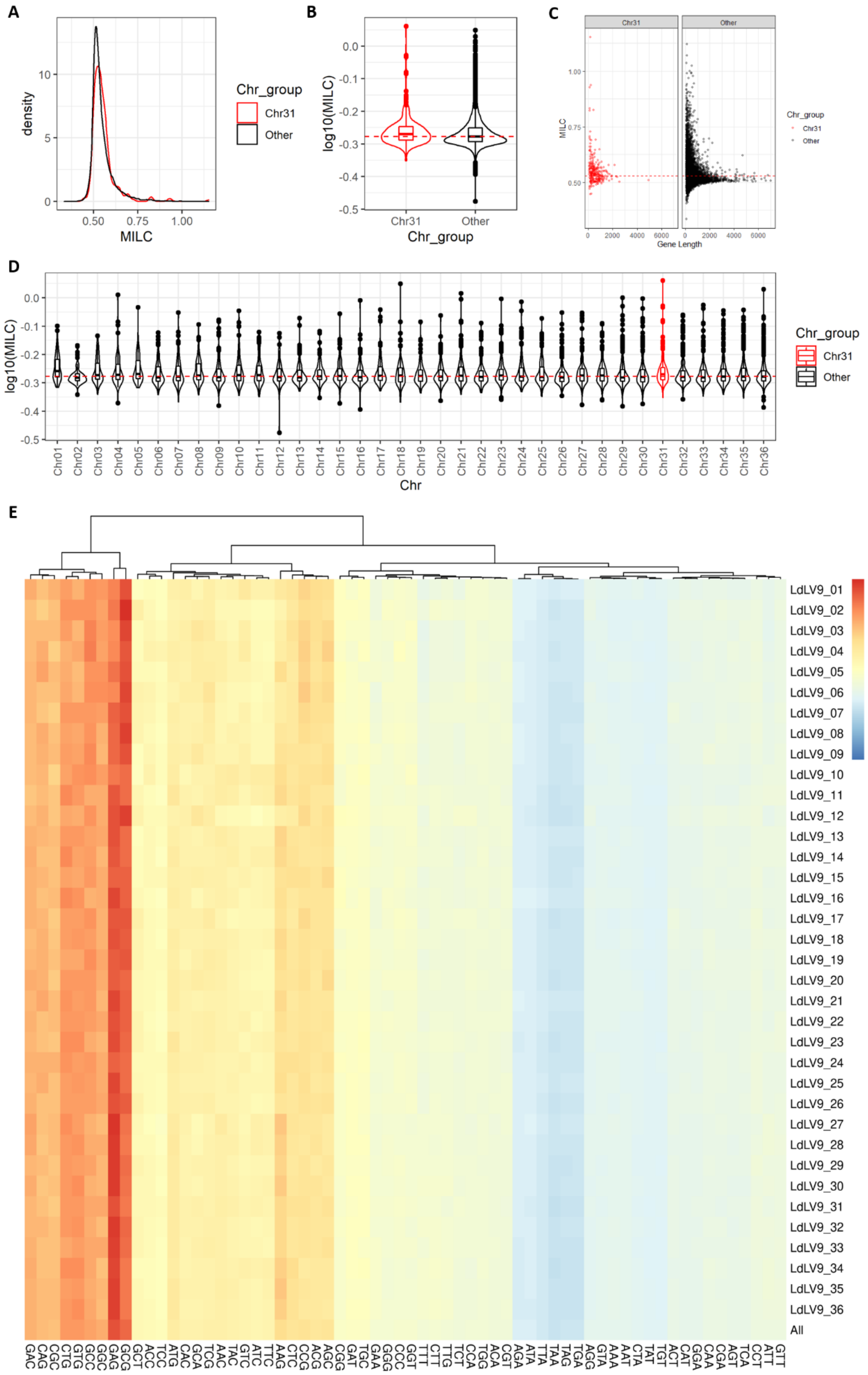
