## Supplementary figures and images for "Aneuploidies are an ancestral feature of trypanosomatids, and an ancient chromosome duplication is maintained in extant species"

### Supplementary figure 6

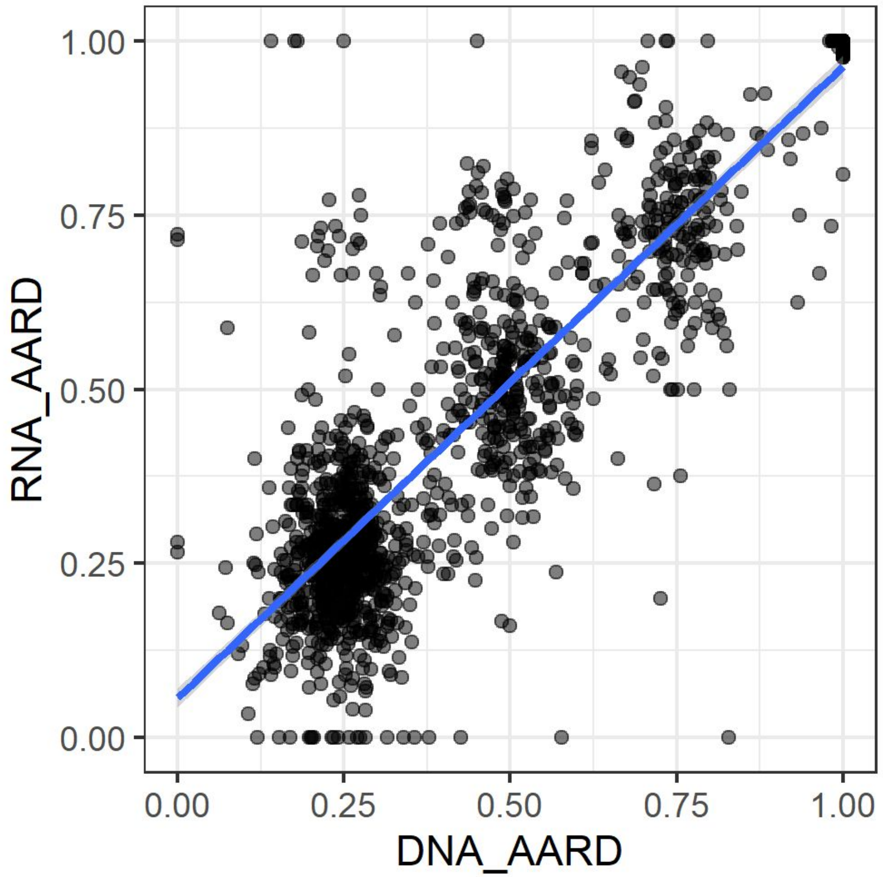

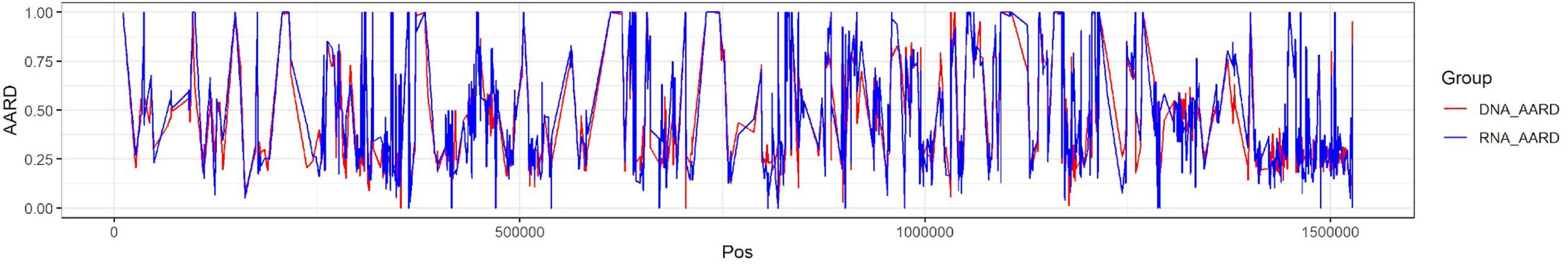

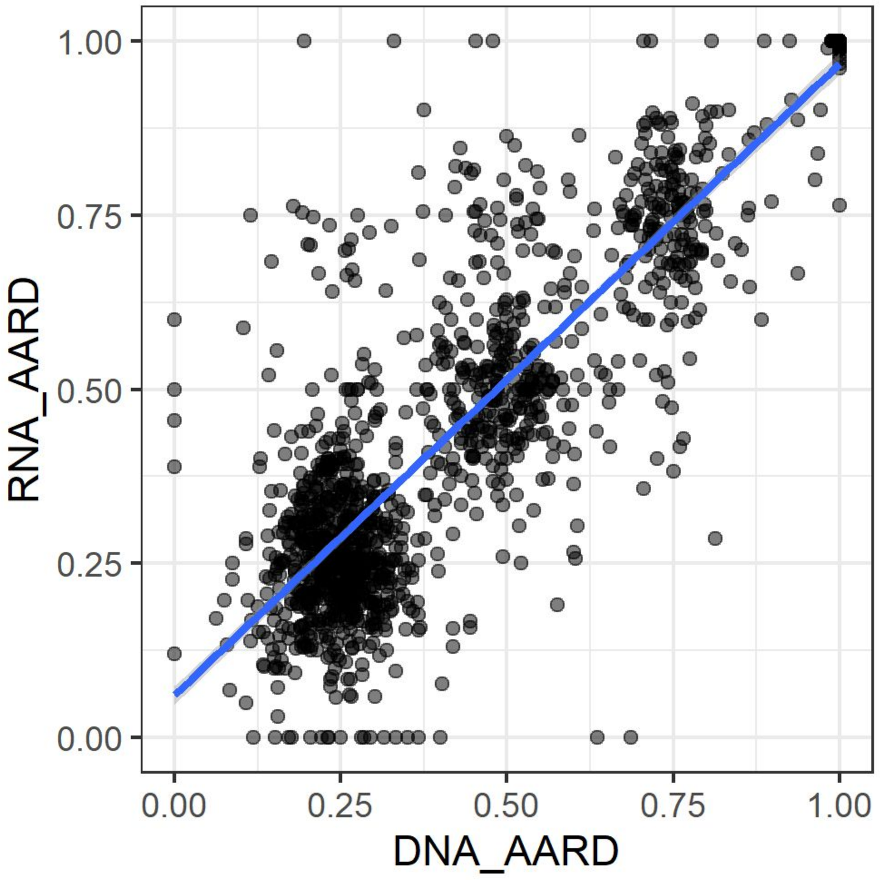

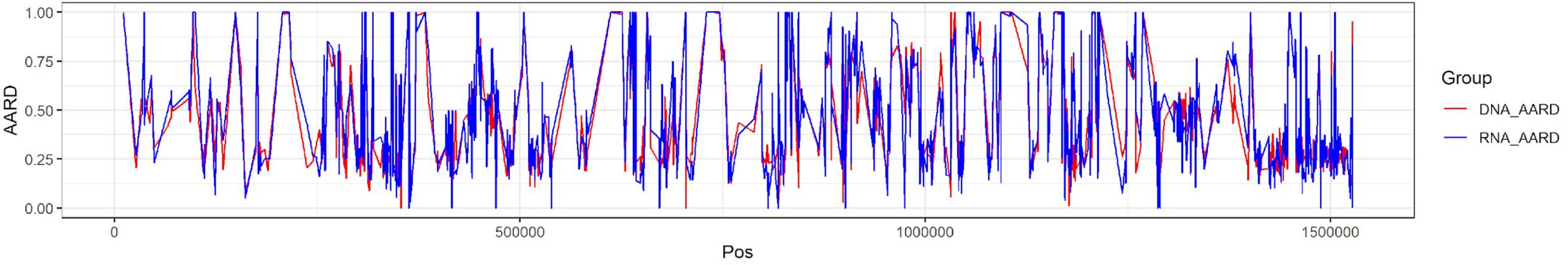

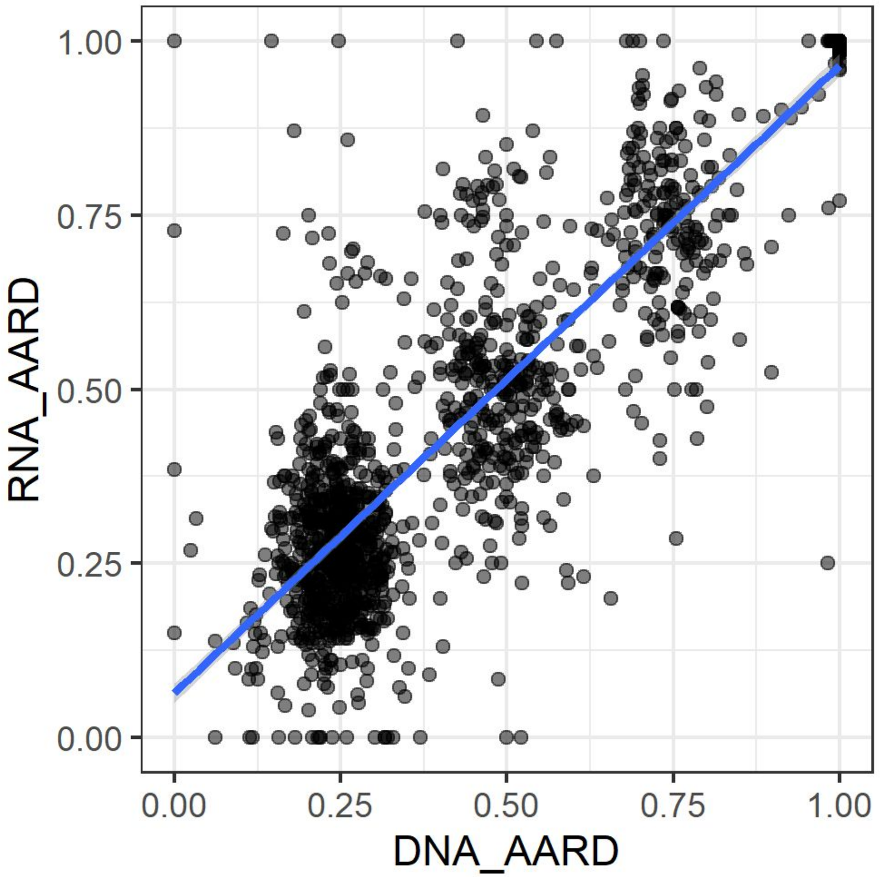

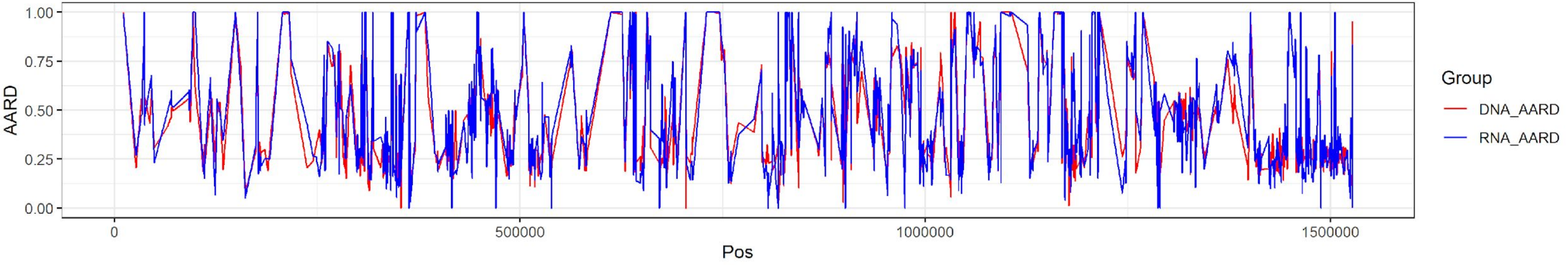

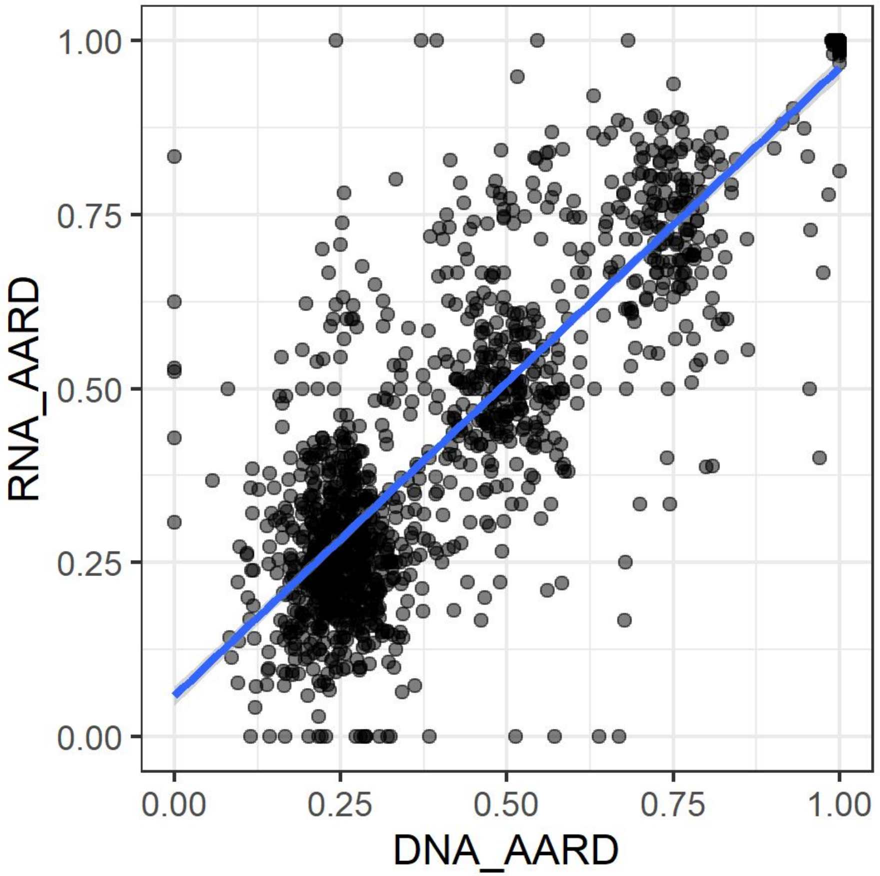

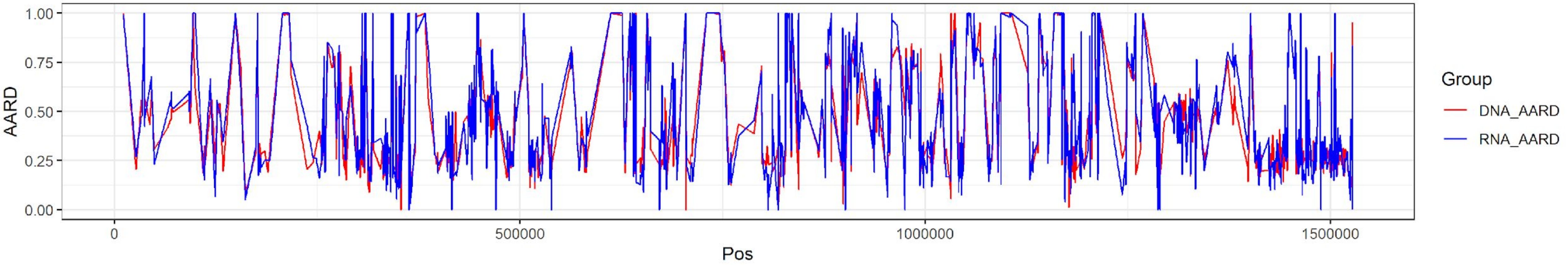

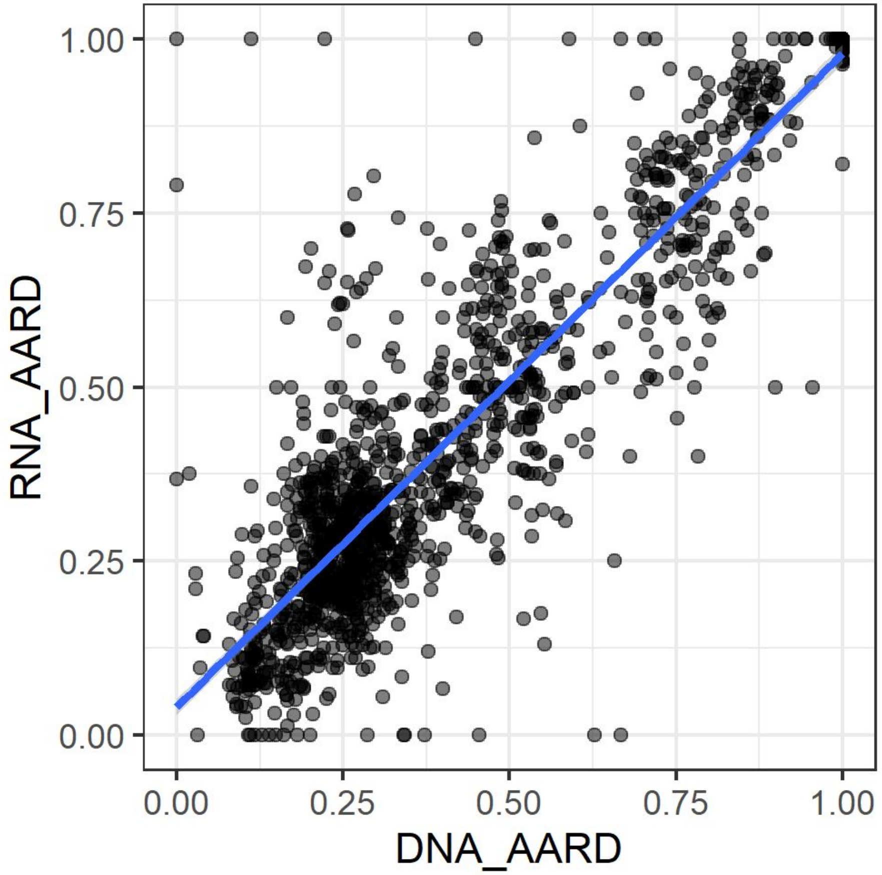

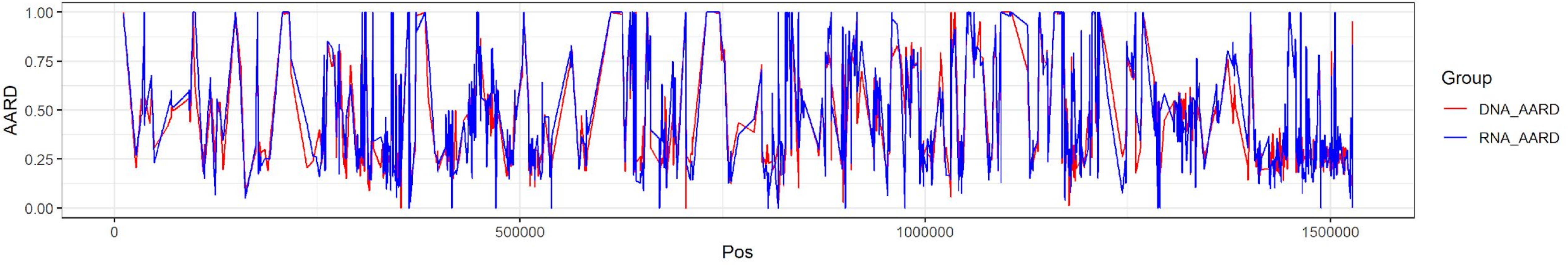

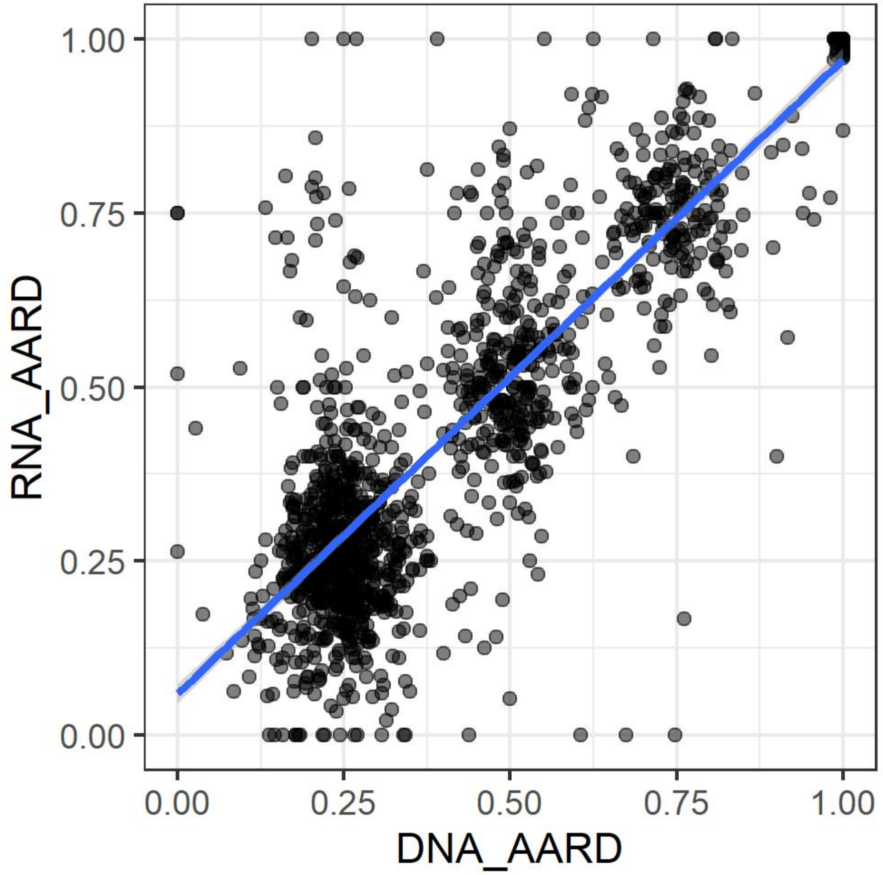

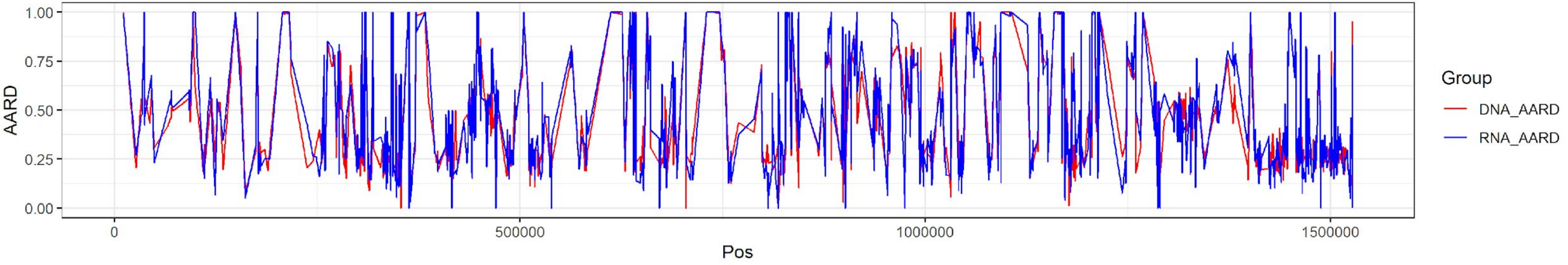

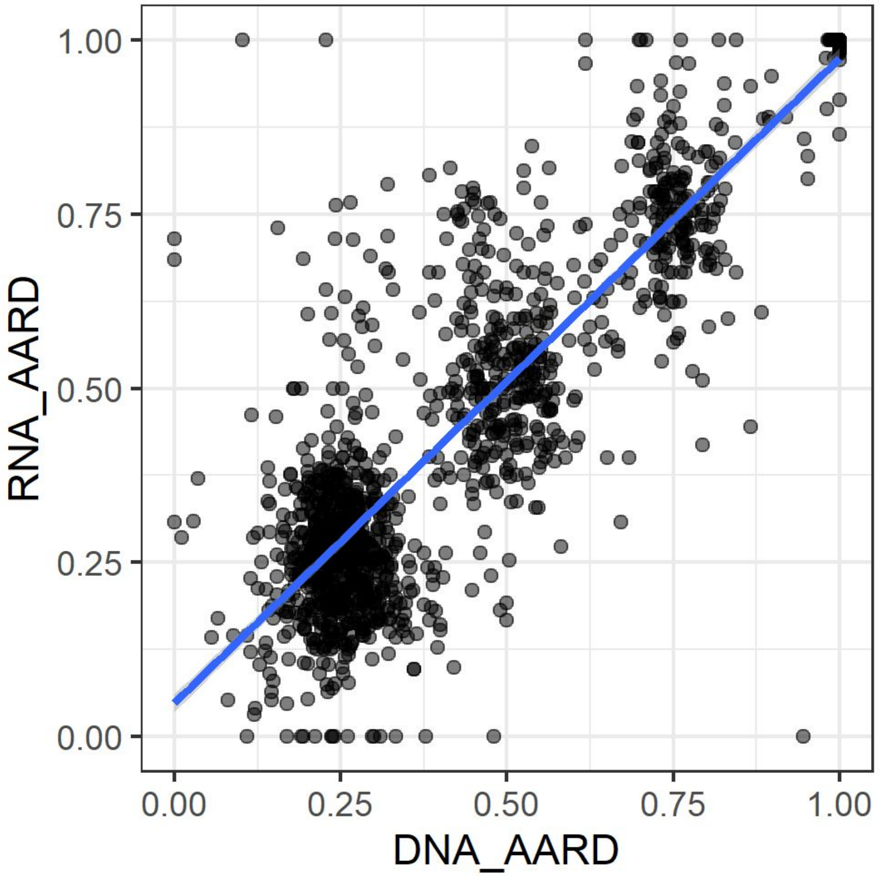

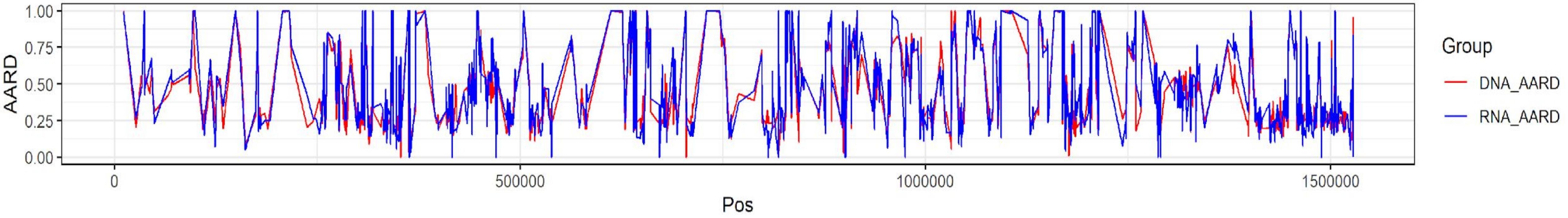
